# Systematic Characterization of Double Emulsion Droplets for Biological Applications

**DOI:** 10.1101/2022.03.04.483054

**Authors:** Suzanne G. K. Calhoun, Kara K Brower, Vineeth Chandran Suja, Gaeun Kim, Ningning Wang, Alexandra L. McCully, Halim Kusumaatmaja, Gerald G. Fuller, Polly M. Fordyce

## Abstract

Double emulsion droplets (DEs) are water/oil/water droplets that can be sorted via Fluorescence-Activated Cell Sorting (FACS), allowing for new opportunities in high-throughput cellular analysis, enzymatic screening, and synthetic biology. These applications require stable, uniform droplets with predictable microreactor volumes. However, predicting DE droplet size, shell thickness, and stability as a function of flow rate has remained challenging for monodisperse single core droplets and those containing biologically-relevant buffers, which influence bulk and interfacial properties. As a result, developing novel DE-based bioassays has typically required extensive initial optimization of flow rates to find conditions that produce stable droplets of the desired size and shell thickness. To address this challenge, we conducted systematic size parameterization quantifying how differences in flow rates and buffer properties (viscosity and interfacial tension at water/oil interfaces) alter droplet size and stability, across 6 inner aqueous buffers used across applications such as cellular lysis, microbial growth, and drug delivery, quantifying the size and shell thickness of >22,000 droplets overall. We restricted our study to stable single core droplets generated in a 2-step dripping-dripping formation regime in a straightforward PDMS device. Using data from 138 unique conditions (flow rates and buffer composition), we also demonstrated that a recent physically-derived size law of Wang et al^1^ can accurately predict double emulsion shell thickness for >95% of observations. Finally, we validated the utility of this size law by using it to accurately predict droplet sizes for a novel bioassay that requires encapsulating growth media for bacteria in droplets. This work has the potential to enable new screening-based biological applications by simplifying novel DE bioassay development.

## Introduction

In the past ten years, droplet microfluidic techniques have enabled new biological assays at unprecedented scale with applications ranging from disease diagnosis^2–4^ to synthetic biology^5–9^ and single cell analysis.^10–14^ Double emulsion (DE) (W/O/W water/oil/water) droplets are of particular interest, as their unique architecture allows them to be used as biological microreactors in drug delivery and synthetic cell engineering and to be sorted according to fluorescence using common flow cytometer instruments.^15–17^ This ability to sort DEs in high-throughput unlocks new opportunities for cellular screening (*e.g.* live-cell secretomics, next-generation sequencing of rare populations, and multiomic single cell analysis).^12,18–20^

However, DE droplets are generally considered more challenging to produce than single emulsion (SE) droplets. Each DE droplet contains a double layer structure comprised of an inner aqueous core encapsulated within an outer oil droplet miscible in an aqueous outer buffer.^16,21,22^ DE droplet stability and shell thickness therefore require balancing multiple forces - inertial, capillary, viscous and shear forces that arise during droplet generation, reaction processing and flow cytometry-at two different interfaces.^1,23–25^ Further complicating assay development, even small changes in the composition of the inner aqueous buffer can considerably change physical fluid properties and alter this balance of forces.^1,26^

Altered DE droplet size and shell thickness (diameter of the inner core, oil shell, and droplet overall) can, in turn, dramatically impact downstream biological assays.^14,16,27,28^ Across all applications, even small differences in inner aqueous core volume can lead to large changes in the diffusion kinetics and surface area-to-volume ratio of this compartment.^4,29,30^ These changes can modify effective soluble concentrations of proteins, enzymes, and reagents within the inner aqueous core, complicating assay standardization. In cell encapsulation assays, changes to the inner core diameter can also alter the loading distribution of cells within droplets.^16,31,32^

Beyond the inner aqueous core, efficient screening, sorting, and recovery of DE droplets via Fluorescence-Activated Cell Sorting (FACS) depends on both overall droplet volume and the thickness and deformability of the oil shell.^16,33,34^ Multicore and multilayer droplets have garnered much interest and study in their potential to assist fabrication of designer particles, but these droplets do not function well in FACS.^17,35–39^ Single core, uniform DE droplets are key for successful, clog-free FACS.^4,16,30,33,40–42^ Recent computational work suggests that high deformability, imparted by specific surfactants and thin oil shells, facilitates robust FACS by allowing droplets to flex in response to strong shear forces in the nozzle.^15,33,43,44^ Thin shells also hinder coalescence by increasing hydrodynamic resistance, which decreases core mobility and causes very thin shells to act as a lubricant.^15,43,45^

DE droplet generation methods and formation dynamics play critical roles in determining droplet stability and size. Generation via microfluidic devices^46,47^ provides improved control of size and monodispersity compared with traditional methods like bulk emulsification,^48^ along with greater design flexibility^36,38,39^ and operation control.^17^ Devices for producing DEs typically contain either a series of coaxial capillaries,^38,46^ two T-junctions,^49^ two flow-focusing junctions,^27,50,51^ and combinations thereof.^52,53^ In most cases, DEs are either generated by either vortexing single emulsions or reinjecting them into a droplet generator to embed them within another droplet,^48,54,55^ or in a glass capillary device, which requires technical skill to align and to operate due to the one-step formation, with its necessity of balancing 3 flow rates at one pinch-off point as inner and outer droplets shear simultaneously.^1,46,48^ These approaches can increase polydispersity by a failure to encapsulate all single emulsions into double emulsions.^17,46,48,56^ As an alternate approach, PDMS devices that generate DEs via a two-step flow regime aligned on a single device are straightforward to operate and create monodisperse droplets ideal for FACS.

A more systematic understanding of how DE droplet size, shell thickness and stability vary with flow rates and solution composition has the potential to greatly simplify initial assay development. Several prior experimental DE size and shell thickness characterization studies in a one-step formation regime have explored the effect of flow rates^35,49,57,58^ and device geometry parameters.^59^ Despite the fact that influences on size likely differ from one-step to two-step formation regimes (those with two distinct pinch-off points for the inner and outer droplets),^1^ few size studies address two-step formation, with existing studies only addressing transitions from one-step formation to two-step^27,60^ and flow rates in T junctions.^49^ Two-step formation, as in our linear dual flow focuser device geometry here, is straightforward to operate with only two flow rates to balance simultaneously (at flow focuser 1 (FF1)), and by precluding several unstable regimes due to a flow resistor in the device geometry. In particular, many studies focus on multicore or 3+ layer emulsions,^17,36,46^ and there is a distinct lack of DE size characterization focusing on single core droplets, important for FACS sorting and thus for many biological applications. Numerical studies, predominantly using one-step formation, have addressed the importance of flow rates, interfacial tension, geometry,^61,62^ viscosity,^63^ and flow regime prediction,^59^ and have formulated size scaling laws.^46,64^ Nevertheless, few systematic experimental studies have been conducted to validate these proposed size laws and stability regimes. In addition, size characterization in DE droplets has been limited to idealized flow solutions (*e.g.* PEG-dextran or PBS), limiting translation to biological assays. As a result, developing a novel DE assay for biological applications typically requires an initial exhaustive empirical search for appropriate flow rates and surfactants, requiring time and resources and creating obstacles for end users.

We previously developed an open-source device design (Dropception) and workflow for robustly generating and sorting DE droplets via FACS.^16^ Here, we build on this work with a systematic scan of DE droplet size variation across a wide range of different buffer solutions and flow rates, with a particular focus on varying the inner aqueous buffer, most likely to change between biological assays. By restricting our work to a stable dripping-dripping flow regime with matched periodicity in a two-step formation device, we focused our investigation exclusively on the single-core droplets essential to FACS. Overall, we quantified size and shell thickness for DE droplets containing 6 different inner aqueous buffers as a function of 138 combinations of aqueous inner and oil flow rates that yielded monodisperse single core DE droplet populations with <5% CV in size. In parallel, we characterized bulk and interfacial properties of each aqueous input solution to understand how interfacial tension and viscosity influence droplet size and stability. We applied a recent size scaling model of one-step formation by Wang et al1 to our two-step formation regime, with our broad systematic dataset acting as an empirical validation of this model for wide biological applications. To test the accuracy of the model, we predicted droplet flow rates for desired shell thickness with a novel inner aqueous buffer, generated droplets, and confirmed that measured droplet size matched predictions. We anticipate that this platform, model and data will prove broadly useful for life scientists seeking to develop new DE-based assays.

## Results and Discussion

### 1. Size Characterization Pipeline

To develop a generalizable and predictive model for DE droplet size and shell thickness as a function of flow rate, we: (1) generated stable single core DE droplets under varying flow rates and incorporating different buffers and surfactants, (2) quantified droplet size and shell thickness via microscopy, and (3) used these data to derive fit parameters for a universal size scaling law that predicts droplet size and shell thickness using only flow rates and surfactant properties as inputs (**Fig. 1**).

**Figure 1.**
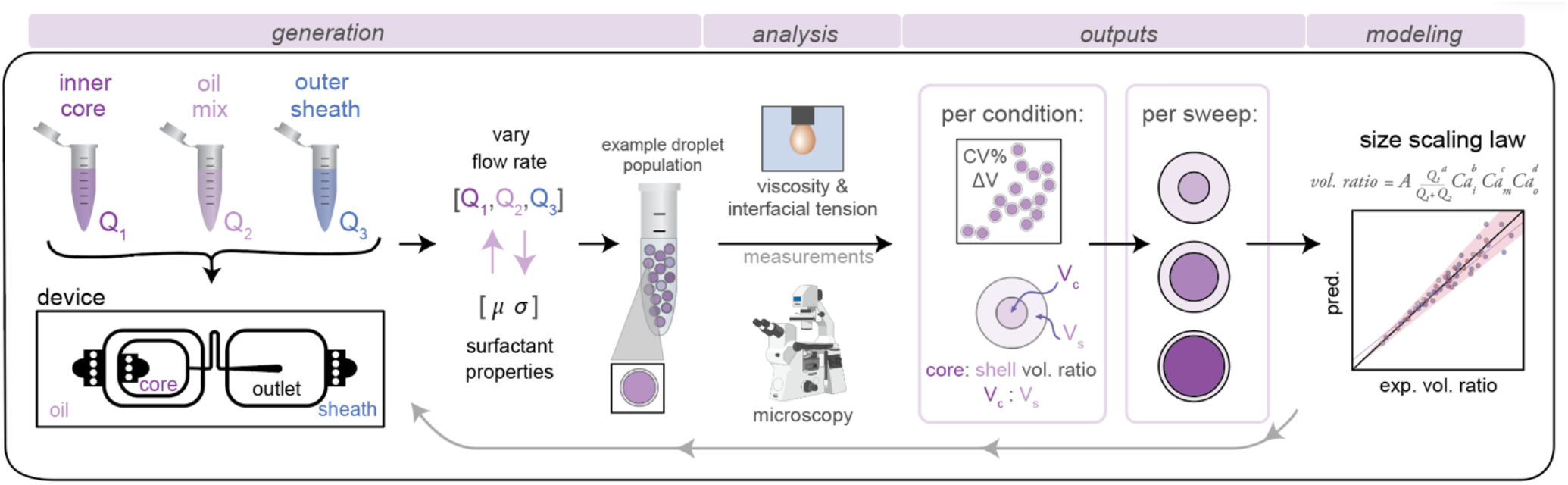
DE size characterization pipeline: (1) **generation** of W/O/W droplets from different inner aqueous buffers, oil mix, and outer sheath buffer across multiple flow rate conditions, (2) **analysis** to quantify droplet size and shell thickness via microscopy, with **outputs** of %CV and core:shell volume ratio of each condition, and viscosity and interfacial tension (IFT) via pendant drop tensiometry and cone and plate rheometry, and (3) **modeling** to derive a universal scaling law capable of predicting droplet shell thickness from flow rates and IFTs.

### 2. Double Emulsion Droplet Generation via Dripping-Dripping Regime

We generated DE droplets using an integrated single-layer microfluidic device (‘Dropception’) that forms both the core W/O droplet and final W/O/W droplet on the same chip.^16,31^ DE droplets are formed via a dual flow focuser geometry: single aqueous buffer-in-oil droplets are formed at the first flow focusing junction (FF1), and then these droplets are encapsulated within an oil shell via pinch-off of the W/O droplet in a fast-flowing outer aqueous sheath solution stream at the second flow focusing junction (FF2).

Between FF1 and FF2 (FF1: 22.5 μm wide x 22.5 μm tall, FF2: 50 μm x 50 μm), a serpentine flow resistor (22.5 μm wide x 22.5 μm tall) increases resistance to equally space W/O droplets prior to pinch-off (**Figs. 1, S1**). At FF2, the channel height increases to 50 μm, which reduces downstream resistance to create a dominant dripping-dripping regime for droplet formation (**Movie S1**) and prevent the jetting-dripping droplet formation previously seen in DE generation systems.^1,27^ In this scheme, FF1 and FF2 function as largely decoupled systems, allowing us to hold outer sheath flow rates (Q3) constant and generate DEs with a constant outer diameter for reproducible FACS sorting while varying inner aqueous (Q1) and oil (Q2) rates to alter inner droplet size. The device geometry of a flow resistor separating the two flow focusers effectively eliminates possibility of several unstable flow regimes, making stable droplets more easily achievable.

This device reduces size variability associated with the reinjection required for two-step droplet generation, is easy to fabricate, and requires only 3 syringe pumps for operation (<$10,000 total cost). While the Dropception device used here produces droplets of total diameter ^~^40 μm (sortable with FACS), it can be globally scaled to produce smaller or larger DEs.

### 3. Size Parameterization Flow Rate Sweep: Base Condition

Using this device, we designed a systematic size parameterization sweep to test how varying flow rates altered DE droplet size and shell thickness. Here we sequentially vary Q1 and Q2 (inner and oil flow rates) while holding Q3 constant to decrease number of variables. To systematically sample stable flow rate conditions, we varied either: (1) Q1 alone while keeping Q2 constant, (2) Q2 alone while keeping Q1 constant, or (3) both Q1 and Q2 simultaneously while keeping their total flow rate constant (Qt = Q1 + Q2) (**Fig. S2**).

To provide high-resolution information at flow rates most likely to be useful for biologists interested in sorting DEs, we first identified a central condition that generated stable single core droplets with a core:shell ratio of ^~^0.60 and then sampled the stable range in regularly spaced intervals in either direction. At edges of the sample ranges, single core droplets gave way to other behaviours (*e.g.* droplets with multiple cores, droplets with satellite oil drops, and droplets without cores) (**Fig. S2**). We define a stable flow rate condition as a dripping-dripping flow regime with matching periodicity of SE formation at FF1 with DE encapsulation at FF2 that produces droplets with >95% single cores as determined by microscopy; we considered any formation behaviour outside these restrictions an instability.

To begin, we tested DEs composed of solutions previously optimized for FACS^16^: (1) an inner aqueous buffer typically used for protein extraction, gentle cell lysis, and ddPCR droplet applications (1X PBS and 1% Tween-20, a polysorbate-family nonionic detergent), (2) an oil solution, consisting of HFE 7500 fluorinated oil with 2.2% Ionic Krytox FSH-157 surfactant, that is biocompatible and frequently used in cell growth and droplet PCR applications,^4,16^ and (3) an outer aqueous sheath solution comprised of 1X PBS with 2% Pluronix F68 and 1% Tween-20. For this droplet composition, a wide range of flow rates (Q1: 110-230 μl/hr, Q2: 280-760 μl/hr) produced highly monodisperse single core droplets with core volume: shell volume ratios (V_c_:V_s_) from 0.181-0.703 (**Figs. 2A,B; Table S5**). As expected, reducing the inner aqueous flow rate (Q1) linearly decreased core volume (**Fig. 2B**, middle), increasing the oil flow rate (Q2) linearly increased shell thickness (**Fig. 2B**, left), and changing Q1 and Q2 simultaneously while maintaining the same total inner flow rate (Q_t_) generated droplets with a constant outer diameter (**Fig. 2B**). Surprisingly, varying either Q1 or Q2 while keeping the other inner flow rate constant generated droplets with a largely constant outer diameter (mean: 39.4 μm) despite large changes in Q_t_. This is consistent with the device’s flow resistor shielding FF2 from most effects of FF1 variation at constant Q3 under the requirement of matched single emulsion and double emulsion encapsulation rates at FF1 and FF2. The dynamic range of the oil flow rate (Q2) that generated monodisperse droplets was wider than that for inner aqueous flow rate (Q1), consistent with the lower viscosity and interfacial tension (IFT) of the oil solution (global statistics, **Table 1**; fluid properties, **Table 2**; ranges, **Table S5**). Overall, the mean CVs for all measured outer diameters and inner core diameters were 2.16% and 1.51%, respectively, with all combinations under the 5% CV line, important for reaction success and large particle FACS, highly sensitive to clogs (**Fig. 2C**).^16,55,65^

**Table 1:** Global statistics for all conditions.

|  | PBS | PBS 1%<br>tween-20 | PBS 1%<br>tween-20<br>replicate | Np40 | Np40<br>replicate | PEG 10% | M9 | M9 glucose | Total |
| --- | --- | --- | --- | --- | --- | --- | --- | --- | --- |
| number of stable points | 14 | 19 | 23 | 31 | 21 | 26 | 32 | 30 | 196 |
| number of instabilities<br>imaged | 0 | 4 | 1 | 4 | 4 | 3 | 3 | 1 | 20 |
| number of instabilities<br>recorded | 8 | 16 | 10 | 11 | 10 | 6 | 6 | 7 | 74 |
| number of trials | 22 | 39 | 34 | 46 | 35 | 35 | 41 | 38 | 290 |
| number of droplets<br>analyzed | 1349 | 1679 | 2238 | 3157 | 2090 | 3309 | 3711 | 3399 | 20932 |
| resultant mean outer<br>diameter CV for stable<br>conditions | 0.014 | 0.021614 | 0.011 | 0.014889 | 0.020847 | 0.019 | 0.019321 | 0.018 | 0.017333875 |
| resultant mean inner<br>diameter CV for stable<br>conditions | 0.01829 | 0.015135 | 0.015822 | 0.012758 | 0.015036 | 0.012793 | 0.013209 | 0.018702 | 0.015218125 |

**Table 2:** Interfacial and bulk fluid parameters.

| Surfactant System | Phase | Components | Density (kg/m<sup>3</sup>) | Dynamic viscosity (mPa s) | Interfacial tension with Oil (mN/m) |
| --- | --- | --- | --- | --- | --- |
| PBS | Inner | PBS | 1005.58 | 0.931 | 0.543 |
|  | Middle | HFE-7500 + 2.2% ionic Krytox | 1619.72 | 1.613 | n/a |
|  | Outer | PBS + 1% Tween-20 + 2% Pluronic-F68 | 1007.97 | 1.303 | 0.318 |
| PBS + 1% Tween-20 | Inner | PBS + 1% Tween-20 | 1006.48 | 0.988 | 0.319 |
|  | Middle | HFE-7500 + 2.2% ionic Krytox | 1619.72 | 1.613 | n/a |
|  | Outer | PBS + 1% Tween-20 + 2% Pluronic-F68 | 1007.97 | 1.303 | 0.318 |
| PBS + 0.9% NP40 | Inner | PBS + 0.9% NP40 inner | 1006.07 | 1.003 | 1.41 |
|  | Middle | HFE-7500 + 2.2% ionic Krytox | 1619.72 | 1.613 | n/a |
|  | Outer | PBS + 1% Tween-20 + 2% Pluronic-F68 | 1007.97 | 1.303 | 0.318 |
| M9 bacterial media | Inner | M9 salts | 1013.0 | 0.861 | 12.84 |
|  | Middle | HFE-7500 + 2.2% ionic Krytox | 1619.72 | 1.613 | n/a |
|  | Outer | M9 salts + 2% Pluronic-F68 | 1013.4 | 1.412 | 0.5220 |
| M9 + 25mM glucose | Inner | M9 salts + 25mM glucose | 1017.5 | 0.967 | 11.600 |
|  | Middle | HFE-7500 + 2.2% ionic Krytox | 1619.72 | 1.613 | n/a |
|  | Outer | M9 salts + 25mM glucose + 2% Pluronic-F68 | 1017.9 | 1.563 | 0.4580 |
| PBS + 10% PEG 6000 mw | Inner | PBS + 10% PEG 6000 mw | 1013.7 | 3.431 | 0.4613 |
|  | Middle | HFE-7500 + 2.2% ionic Krytox | 1619.72 | 1.613 | n/a |
|  | Outer | PBS + 10% PEG 6000 mw + 2% Pluronic-F68 | 1014.1 | 6.395 | 0.4550 |

**Table 3:** Typical biological assays associated with each buffer.

| Surfactant System | Biological Relevance |
| --- | --- |
| PBS | Base salt buffer |
| PBS + 1% Tween-20 | Gentle cellular lysis (mammalian and bacterial) |
| PBS + 0.9% NP40 | Cellular lysis (scATAC-Seq and other NGS) |
| M9 bacterial media | Bacterial growth media |
| M9 + 25mM glucose | Bacterial growth media with carbon source |
| PBS + 10% PEG 6000 mw | Molecular crowding agent for ddPCR and other droplet molecular biology |

**Figure 2.**
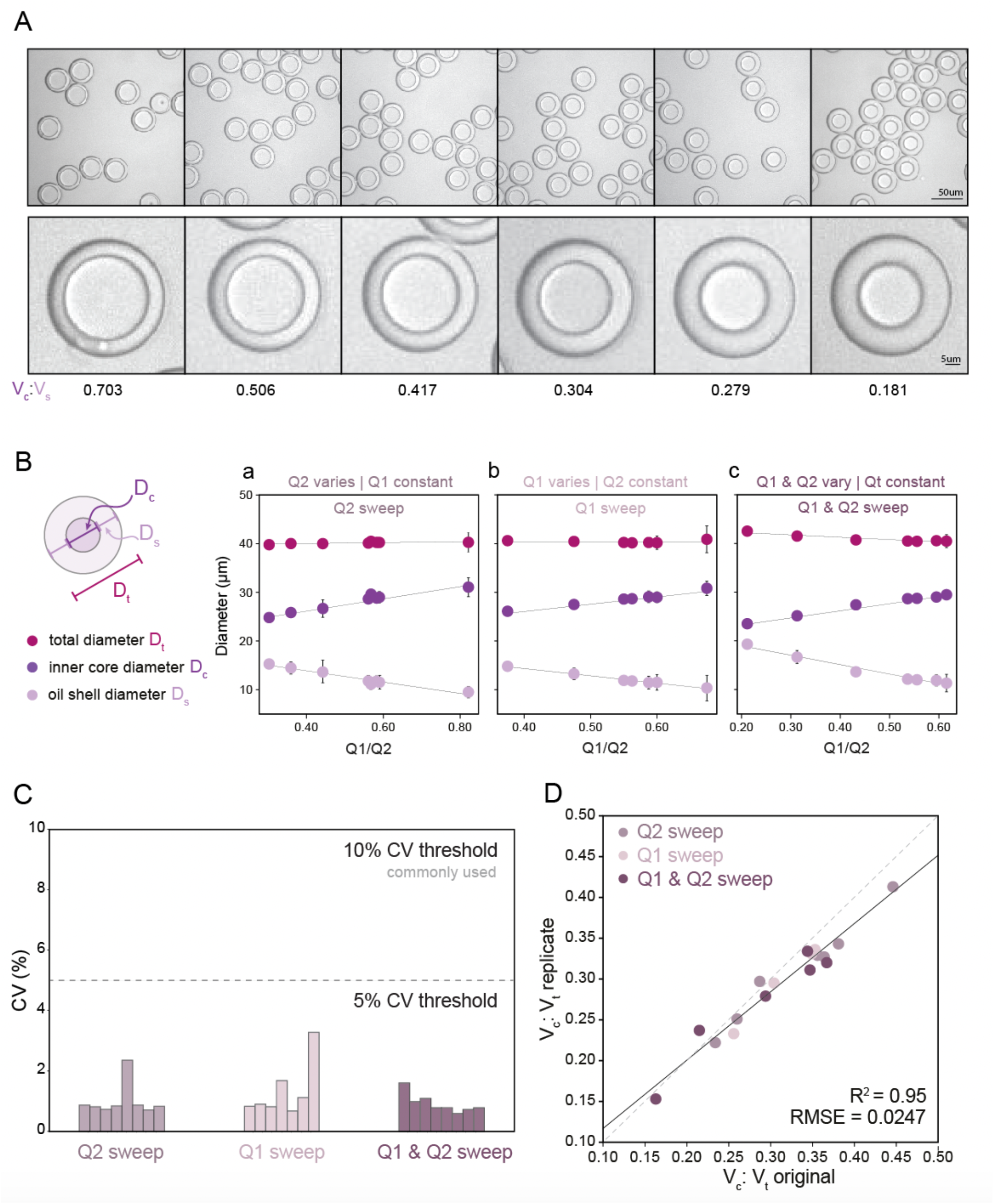
Measured droplet diameters as a function of flow rates for droplets containing PBS + 1% Tween-20. **(A)** Representative bright field microscopy images of monodisperse DE droplets across varying core: shell volume ratios. **(B)** Measured total diameters (magenta markers), oil shell diameters (lavender markers), and inner core diameters (purple markers) for flow conditions varying oil flow rate (Q2) only (left), inner aqueous flow rate (Q1) only (middle), or simultaneously varying Q1 and Q2 (right). Markers indicate median, error bars represent standard deviation, and solid lines show a linear regression. **(C)** Measured diameter coefficient of variation (CV) (%); see Table S5 for number of droplets for each flow condition. **(D)** Measured core: total volume ratios across 2 replicates for all flow conditions (see Table S5 for number of droplets per condition). Dashed line indicates 1:1 relationship, solid line indicates linear regression.

To test reproducibility of measured rates for these solutions and device geometry, we repeated all measurements using a new Dropception device in a different laboratory (**Fig. S3**). Measured diameters showed strong agreement across replicates (**Fig. 2D**, R^2^, residual norm. = 95%, 0.0247), demonstrating the ability to reliably use flow rates to specify aqueous core reactor diameters from 21.9 – 29.4 μm and overall droplet diameters from 38.6 – 41.0 μm for this surfactant formulation and device design. While previous investigations have probed 3-10 conditions, here we generated and quantified 31 unique monodisperse droplet populations for this surfactant system, demonstrating droplet stability across broad flow rate and droplet size ranges.

### 4. Investigation of common biological solutions reveals wide range of interfacial tension and viscosities

Next, we expanded our investigation of DE size to include 6 biologically relevant buffers and surfactants used in applications including bacterial growth assays, mammalian and bacterial cell lysis for NGS applications, and ddPCR (**Table 2**). Rheological and interfacial properties of fluorinated oils and biological buffers as typically used in droplet generation for dual layer structures and multi-surfactant systems have been largely understudied to the best of our knowledge.^1^ To probe this question, we first quantified the viscosity of each buffer using a cone and plate rheometer and the interfacial tension (IFT) of buffer/oil interfaces using pendant drop tensiometry (**Figs. 3, S7, Tables 2, S1**).^66,67^ These data reveal that: (1) these 6 different buffers span a wide range of viscosities (0.861-3.431 mPa-s) and IFT values (0.543 – 12.41 mN/m), (2) oil surfactants heavily influence IFT, driving a nearly 100-fold decrease in IFT (PBS in oil vs PBS in surfactant oil: 40 vs. 0.543 mN/m), (3) added salt initially lowers IFT and increases droplet deformation but high concentration can reverse this effect, and (4) outer sheath formulations, which contain both a non-ionic polysorbate surfactant and a long-chain detergent, have near-equal IFT, despite large variations in IFT in their respective inner solutions without these emulsifiers (**Fig. 3C, Table 2**). These results highlight the importance of oil surfactant in lowering interfacial tension and thus promoting droplet deformation. In addition, the observed nonlinear dependence of IFT on salt concentration is consistent with changes in the Hydrophylic-Lipophylic balance (HLB), the fraction of interface surfactant molecules in the oil versus aqueous phases (**Fig. S6**). These positional changes lead to an initial decrease and subsequent increase in IFT as salt concentration increases and pushes the molecule out of the aqueous phase into the oil phase^68,69^ (**SI Extended Discussion Note 2**).

**Figure 3:**
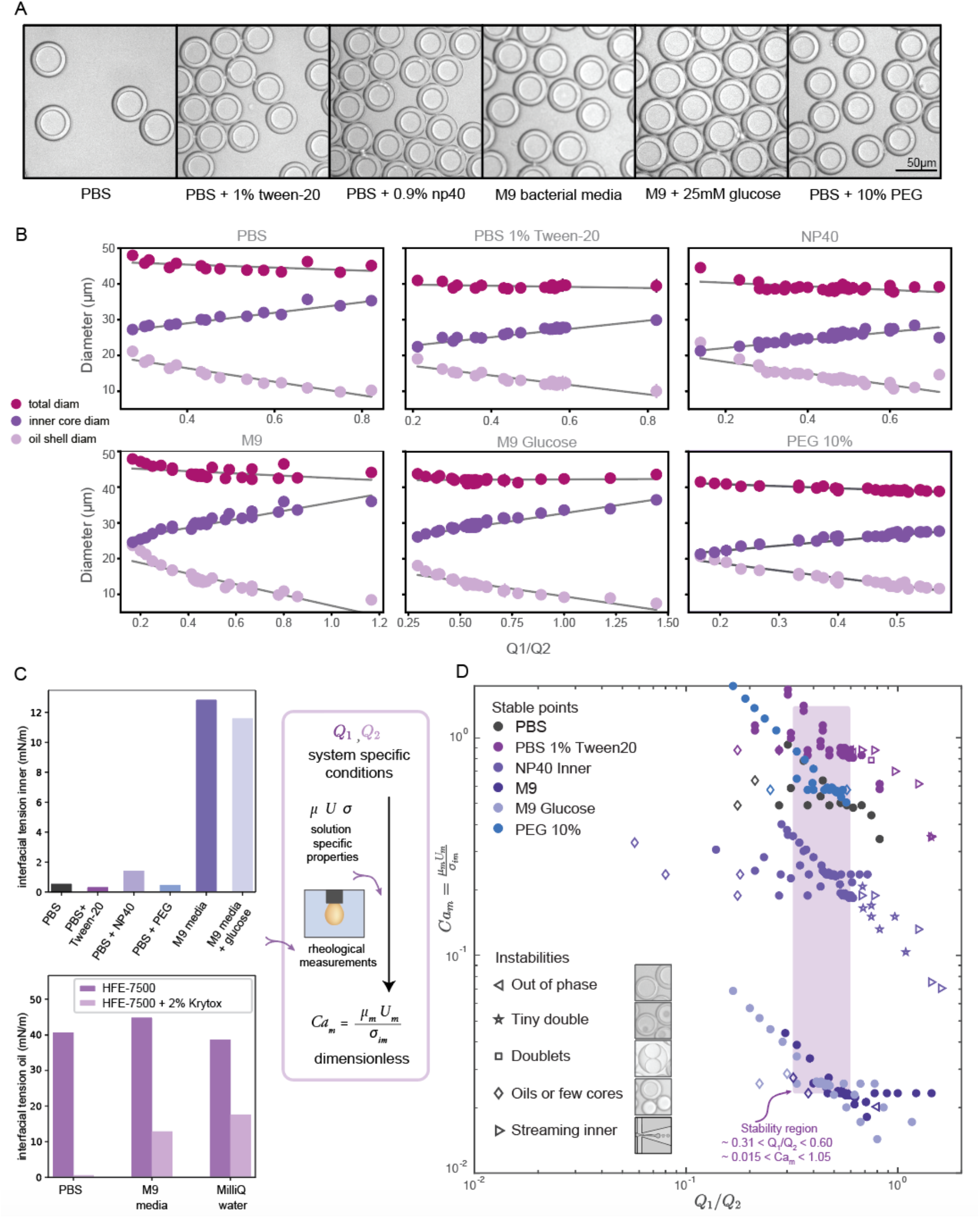
DE droplet stability and morphology across varying inner core solutions & surfactants. **(A)** Representative bright field images of monodisperse DE droplets containing 6 different biologically relevant inner solutions. **(B)** Measured total (magenta markers), oil shell (lavender), and inner core (purple) diameters as a function of flow rate ratio for all conditions for each buffer. Markers indicate median diameters (N for each condition in Table S5) and error bars indicate standard deviation. **(C)** Measured interfacial tension for 6 inner aqueous solutions in the presence of HFE-7500 oil + 2.2% ionic Krytox (top), and for HFE-7500 oil with (light purple) and without (dark purple) 2.2% ionic Krytox across 3 aqueous inner solutions; measurements can be combined with flow rates to calculate dimensionless Capillary numbers. **(D)** Capillary number Ca_m_ *vs.* Q1/Q2 flow rate ratio across all flow rate combinations and buffer conditions. Solid markers indicate conditions that yield monodisperse single core DEs; open markers indicate unstable conditions that yield droplets with mixed morphologies. Purple box indicates flow rate ratio range that produces stable monodisperse droplets across all conditions.

### 5. DE stability depends on physical properties of inner solution

Next, we conducted size parametric sweeps across each of these 6 biologically-relevant solutions (**Figs. 3A,B, S4, Table 1**). In total, we profiled thousands of droplets for each of 196 buffer and flow rate combinations (**Tables S5-7**). For each buffer condition, 14-30 flow rate combinations systematically sampled across the parameter space yielded monodisperse single core droplets, all with <5% CV in droplet inner and outer diameter (**Fig. S5**), the gold standard for robust droplet generation. As seen previously (**Fig. 2**), droplet volume and diameter varied linearly with changing Q1 and Q2 and total droplet volume remained relatively constant (**Figs. 3B, S4**). Consistent with prior reports for two-step formation devices, we observed only two periodic DE generation regimes, dripping-dripping and dripping-jetting formation.^1^

The flow rate ranges over which each buffer yielded stable single core droplets differed dramatically (**Table S5**). For example, DEs containing a high viscosity 10% PEG inner buffer were stable over a much narrower range of Q1 flow rates than those with the M9 high-salt media containing sugars (150–220 μl/hr *vs.* 150–350 μl/hr, respectively) (**Table S5**). Furthermore, the range of stable flow rates often differed significantly between Q1 and Q2: for instance, 10% PEG DEs have a very narrow range of stable flow rates in Q1 (150-220 μl/hr) but a large range of stable flow rates in Q2 (350-1200 μl/hr) (**Table S5**). These differential effects on droplet stability may be explained by the interplay between viscous stresses and interfacial stresses acting on the fluid streams, as disruptions in this balance lead to instabilities, deviations from ideal flow behavior.^1,24^ For single core monodisperse droplets, we define ideal flow behavior as the dripping-dripping flow regime with matching periodicity between W/O droplet generation at FF1 and W/O/W droplet formation at FF2; deviation in flow behavior, or instabilities, are caused by changing flow dynamics at the flow focusers, due to differing fluid properties with differing surfactants (**Figs. S8, S11**).

To quantify this balance of forces in the flow behavior, we then used these flow rates and the measured fluid properties to calculate the Ca number for each condition, which specifically quantifies the strength of viscous forces relative to cohesive forces at an interface. For DE droplets, Ca_m_ determines the formation behavior of inner droplets and the pinch-off mechanics of the fast-flowing middle oil phase while Ca_i_ and Ca_o_ focus on the properties of the inner and outer phases respectively:^24,25^

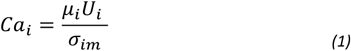

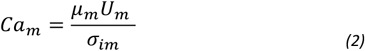

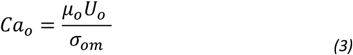

where *μ* is the dynamic viscosity of the liquid, *U* is the characteristic velocity (here a quotient of flow rate Qx and nozzle cross sectional area), and σ is the interfacial tension between two phases.

When visualized non-dimensionally in a plot of Ca_m_ vs. the Q1/Q2 ratio (**Fig. 3D**), different types of droplet instabilities reproducibly cluster together across conditions. Low Q1 relative to Q2 typically leads to oil-only coreless droplets or satellite oils, independent of Ca_m_ (**Fig. 3D**, *left*). At a high Q1 relative to Q2, Instability type varies with Ca_m_, with high likelihood of streaming inner phase instabilities giving way to higher likelihood of multi-core (tiny double and doublet) instabilities as Ca_m_ decreases (**Fig. 3D**, *right*). Droplets containing fluids with higher Ca_m_ transition to streaming and other instabilities at a lower Q1/Q2 ratio than those with lower Ca_m_, while low Ca_m_ systems result in coreless droplets at higher Q1/Q2 ratios than high Ca_m_ systems. These droplet morphologies result from deviations from ideal flow behavior into unmatched periodicity at FF2 and the dripping-jetting regime (**SI Extended Discussion Note 3**).

Across all buffer conditions, we identify a single range of flow rate ratios that universally yields stable monodisperse single core droplets (**Fig. 3D**, purple box; approximately 0.31 < Q1/Q2 < 0.60, 0.015 < *Ca*_*m*_ < 1.05, for similar flow rates), providing an initial flow rate combination to use when trying to optimize DE generation in novel biological systems.

### 6. Universal size law reveals heavy dependence on mass conservation by flow rate for monodisperse droplets

A single equation capable of accurately predicting droplet size as a function of fluid properties and flow rates could significantly reduce the time, effort, and reagents required to identify optimal flow rates for a new buffer condition. Wang et al1 recently conducted an extensive computational study of formation dynamics in dual flow focuser devices, and developed a size scaling law for DE droplets formed in a one-step regime as a function of flow rates and fluid properties, building on previous size scaling laws.^63,64,70^ Here, we test if this law can be applied to predict droplet sizes for single core DE droplets produced within our two-step formation device geometry. Five empirically-derived fit parameters allow the model to accurately predict the relative volume of the droplet core to the total size (V_c_:V_t_ core: total volume ratio) for 95% of the observations across all 138 conditions within an 10.7% margin (**Fig. 4A, Table S8**, Model A), with normally distributed residuals (**Fig. 4B**):

**Figure 4.**
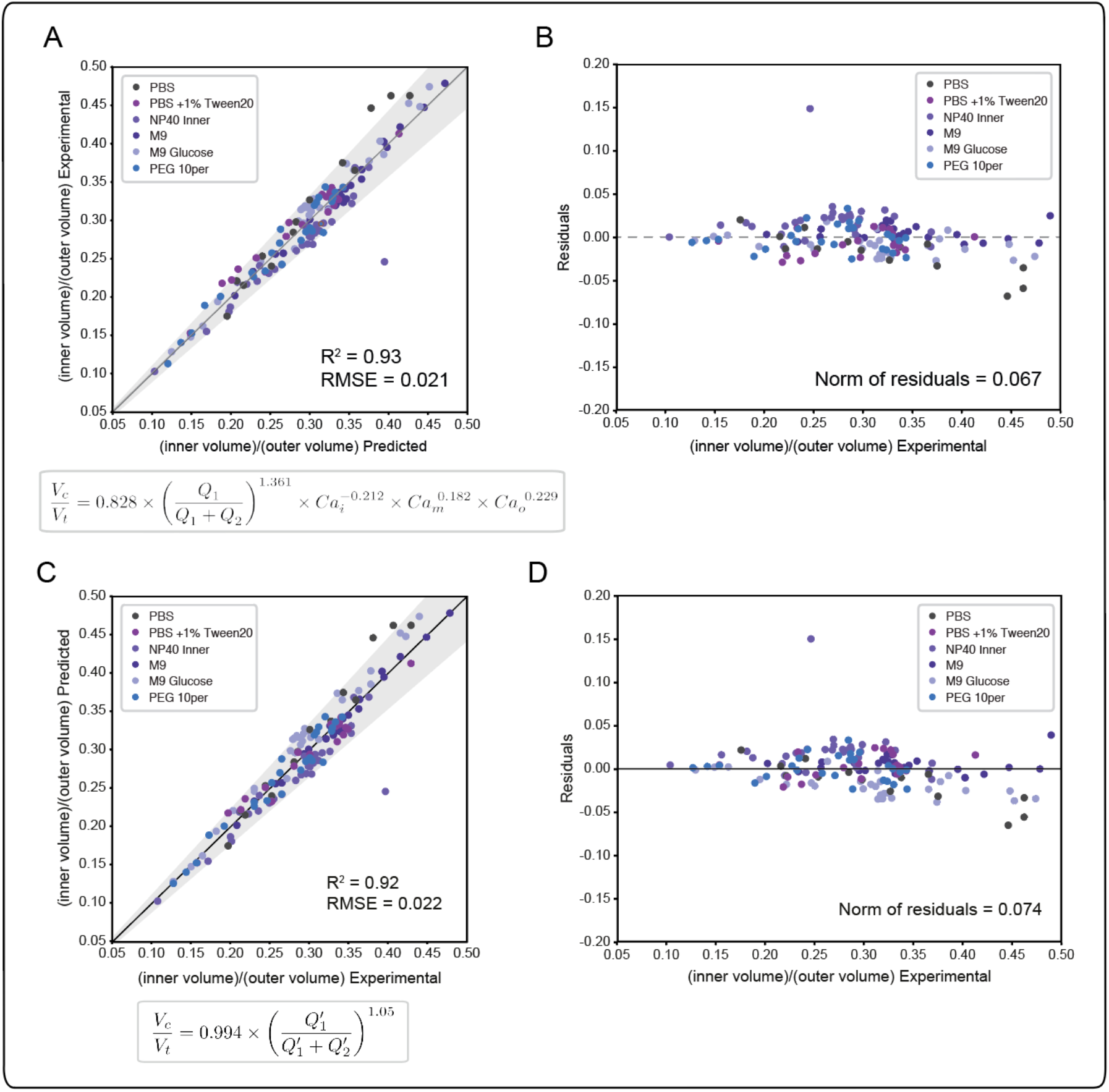
Universal size scaling law capable of predicting volume ratio as a function of flow rates and capillary numbers. (Left) Predicted vs experimental core volume: total volume ratio for 196 droplet conditions with buffer composition indicated by marker color. Dark line indicates 1:1 line; light line indicates linear regression; grey shading indicates confidence interval; universal size scaling law equation is shown below; markers indicate median value per condition with number of droplets listed in Table S5. **(Right)** Calculated residuals of experimental results from model as a function of volume. **(A,B)** Model A, a general form with capillary number dependence - most broadly applicable. Explains 95% of observations within a 10.7% interval. **(C,D)** Model B, without capillary numbers, a simplification toward ideal mass conservation that is only possible with our restrictions to ideal flow behavior. Explains 95% of observations within a 11.6% interval.

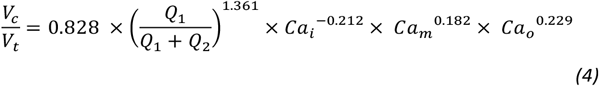

In this size law, the relative volume of the droplet core compared to total volume has a high dependence on relative flow rate contributions of the inner solution (Q1) to oil (Q2), with small exponents on Ca_m_ and Ca_o_ intimating little dependence on the outer sheath (Q3). Because Q3 is held constant throughout and thus measured outer droplet diameter shows only small deviations, this size law also returns information about absolute droplet size (**Fig. 3B**).

Our restrictions to ideal flow behavior then allow us to simplify the model to exclude capillary number dependence via reduced and zero exponents for Ca_x_ terms (**Fig S9B, Tables S8, S9**, Model B). A simplified model resulting from the best fit of these analyses explains 95% of observations within a margin of 11.6% (**Tables S8, S9**):

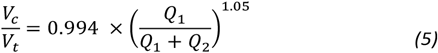

Interestingly, we find that our model incorporating capillary number terms performed only slightly better than the simplified model that does not (R^2^ = 92.5% vs. 93.1%, interval of explained observations = 10.7% vs. 11.6%, **Fig. S9, Table S8**), suggesting that droplet size is determined primarily by simple mass conservation. Ideal mass conservation leads the prefactor and exponent to both be 1, and we observe the fitted parameters are very close to 1. This phenomenon is because generation of single-core double emulsions using our two-step flow regime approaches ideal behavior; our dual linear flow focuser device geometry promotes dripping-dripping and, because FF1 and FF2 are decoupled by a flow resistor, allows a user to easily match periodicity of single emulsion generation at FF1 to double emulsion encapsulation at FF2 (matched periodicity at Q3 pinch off) by adjusting relative Q1 and Q2 contributions (**Fig. S11, SI Extended Discussion Note 3**). When instabilities that depart from this ideal behavior (**Fig. 3D**) are included, the model deviates (**Fig. S10**). Thus, the simple mass conservation model and absolute flow rate conditions explored here, generated with our restrictions in place, apply across systems, where these assumptions are maintained. Where analyses lead to deviations from this procedure, the model can be used to provide specific predictive value (as shown in **Fig 5**) if flow rates are restrained to the stable range (**Fig 3D**) and minor adjustments in Q1 & Q2 are made to match periodicity with Q3 to achieve monodisperse single core droplets. An alternative capillary number based model expression without a prefactor and an inner core volume model had equivalent and less optimal fits respectively (**SI Extended Discussion 4, Fig. S9, Tables S8, S9**). The more general capillary number-based model is important for occasions when the ideal flow behavior breaks down, at instability boundaries and in schemes outside our restrictions.

**Fig. 5.**
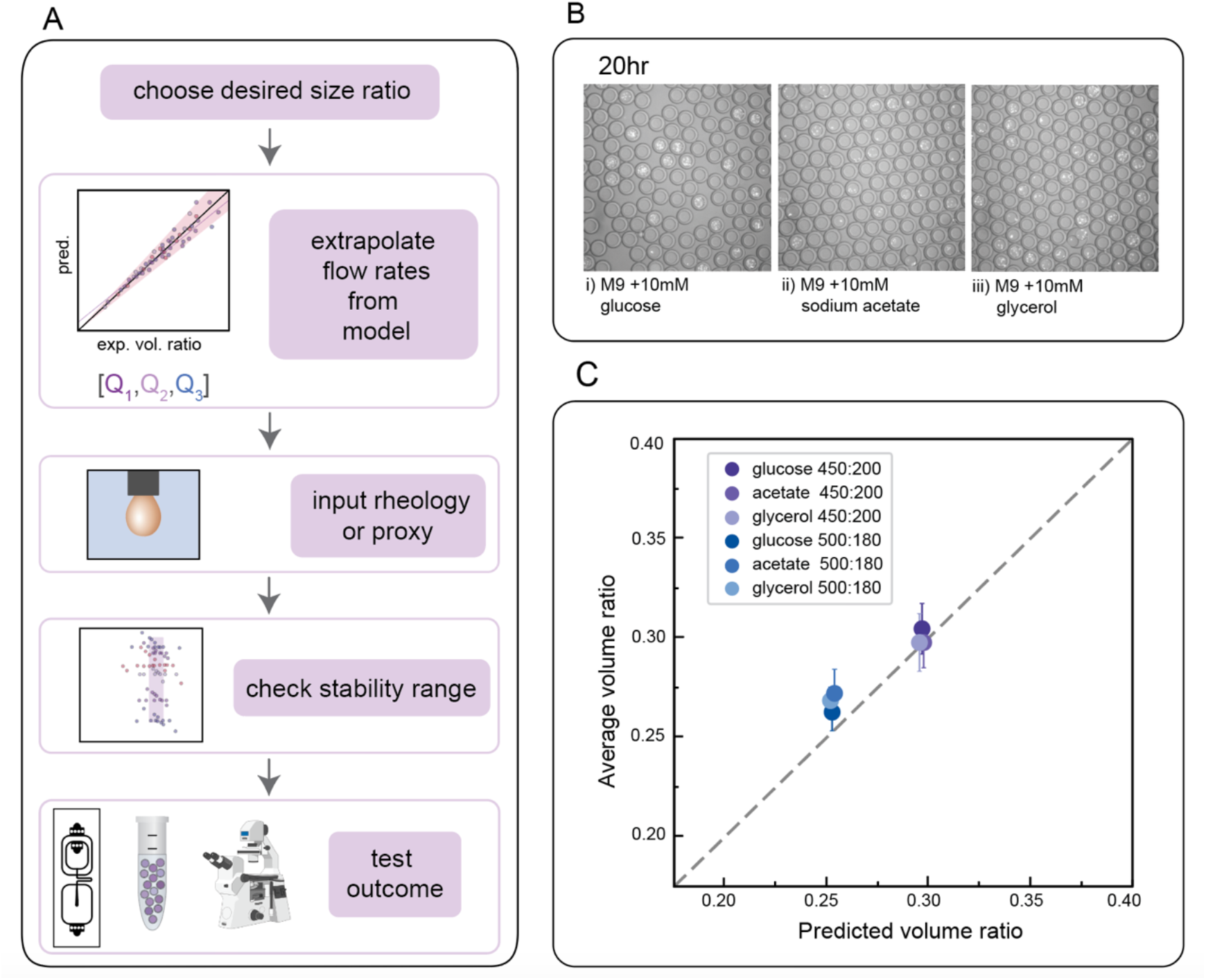
Predicting flow rates required to produce droplets with a desired geometry for anaerobic *E. coli* growth assays. **(A)** Schematic of workflow to test accuracy of flow rate predictions from universal size scaling laws. **(B)** Representative microscopy images of *E. coli* growth in DEs containing M9 media plus three different carbon sources after 20 hours. **(C)** Measured *vs.* predicted core: total volume ratios. Markers represent median of 10 droplets and error bars show standard deviation. Dashed line is 1:1 relationship.

These data and corresponding models are a valuable resource for flow rates that generate single core DE droplets suitable for FACS regardless of surfactant, when using this two-step device with conditions of flow regime and periodicity satisfied.

### 7. Size law accurately predicts size in de novo biological use cases

Finally, we directly validated the predictive power of this universal size scaling law via blind predictions of droplet stability and size for a biological assay using a previously uncharacterized inner buffer. Microfluidic encapsulation of single bacteria into droplets can enable a wide variety of applications, from high-throughput screening for media conditions that allow recovery of previously uncultured bacteria from metagenomic samples to screens for antibiotic resistance strains in a mixed community.^71–74^ Such growth assays typically require M9 media supplemented with physiologically-relevant carbon sources (*e.g.* glucose, sodium acetate, or glycerol) to tune relative bacterial growth rates. To test if our universal size scaling law can help identify conditions that produce droplets of a desired size and shell thickness (mimicking the same process a user might employ), we: (1) specified 2 desired final core volume: total droplet volume ratios (0.25 and 0.30), (2) measured the dynamic viscosity (μ) and interfacial tension (σ) for M9 with either 10 mM glucose, sodium acetate, or glycerol catabolic substrates in the presence of HFE-7500 supplemented with 2.2% ionic Krytox, (3) solved for Q1 and Q2 flow rates that would generate droplets with the desired volume ratios using both models, which returned similar flow rates, (4) generated DE droplets encapsulating a GFP-expressing *E. coli* strain bacteria in each media with these specified flow rates, and (5) quantified actual droplet size and shell thickness (**Fig. 5A**).

Flow rates suggested by the models fell within the range previously found to generate stable droplets in the size characterization data (**Fig. 3D**). For all 3 media formulations, flow rates of [200, 450] and [180, 500] (Q1, Q2) yielded monodisperse single core droplets with few aberrations (**Fig. 5B,C**); measured volume ratios were consistent with predictions within 10.5%, 14.9% and 11.3% variation for M9 with 10mM glucose, sodium acetate, and glycerol, respectively, for 95% of observations for the model considering only mass conservation (Model 1C), and 8.3%, 10.0% and 10.6% for the model incorporating capillary numbers (Model 1A). These results thereby validate the primary dependence on mass conservation seen in the systematic sweep data (**Fig. 5D**). As expected, *E. coli* cells cultivated in double emulsions for 20h at 37°C grew on all three catabolic substrates in both flow conditions (**Figs. 5, S12**). The agreement between predictions and empirical measurements of DE size demonstrates, along with direct flow rate options, the utility of these tools in developing novel biological applications for single core double emulsions.

## Conclusions

Double emulsion droplets are nanoliter-to-picoliter volume reactors that can be sorted by their fluorescence using commercial flow cytometers, and therefore provide a powerful tool for ultra high-throughput biochemical and cellular assays. However, production of DE droplets has long been considered technically challenging due to their complex 3-phase architecture (an inner aqueous core surrounded by an oil shell suspended within an aqueous outer buffer) and the importance of DE droplet size and shell thickness for downstream assay efficiency. Here, we performed systematic parameterization of droplet size and shell thickness over 196 different combinations of flow rates and inner buffers (>22000 DE droplets in total) using a straightforward two-step device. Our restrictions to a dripping-dripping regime and matching Q3 periodicity focused our investigation on single core monodisperse droplets, essential for FACS applications, and bulk and interfacial characterization of each buffer led to identification of a consistent stable flow rate ratio range. With these data, we applied and empirically validated a recent size scaling law to accurately predict DE droplet size and shell thickness as a function of flow rates and buffer properties. This dataset and model establish a valuable resource for implementation of new assays by streamlining identification of appropriate flow rates.

Further expansions of this work to additional surfactants, including reagents such as BSA and proteins that have more complex viscoelastic or skinning effects, could allow better understanding of these complex flows and further simplify novel assay development. Additionally, oil surfactants have a large effect on reaction efficiency and cell viability in droplets. Here, we held our oil formulation constant to that optimized in our prior DE FACS workflow, but effects of oil variation on droplet size and stability remain underexplored. While it seems that droplets of varying shell thickness do survive FACS, thick-shelled droplets appear anecdotally to run more slowly and cause more clogging; quantification of these trends would assist in FACS implementation. Lastly, adjustments of Q3 for emulsions containing multiple cores may have utility in complex biological systems and would be a logical expansion of this model.

As one of the first in-depth parameterizations of DE size and stability and with an explicit focus on monodisperse single core droplets, we anticipate that this systematic dataset and size scaling model will allow users to reliably generate DE droplets with a desired size phenotype for HT biological screens via FACS.

## Supporting information

Electronic Supplemental Information

## Acknowledgements

The authors thank Rochelle Radzyminski and Margarita Khariton for their help with image processing and discussion during the formulation of this study. We are grateful to all members of the Dropception team as well as Fordyce, Fuller and Kusumaatmaja laboratories for their advice and helpful feedback on the manuscript. S.C. acknowledges funding as a ChEM-H CBI fellow and a Stanford Graduate SGF fellow. K.B. acknowledges funding as ChEM-H CBI fellow, an NSF GFRP fellow, and a Siebel Scholar. A.M. acknowledges funding as a Simons Foundation Postdoctoral Fellow in Marine Microbial Ecology (award #600755). P.M.F. is a Chan Zuckerberg Biohub Investigator and this work was supported by NIH DP2-GM123641 awarded to P.M.F.

## Materials and Methods

### A. Device Lithography

All designs were generated in AutoCAD (Autodesk). All designs used in the study at all stages of iteration are available on an Open Science Framework project. Devices were fabricated via standard soft lithography protocols as described previously.^16,31^ Selective hydrophilic wettability was imparted to FF2 via air plasma treatment for 10 minutes with tape applied to cover outlet and outer sheath inlets as described previously.^31,75^

### B. Droplet Generation

DEs were generated using 3 syringe pumps (PicoPump Elite, Harvard Apparatus) for the inner, oil, and outer carrier fluids. Syringes (1 – 10 mL; PlastiPak plastic syringes, BD) were connected to the microfluidic device with polyethylene tubing (PE/2, Scientific Commodities). Droplet generation rates were typically 1-10 kHz. For the initial condition (**Fig. 2**), the inner phase for the aqueous droplet core was composed of 1% Tween-20 (Sigma) and FITC-BSA in 1x PBS (Invitrogen). For all measurements, the oil phase was composed of HFE-7500 (Sigma) and 2.2% Ionic PEG-Kyrtox (FSH 157, Miller-Stephenson) and the outer phase was composed of 1% Tween-20 (Sigma) and 2% Pluronix F68 (Kluplour 188, Sigma) in PBS. Typical flow rates were 400:230:6500 (O:I:C) μL/hr. Droplet generation was monitored and recorded via a stereoscope (Amscope) and high-speed CMOS camera (ASI 174MM, ZWO). Droplets were stabilized for 4 minutes prior to a set collection time of 6 minutes. At each condition, we acquired a 500-frame video to assess stability and breakoff phenotype.

### C. Study design – Choice of flow rates per buffer

For each condition, we first identified a flow rate ratio that produced monodisperse droplets with a core volume:shell volume ratio close to 0.60. Next, we generated droplets at 4 flow rate combinations closely spaced around this initial condition (+/− 5 μl/hr), then varied flow rates by larger intervals to scan for the limits of the flow rate ratio regime capable of stably generating monodisperse droplets. For the Q_t_ sweep, we divided the expected stable range of Q1 or Q2 into 6 parts and inversely varied Q1 and Q2 to maintain a constant Q_t_. In each condition, we stabilized for 4 minutes and then collected DEs for 6 minutes, allowing for assessment of a statistical population via microscopy. Each set of flow rate ratios contained >10 stable points and >5 unstable points across the bounds of the dynamic range recorded (**Table 1**).

### D. Image Acquisition

Prior to imaging, we loaded approximately 2 μl from the droplet pellet and 8 μl of aqueous outer solution into Countess cell counting chamber slides, forming a droplet monolayer. We then imaged droplets on a Nikon Ti Eclipse microscope at 10X magnification using both brightfield illumination and a FITC-compatible fluorescence channel. Multiple images were acquired per droplet population (> 10 images per condition) in non-overlapping FOVs for FOVs containing a single layer of DE droplets. We found that accurate focus and waiting for droplets to stabilize to minimize drifting were important for accurate image analysis downstream. For each condition, we acquired a 500-frame video of droplet generation using ASICAP software just after starting droplet collection.

### E. Droplet size characterization

Droplets from 10 images per flow condition were characterized in MATLAB using a custom image processing pipeline available in our Open Science Framework: https://osf.io/pt6qu/?view_only=f1690e6efd7a4773b7e26fec5a65aada.

### F. Interfacial tension measurements: Pendant drop

Interfacial tension (IFT) of each solution was measured via pendant drop tensiometry using drop shape analysis^66,67,76^ As the oil is denser than the inner solutions, we utilized a pendant oil droplet suspended within an inner aqueous buffer solution for the analysis (**Fig. S7**). Oil droplets were formed and suspended from a syringe with a 27 gauge metal capillary nozzle, within a 5 mL bulk of an inner solution. Droplet shape analysis using a custom Matlab code was conducted when the droplet was as stable as possible, as is established in the rheology literature.^66,76^ At equilibrium with a stationary drop, cohesive forces (interfacial tension) and gravitational deformation are balanced. Therefore, the simplified Young-Laplace equation and the hydrostatic pressure can be equated and solved, giving interfacial tension. Final IFT measurements were the mean of calculations from 3-4 analyzed drops.

### G. Viscosity Measurements

Dynamic viscosity for each solution was measured using a commercial rotational cone and plate rheometer. Measurements were obtained by conducting a logarithmic flow sweep across 4 orders of magnitude of shear rate (2.86479 – 2864.79 Hz) with a 2° cone at 20 °C. The average viscosity in the linear regime is reported as the shear rate independent viscosity of the medium.

### H. Size Scaling Law Fitting Method

For each model, a function that accepts capillary numbers, flow rate ratios, and unknown coefficients was defined in Python3. The curve_fit function in the SciPy optimization package was used to estimate best fit values for these coefficients based on experimental data. Depending on the desired prediction, either inner-to-outer volume ratios or inner volumes as the dependent data were calculated based on known capillary numbers as the independent data. In all cases, we used an initial guess of 1 for each parameter. The curve_fit function returned optimal values for the unknown parameters and their estimated covariance.

### I. E coli growth - Biological test case Method

*E. coli*-GFP was routinely cultivated on Luria-Burtani (LB) agar plates supplemented with 30 μg/mL kanamycin and liquid M9 basal medium with 25 mM glucose at 37°C. Overnight stationary phase cultures were washed twice with basal M9 medium to remove any residual carbon source. Washed cells were diluted to an optical density (OD600) ^~^ 0.05 in the inner solution to achieve single cell loading based on a Poisson distribution. Diluted cell suspensions were supplemented with either 10 mM of glucose, acetate, or glycerol and used as inner solutions. The outer solution contained M9 medium supplemented with 2% Pluronix and 1% Tween-20 for stable droplet formation. After generation of double emulsions, the outer solution was replaced with M9 medium containing only 2% Pluronix to ensure *E. coli*-GFP grew on the packaged substrate, rather than catabolizing Tween-20 surfactant. E. coli-GFP double emulsions were incubated statically at 37°C overnight. Microscopy samples were imaged at 0 and 20 hr timepoints to measure droplet sizes and *E. coli*-GFP fluorescence.

