## Supplementary material for "Systematic Characterization of Double Emulsion Droplets for Biological Applications": Electronic Supplemental Information

Please do not adjust margins

##### Supplemental Information

1. **Extended Discussions**

1. **Extension Discussion Note 1:** Droplet Generation Protocol
2. **Extension Discussion Note 2:** Hydrophilic Lipophilic balance and Implications on Interfacial properties of Biological Solutions
3. **Extension Discussion Note 3:** Instability types
4. **Extension Discussion Note 4:** Inner core volume size scaling law

2. **Supplemental Figures**

1. **Fig. S1:** Dropception DE generation device
2. **Fig. S2:** DE systematic characterization design and absolute flow rates
3. **Fig. S3:** Replicate experiment measuring DE diameters for PBS + 1% Tween-20
4. **Fig. S4**: Measured DE droplet diameters and volumes with 6 inner aqueous buffers
5. **Fig. S5:** Measured Diameter Coefficient of variation CV% across flow rates and solutions
6. **Fig. S6:** Schematic illustrating Hydrophilic Lipophilic Balance (HLB) and potential impact on interfacial tension (IFT)
7. **Fig. S7:** Pendant drop method to measure interfacial tension (IFT)
8. **Fig. S8:** Example images showing how varying flow rates alter droplet morphology and stability
9. **Fig. S9:** Additional possible size scaling laws and comparisons between predictions and measurements.
10. **Fig. S10:** Goodness of Fit for simplified mass conservation size scaling law
11. **Fig. S11:** Stability and instability types in DE droplet formation
12. **Fig. S12:** Representative microscopy images of DE droplets containing fluorescent E. coli

13. **Video S1:** DE Generation using the Dropception device

#### 3. Supplemental Tables

1. **Table S1:** Extended interfacial and bulk fluid parameters
2. **Table S2:** Salt concentration sweep
3. **Table S3:** E. coli experiment solutions
4. **Table S4:** PBS + 1% Tween-20 Original and replicate
5. **Table S5:** All trials no replicates with fluid property parameters
6. **Table S6:** Replicate sweeps with fluid property parameters
7. **Table S7:** Instability condition types
8. **Table S8:** Goodness of fit comparison between volume ratio models
9. **Table S9:** Goodness of fit for each trial

### Extended Discussion: Note 1

#### 1. Droplet Generation Protocol

1. Prepare all solutions listed in Table ED1.
2. Prepare syringes and cut tubing listed in Table ED1. Consistency of length and tautness are crucial for tubing, whereas absolute length is not.
3. Screw needle nozzles on syringes and attach tubing to needle.
4. Situate syringes in syringe pumps, being sure to attach them securely.
5. Set flow rates with outlet, inner, and oil phases at flow rates identical to, lower, and higher, respectively, than the target central condition.
6. Fast forward syringe pumps until liquid is at the end of the tubing.
7. Prepare an Eppendorf tube for collection of outlet tubing output.
8. Selectively treat the outlet and outer solution inlet with air plasma for 10 min according to Brower et al[1].
9. Immediately set device on scope, insert outer tubing into outer solution inlet and begin flow of the outer solution to preserve hydrophilicity of the PDMS.
10. Insert tubings from outlet and other syringe pumps into the device
11. Allow droplet generation to stabilize over several minutes. Be patient!
12. Monitor for reflux of the other phase into the inner and oil channels. If reflux occurs, immediately reduce the flow rate of the offending phase and increase the flow rate of the other phase. Extended time in contact with the incorrect phase can permanently change the wettability or hydrophilicity/hydrophobicity of the channel walls, causing problems in subsequent droplet generation.
13. If required, pressing on the device at the inlets or flow focusers can help remove bubbles or reflux, and help instigate consistent flow through all channels. However, be careful, as this can also damage the device if too vigorous.
14. Let setup stabilize for 30 min to allow device and syringes to equilibrate before conducting experiments if accurate and consistent droplet sizes are important.
15. After each change in flow rates, allow droplet production to stabilize for at least 4 minutes. Here, we stabilized for 4 minutes and then switched to a clean vial and collected for 6 minutes.
16. Devices that were plasma treated and filled with outer solution but not used for droplet generation can be prepared for reuse by flowing water through the device through the outer solution inlet and drying the device on a hot plate at 65°C for 20 min or more.

**Table ED1:**

| **Solutions** | **Details** |
|---|---|
| Inner solution | PBS + 1% Tween-20 |
| Oil | HFE-7500 + 2.2% ionic Krytox FSH-157 |
| Outer sheath | PBS + 1% Tween-20 + 2% Pluronix-F68 |
| **Syringes** | |
| Inner solution | Plasti-pak plastic, 1mL |
| Oil | Plasti-pak plastic, 5 mL |
| Outer sheath | Plasti-pak plastic, 10 mL |
| **Tubing** | |
| Inner solution | $1^{st}$ from left. 41cm length (for this study) |
| Oil | $2^{nd}$ from left. 56cm length (for this study) |
| Outer sheath | $3^{rd}$ from left. 54.5cm length (for this study) |
| Outlet | 15cm length (for this study) |

#### Extended Discussion: Note 2

##### 2. Hydrophilic Lipophilic balance and Implications on Interfacial and Bulk properties of Biological Solutions

Investigating variation in the physical properties of the solutions involved is important to understanding DE formation regimes and the resulting droplet size and shell thickness. Interfacial tension (IFT) of droplet systems with HFE-7500 oil is typically 1-5 mN/m,[2] but our custom oil has a low baseline IFT with 1% Tween-20 in PBS of 0.319 mN/m.

Measuring the interfacial tension of our solutions revealed a nonlinear dependence on salt, with an initial decrease and a subsequent increase in interfacial tension as salt concentration increased. We hypothesize this stems from changes in the Hydrophylic-Lipophylic balance (HLB) of the surfactant, where variation in salt concentrations physically alter the relative fraction of interface surfactant molecules in the oil versus aqueous phases.[3,4] Surfactant molecules on the interface initially progress from primarily existing in the aqueous phase at zero salt concentration to protruding more into the oil phase as salt concentration increases, thereby decreasing the interfacial tension. This decrease in interfacial tension continues until the surfactant molecules have half their geometric length in each of the oil and aqueous phases, attaining the minimum surface energy and thus the minimum interfacial tension. Consistent with prior reports that the HLB can reduce IFT 10-100x, we see 2- and 20-fold decreases in IFT from MilliQ water to M9 inner (24.3 mM salts) and PBS inner (37.9 mM salts), respectively[4] (**Table S2**). As salt concentrations increase further, the surfactant molecules move further into the oil phase and the interfacial tension increases once again (as for 10x PBS, **Table S1**).[3] Further supporting this hypothesis, combining these buffers with HFE-7500 in the absence of surfactant gives much higher IFT values (**Table S2**). The outer phase has an increased total amount of surfactant with the addition of Pluronix F-68; likely via speeding the decrease to the minimum IFT, this results in lower measured interfacial tension, including a 20-fold IFT decrease at intermediate salt concentrations from M9 inner to M9 outer.

The viscosities of the tested solutions are comparable to water (0.890 mPa s). The only exceptions to this are solutions with Polyethylene glycol (PEG), a known viscosity modifier[5], which increases the viscosity of the solutions to 3.43 mPa s. The effect of PEG is enhanced in the outer solution, nearly doubling to viscosity 6.40 mPa s, likely due to the cooperative formation of larger polymeric structures (e.g. mixed micelles) with Pluronix molecules.[5]

#### Extended Discussion: Note 3

##### 3. Instability types

A dual-flow focusing device geometry allows for a dripping-dripping droplet generation regime where DE droplets are formed in 2 stages: W/O generation in FF1 and W/O/W shell formation in FF2. In this regime, monodisperse droplet generation requires matching the periodicity of droplet pinch-off at each flow focuser.

The ideal time needed to generate each DE is then t/N, where t is the flow time and N is the number of droplets generated during time t (**Fig. S11**). The volume of the inner droplet is given as Q1 * t/N, where Q1 is the flow rate of the inner solution. Under the same assumptions, the volume ratio of the core to total outer droplet is given by

$$\frac{V_c}{V_t} = \frac{Q_1 \cdot t/N}{Q_1 \cdot t/N + Q_2 \cdot t/N} = \frac{Q_1}{Q_1 + Q_2}$$

where Q2 is the flow rate of the middle phase, $V_c$ is volume of the core (inner droplet) and $V_t$ is volume of the total droplet. The ideal mass conservation expression we see here is approached by Model B in Section 6 as prefactor and flow rate ratio exponent both are very close to 1.

Two possible deviations from an exact dependence on flow rate ratios can alter droplet size and shell thickness.[6–8] In the first case, droplet formation may be periodically slowed down, resulting in clogging of inner droplets and some portion of the ideal inner droplet volume contributing to its preceding droplet. We attribute this to a large $Ca_m$, small $Ca_i$, or a small $Ca_o$, and represent it as follows:

$$\frac{V_c}{V_t} < \left(\frac{V_c}{V_t}\right)_{ideal} = \frac{Q_1}{Q_1 + Q_2}$$

In the second case, the middle phase fluid is pinched off faster than expected, resulting in satellite droplets that consist only of the middle phase. We attribute this to a small $Ca_m$, large $Ca_i$, or large $Ca_o$, and represent it as follows:

$$\frac{V_c}{V_t} > \left(\frac{V_c}{V_t}\right)_{ideal} = \frac{Q_1}{Q_1 + Q_2}$$

#### Extended Discussion: Note 4

##### 4. Inner core volume size scaling law

Conversely to the volume ratio models discussed in section 6 (**Figs. 4, S9, Tables S8, S9**), capillary numbers become more influential in modelling inner droplet volume directly. Following prior work from Garstecki et al and Liu and Zhang,[9,10] we use the scaling model

$$\frac{V_c}{h \times w \times w} = \left(0.270 + 0.760 \times \left(\frac{Q_1}{Q_2}\right)\right) \times \ Ca_m{}^{-0.0672}$$

The two terms in the bracket represent contributions from the blocking and squeezing phases during droplet formation. The two scalar parameters empirically account for the fact that the droplet length in the resistor is not exactly identical to the channel width, and that in the squeezing regime some of the middle phase fluid bypasses the inner phase fluid.[9] We also include a power-law relation with the capillary number of the middle phase.[11] Note that we have nondimensionalized $V_c$ by the dimensions of the oil inlet. Because this model is derived from a T-junction regime for single emulsion generation rather than a dual-flow focuser, it makes no explicit assumption about Q3 as in the volume ratio model. This model accurately predicts inner droplet volumes within a 31% margin with a less optimal fit than our prior model ($R^2$: 0.70, RSME: 0.14)(**Fig S9, Table S9**). Similarly, the model has worse performance with high salt buffers, with biased residuals in these cases (**Fig. S9**). However, this unbiased approach to inner core volume may prove useful for end users wishing to modulate Q3 explicitly and within buffers of low $Ca_m$ (e.g., PBS, PBS-Tween, etc).

#### Figure S1: Dropception DE generation device.

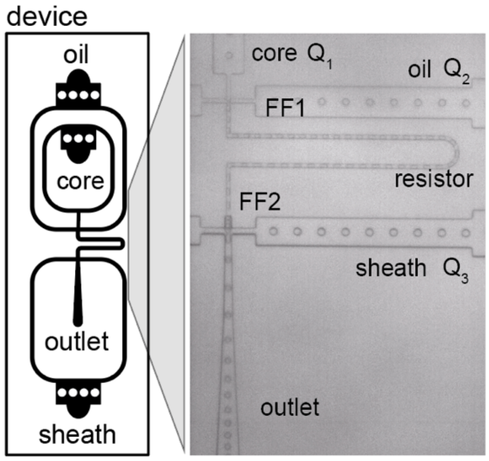

**Figure S1: Dropception DE generation device**. Single inner aqueous in oil emulsions are formed at flow focuser 1 (FF1), which then travel through the resistor. These are wrapped in outer aqueous sheath at flow focuser 2 (FF2) to become double emulsions and flow out the outlet channel.

**Figure S2: DE systematic characterization design and absolute flow rates.**

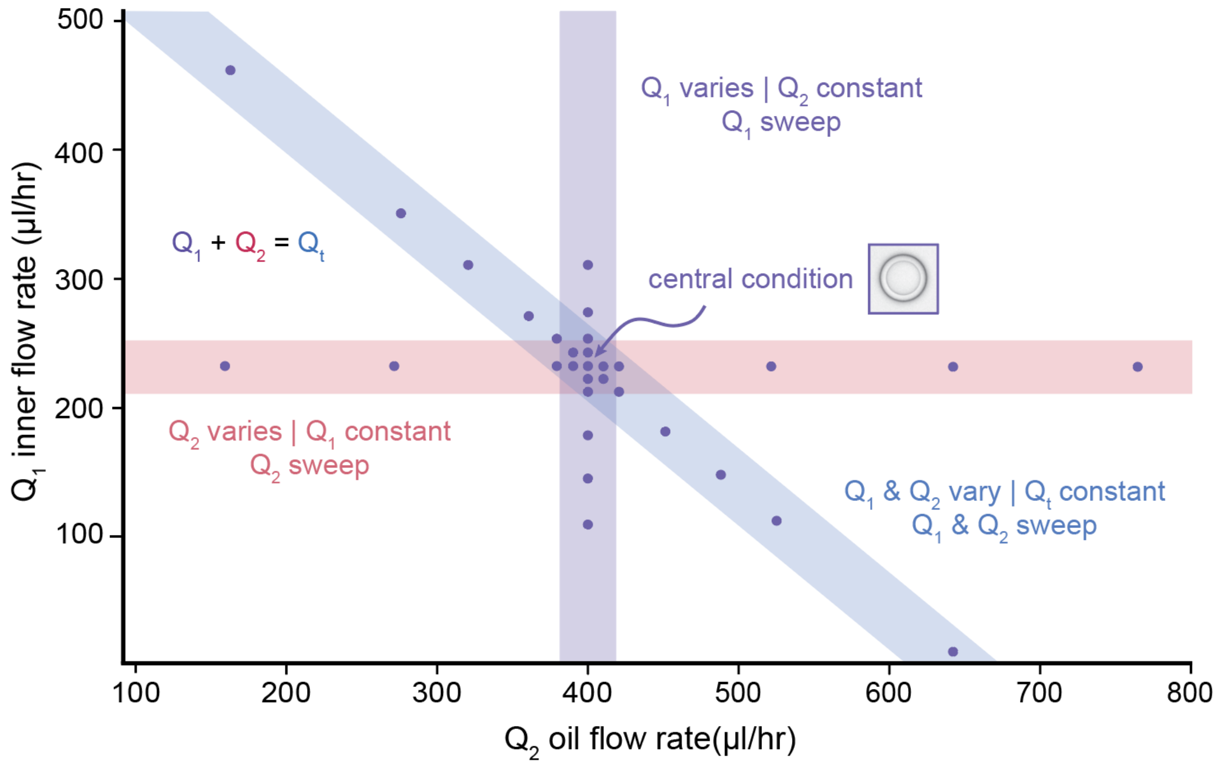

**Figure S2: DE systematic characterization design and absolute flow rates.** Q1 inner absolute flow rates *vs.* Q2 oil flow rates measured in this study in μl /hr. Red band indicates flow rates used to vary Q2 while keeping Q1 constant, purple band indicates flow rates used to vary Q1 while keeping Q2 constant, and blue band indicates flow rates used to vary both simultaneously while holding Qt = Q1 + Q2 constant. Central condition marks initial flow rate condition of ~0.60 core volume: shell volume. Flow rates shown here are for PBS + 1% Tween-20 buffer.

**Figure S3: Replicate experiment measuring DE diameters for PBS + 1% Tween-20.**

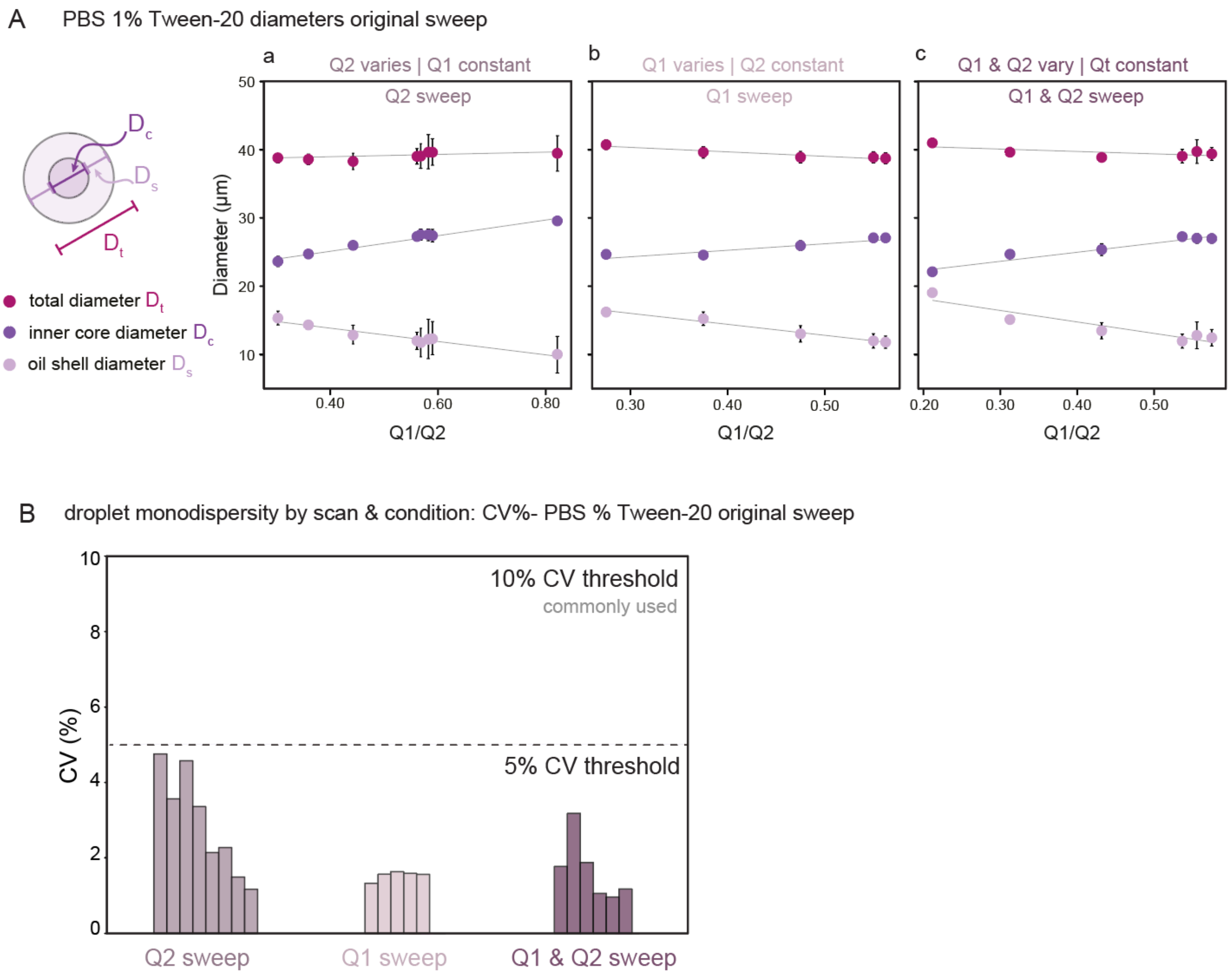

**Figure S3: Replicate experiment measuring DE diameters for PBS + 1% Tween-20** (performed using a new device in a different laboratory). **(A)** Measured inner core (dark purple), oil shell (light purple), and total (magenta) diameters for flow conditions varying oil flow rate (Q2) only (left), inner aqueous flow rate (Q1) only (middle), or simultaneously varying Q1 and Q2 (right). Markers indicate median, error bars represent standard deviation, and solid lines show a linear regression. **(B)** Measured diameter coefficient of variation (CV) (%) across droplets from each flow condition (droplet numbers for each condition are provided in Table S5).

**Figure S4: Measured DE droplet diameters and volumes with 6 inner aqueous buffers**

#### A. Droplet Diameter vs. Q1/Q2

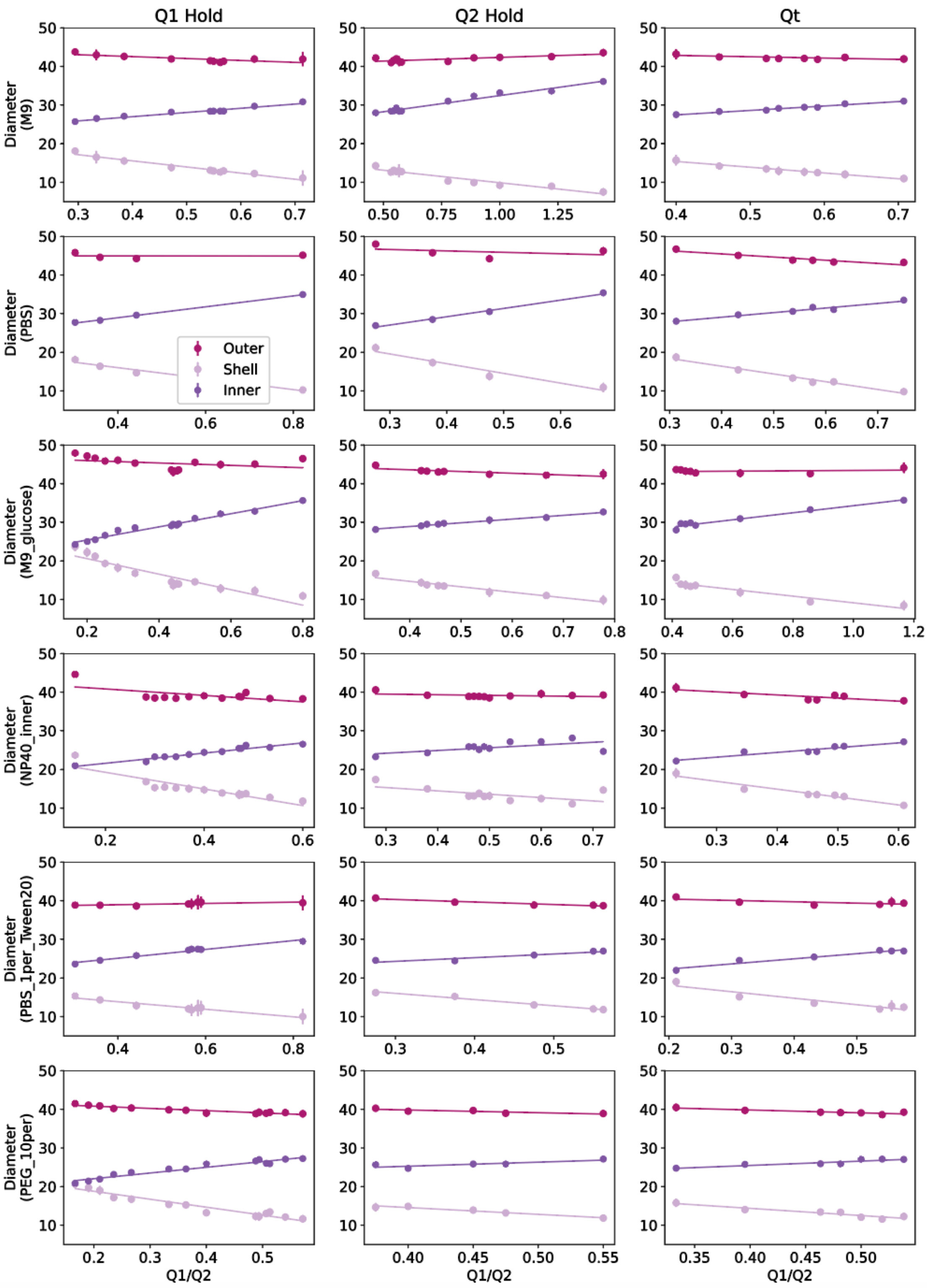

#### Figure S4: B. Droplet Volume vs. Q1/Q2

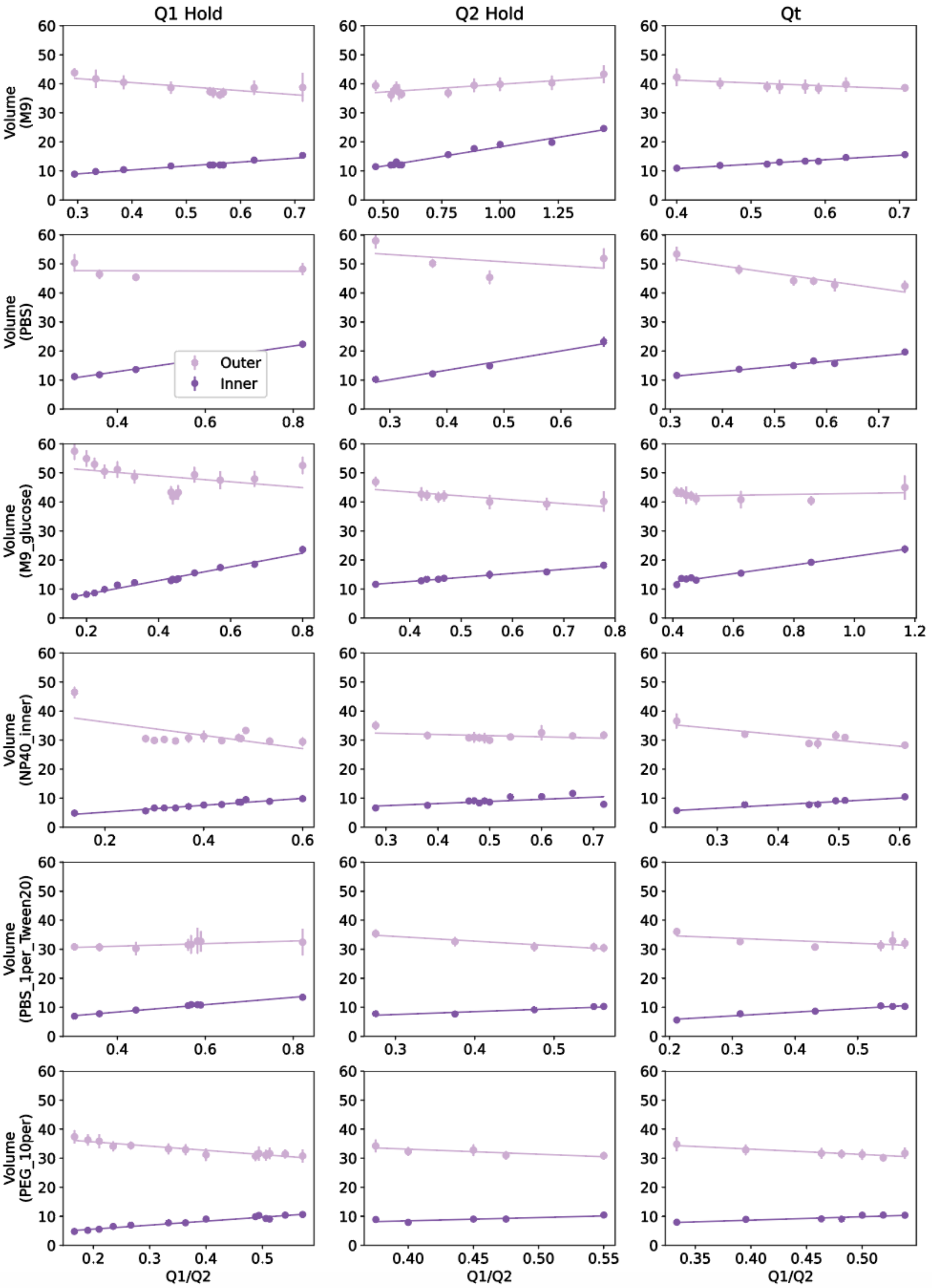

**Figure S4: C. Droplet Diameter vs. Q1/(Q1+Q2)**

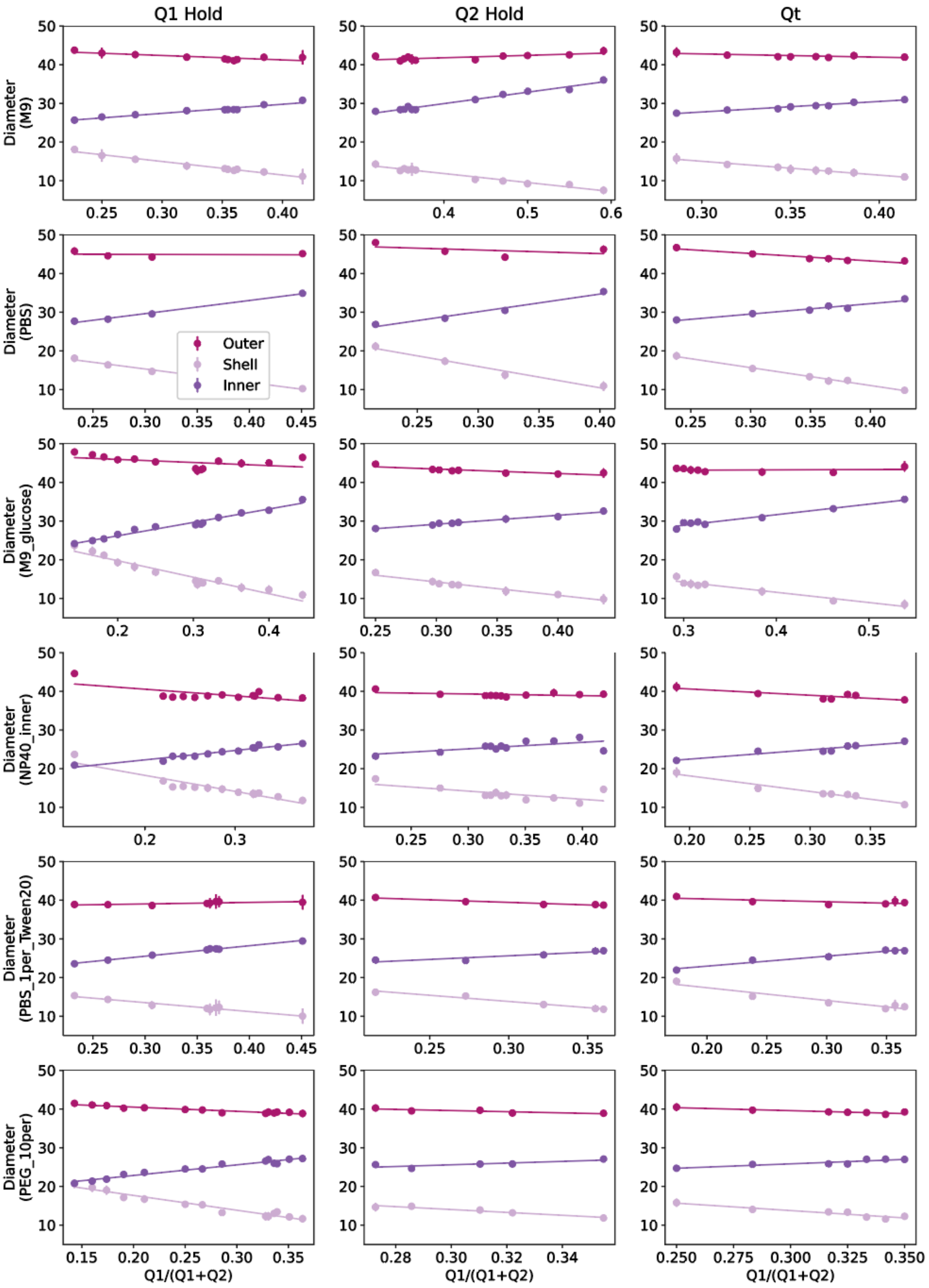

#### Figure S4: D. Droplet Volume vs. Q1/(Q1+Q2)

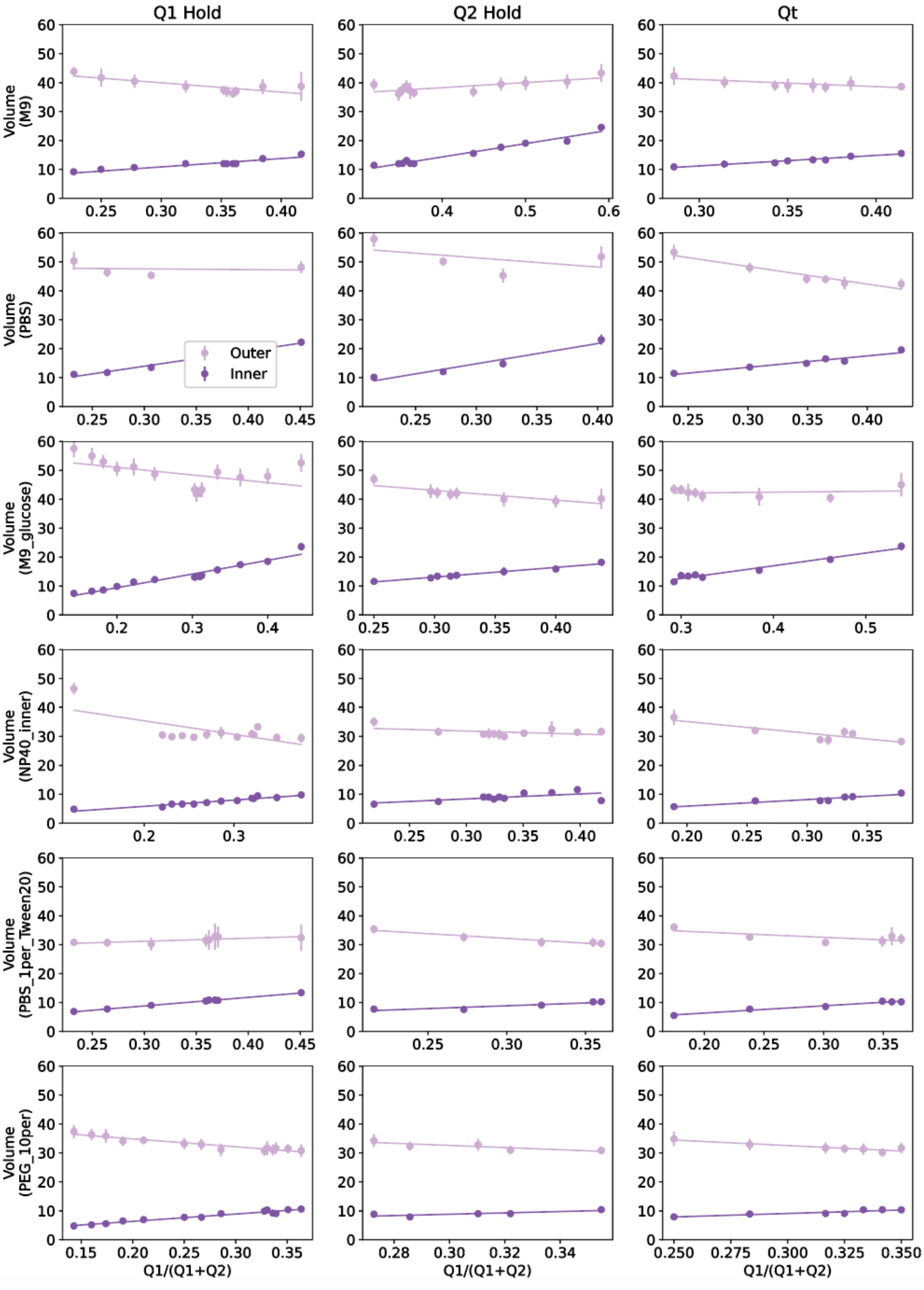

**Figure S4: Measured DE droplet diameters and volumes with 6 inner aqueous buffers.** Measured diameters/volumes for flow conditions varying oil flow rate (Q2) only (left), inner aqueous flow rate (Q1) only (middle), or simultaneously varying Q1 and Q2 with Qt = Q1+Q2 constant (right). Markers indicate median, error bars represent standard deviation, and solid lines show a linear regression. **(A)** Measured inner core (dark purple), oil shell (light purple), and total (magenta) diameter vs. Q1/Q2 flow rate ratio **(B)** Measured inner core (dark purple), and total outer (magenta) volume vs. Q1/Q2 flow rate ratio. **(C)** Measured inner core (dark purple), oil shell (light purple), and total (magenta) diameter vs. Q1/(Q1+Q2) flow rate ratio **(D)** Measured inner core (dark purple), and total outer (magenta) volume vs. Q1/(Q1+Q2) flow rate ratio.

**Figure S5: Measured Diameter Coefficient of variation** CV% across flow rates and solutions

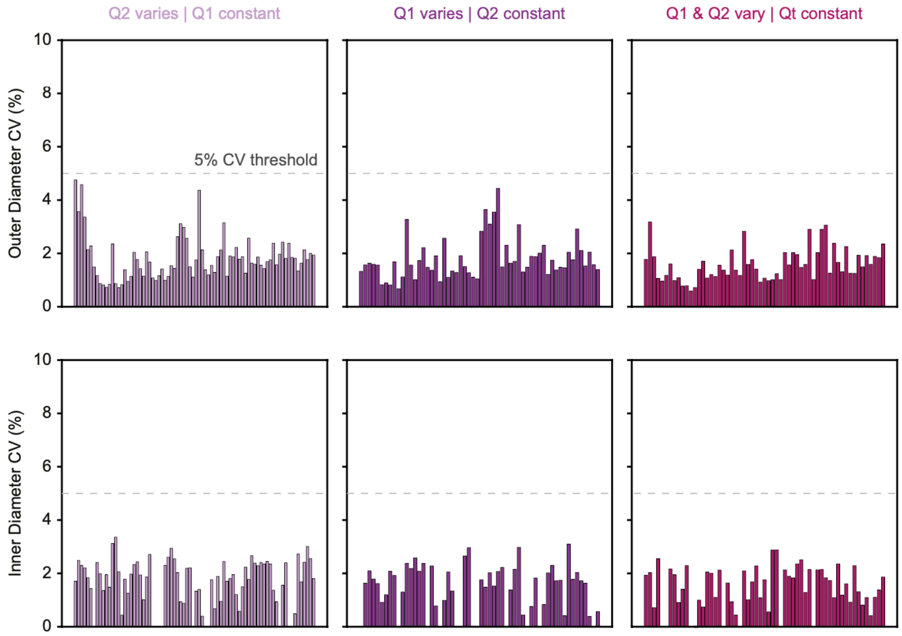

**Figure S5: Measured Diameter Coefficient of variation CV% across flow rates and solutions.** Coefficient of variation (CV) (%) of outer total diameters (top) and inner diameters (bottom) for 138 unique flow rate conditions containing 6 different inner buffers, for flow conditions varying oil flow rate (Q2) only (left), inner aqueous flow rate (Q1) only (middle), or simultaneously varying Q1 and Q2 (right) while holding Qt = Q1+Q2 constant. Each line represents CV% for one flow rate and solution composition combination; the number of droplets measured for each condition is given in Table S5.

#### Figure S6 Schematic illustrating Hydrophilic Lipophilic Balance (HLB) and potential impact on interfacial tension (IFT)

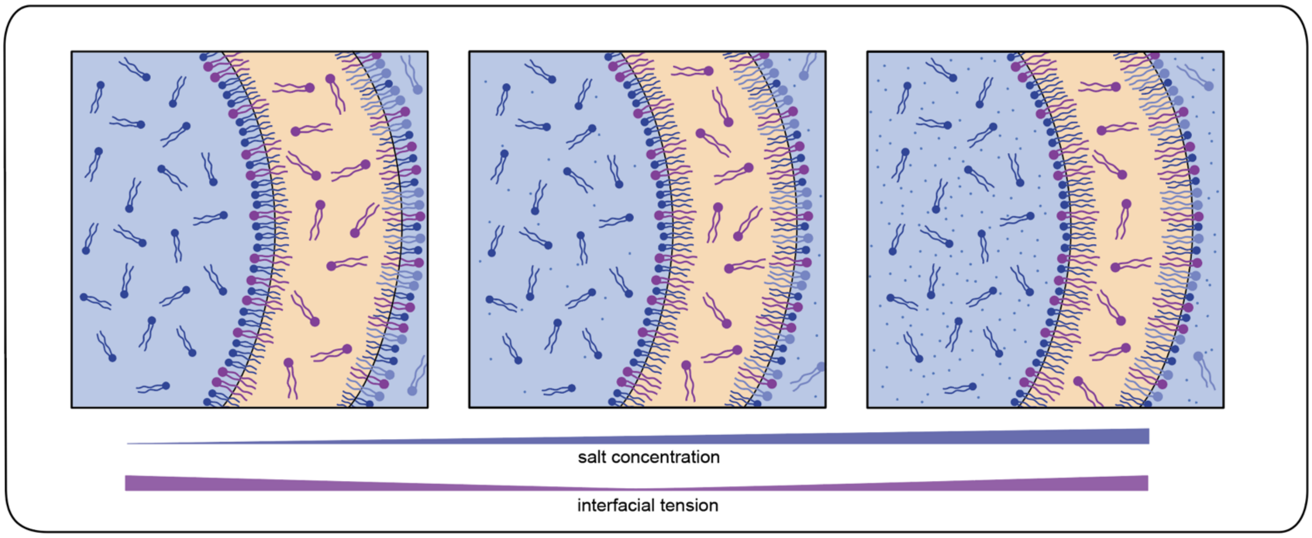

**Figure S6: Schematic illustrating Hydrophilic Lipophilic Balance (HLB) and potential impact on interfacial tension (IFT) resulting in nonlinear changes in IFT as a function of salt concentration.** Relative fraction of interface surfactant molecules in aqueous (blue) vs. oil (yellow) phase varies with increasing salt concentration. Low salt concentration (left) with interface surfactant molecules primarily in the aqueous phase and high IFT, intermediate salt concentration (center) with balanced surfactant molecules leading to minimum IFT, and high salt concentration (right) with surfactant molecules primarily in the oil phase and high IFT.

#### Figure S7: Pendant drop method to measure interfacial tension (IFT).

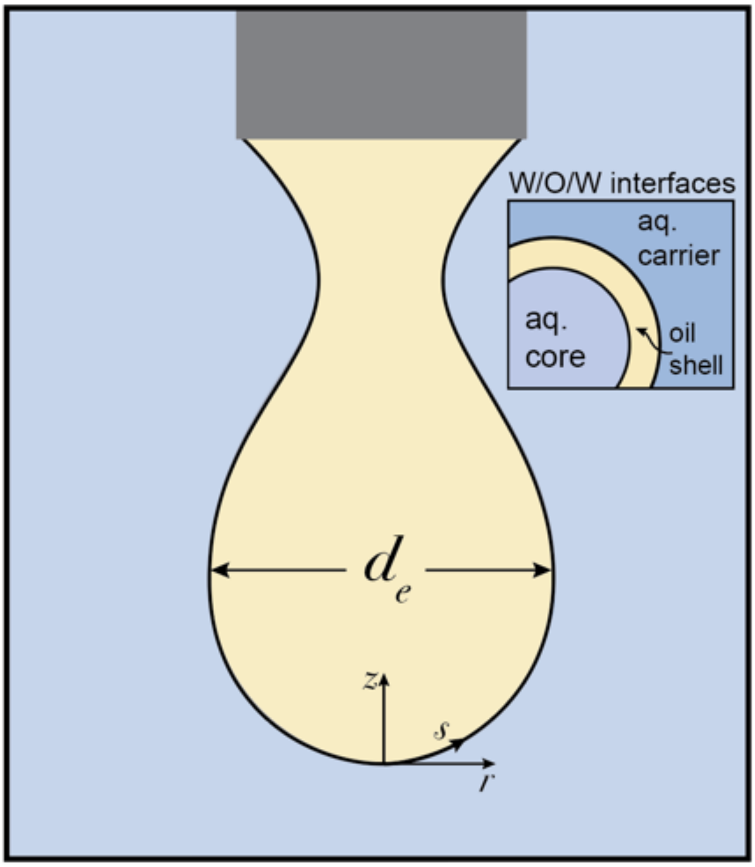

**Figure S7: Pendant drop method to measure interfacial tension (IFT).** Reverse phase pendant drop method was used, suspending an oil drop within an inner core aqueous solution and using drop shape analysis to calculate IFT. Here $z$, $s$, and $r$ mark coordinate system.

**Figure S8: Example images showing how varying flow rates alter droplet morphology and stability**

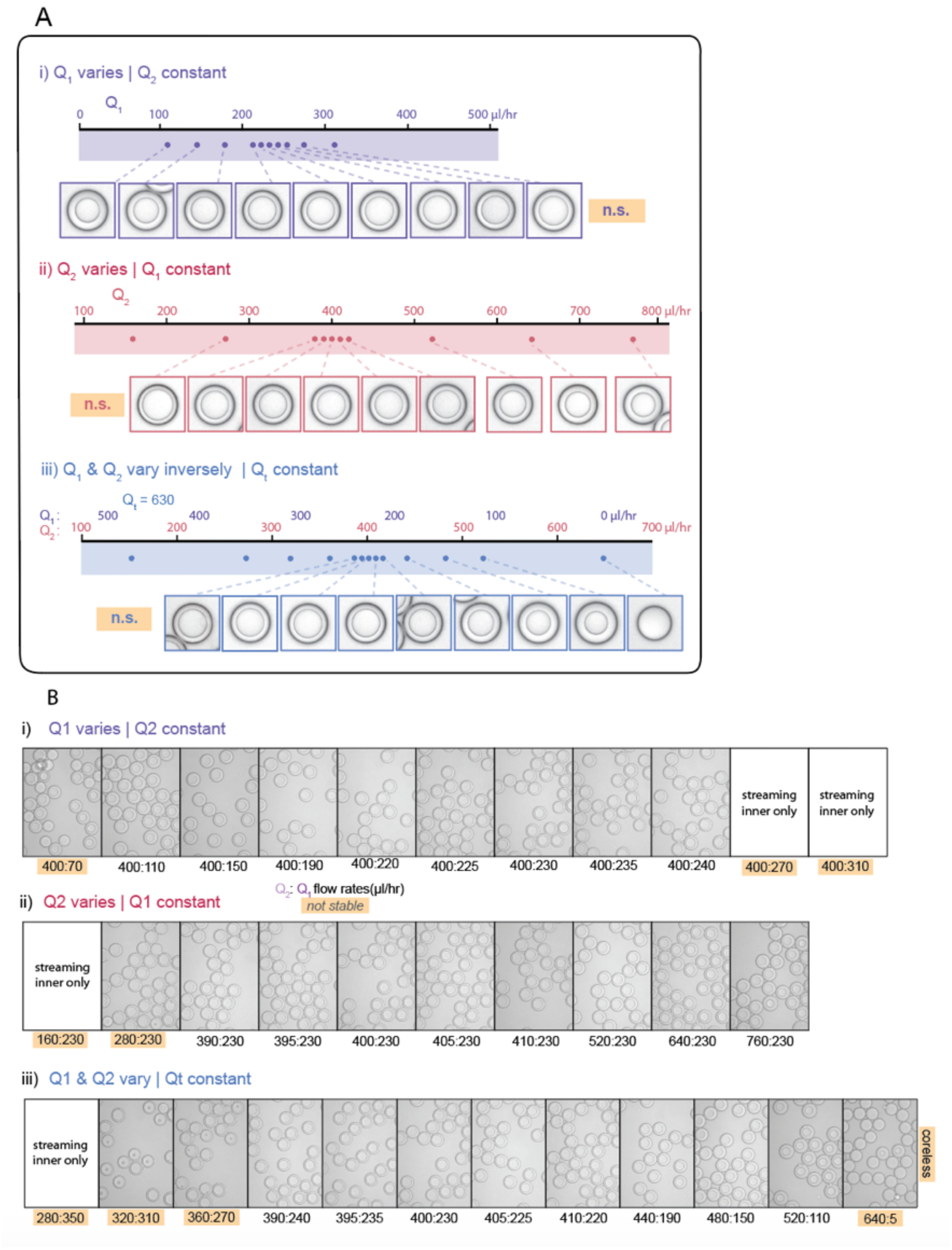

**Figure S8: Example images showing how varying flow rates alter droplet morphology and stability. (A)** Flow rates collected for flow conditions varying oil flow rate (Q2) only (top, purple), inner aqueous flow rate (Q1) only (middle, red), or simultaneously varying Q1 and Q2 with Qt=Q1+Q2 constant (bottom, blue), with representative images of a single droplet for stable conditions. Flow rates & images shown here are for PBS + 1% Tween-20 buffer droplets. **(B)** Representative microscopy images showing stability and morphology for flow conditions varying oil flow rate (Q2) only (top), inner aqueous flow rate (Q1) only (middle), or simultaneously varying Q1 and Q2 with Qt=Q1+Q2 constant (bottom). Flow rate conditions marked in orange result in non-ideal flow behavior and thus are classified as instabilities; flow rates and images here are PBS + 0.9% np40 inner buffer droplets.

#### Figure S9: Additional possible size scaling laws and comparisons between predictions and measurements

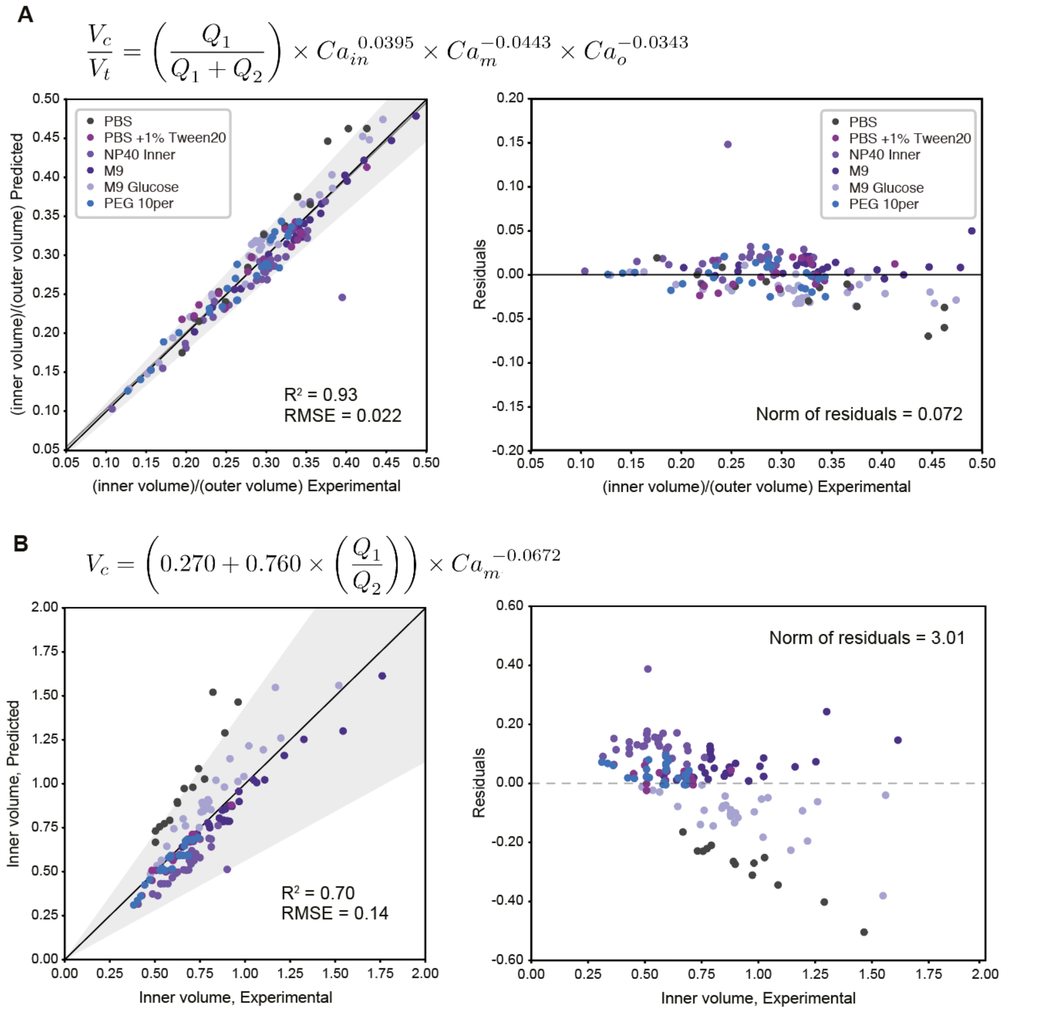

**Figure S9: Additional possible size scaling laws and comparisons between predictions and measurements, (Left)** Predicted vs experimental core volume: total volume ratio for 196 droplet conditions, separated by buffer composition. Dark line indicates 1:1 line; light line indicates linear regression; grey shading indicates confidence interval; $R^2$ and RMSE and model equation used are indicated. **(Right)** Calculated residuals of experimental results from model as a function of volume**. (A) Model C.** Droplet Core (inner): total (outer) volume ratio modelled with relevant solution properties (in the form of capillary numbers) and without an initial scalar explains 95% of observations within a 10.7% interval. **(B) Inner volume Model.** Droplet inner volume modelled with capillary numbers explains 95% of observations within a 31% interval.

#### Figure S10: Goodness of Fit for simplified mass conservation size scaling law

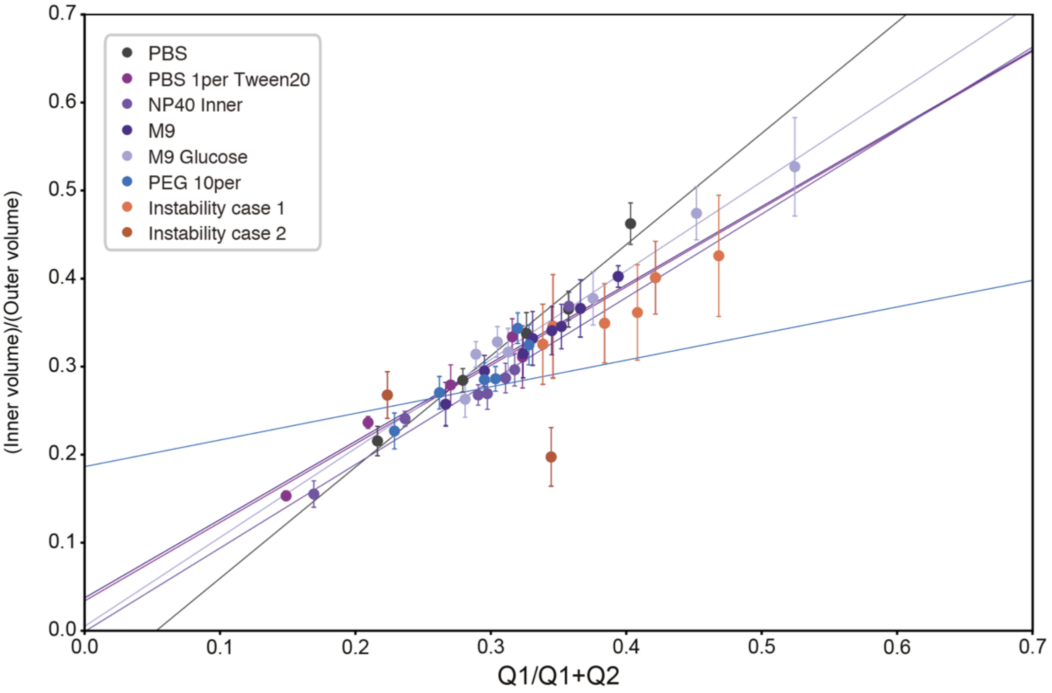

**Figure S10: Goodness of Fit for simplified mass conservation size scaling law.** Measured volume ratio vs ideal volume term (mass conservation only), showing deviation from the simplified size scaling model. Only Qt hold conditions are plotted. The two instability types are included in orange and red. Larger distance from model fit line indicates worse model fit. Markers indicate median, error bars represent standard deviation, and solid lines show model fit for each buffer composition.

#### Table S8: Goodness of fit comparison between volume ratio models (core volume: total volume $V_c$:$V_t$)

| Model | Slope | $R^2$ | RMSE | % deviation for 95% confidence | Residual norm |
|---|---|---|---|---|---|
| **A (volume ratio)** | 0.937 | 0.931 | 0.0209490 | +/- 10.7 | 0.0667069 |
| **B (volume ratio)** | 0.924 | 0.924 | 0.0219914 | +/- 11.6 | 0.0735105 |
| **C (volume ratio)** | 0.943 | 0.925 | 0.0218094 | +/- 10.7 | 0.0722985 |

#### Table S9: Goodness of fit for each trial

| Model | Trial | $R^2$ | RMSE | % deviation for 95% confidence | Residual norm |
|---|---|---|---|---|---|
| **A (volume ratio)** | PBS_1per_Tween20 | 0.944 | 0.0139596 | +/- 10.7 | 0.0037025 |
| **A (volume ratio)** | PBS | 0.903 | 0.0283926 | +/- 17.2 | 0.0112860 |
| **A (volume ratio)** | NP40_inner | 0.669 | 0.0328502 | +/- 11.3 | 0.0334532 |
| **A (volume ratio)** | PEG_10per | 0.948 | 0.0141316 | +/- 9.60 | 0.0051923 |
| **A (volume ratio)** | M9 | 0.981 | 0.0102821 | +/- 5.00 | 0.0033831 |
| **A (volume ratio)** | M9_glucose | 0.934 | 0.0234474 | +/- 12.3 | 0.0164934 |
| **B (volume ratio)** | PBS_1per_Tween20 | 0.944 | 0.0139596 | +/- 10.7 | 0.0037025 |
| **B (volume ratio)** | PBS | 0.903 | 0.0283926 | +/- 17.2 | 0.0112860 |
| **B (volume ratio)** | NP40_inner | 0.669 | 0.0328502 | +/- 11.3 | 0.0334532 |
| **B (volume ratio)** | PEG_10per | 0.948 | 0.0141316 | +/- 9.60 | 0.0051923 |
| **B (volume ratio)** | M9 | 0.981 | 0.0102821 | +/- 5.00 | 0.0033831 |
| **B (volume ratio)** | M9_glucose | 0.934 | 0.0234474 | +/- 12.3 | 0.0164934 |
| **C (volume ratio)** | PBS_1per_Tween20 | 0.949 | 0.0132644 | +/- 12.2 | 0.0033429 |
| **C (volume ratio)** | PBS | 0.883 | 0.0310915 | +/- 18.6 | 0.0135336 |
| **C (volume ratio)** | NP40_inner | 0.692 | 0.0316759 | +/- 10.7 | 0.0311043 |
| **C (volume ratio)** | PEG_10per | 0.948 | 0.0141471 | +/- 10.1 | 0.0052037 |
| **C (volume ratio)** | M9 | 0.968 | 0.0135780 | +/- 6.30 | 0.0058996 |
| **C (volume ratio)** | M9_glucose | 0.947 | 0.0209877 | +/- 11.4 | 0.0132145 |
| **Inner volume** | PBS_1per_Tween20 | 0.925 | 0.0321632 | +/- 14.9 | 0.0196549 |
| **Inner volume** | PBS | -0.734 | 0.3410208 | +/- 85.2 | 1.6281327 |
| **Inner volume** | NP40_inner | -0.976 | 0.1428169 | +/- 29.5 | 0.6322966 |
| **Inner volume** | PEG_10per | 0.820 | 0.0463829 | +/- 16.8 | 0.0559598 |
| **Inner volume** | M9 | 0.849 | 0.0805129 | +/- 13.9 | 0.2074346 |
| **Inner volume** | M9_glucose | 0.746 | 0.1247599 | +/- 24.8 | 0.4669510 |

#### Figure S11: Stability and instability types in DE droplet formation

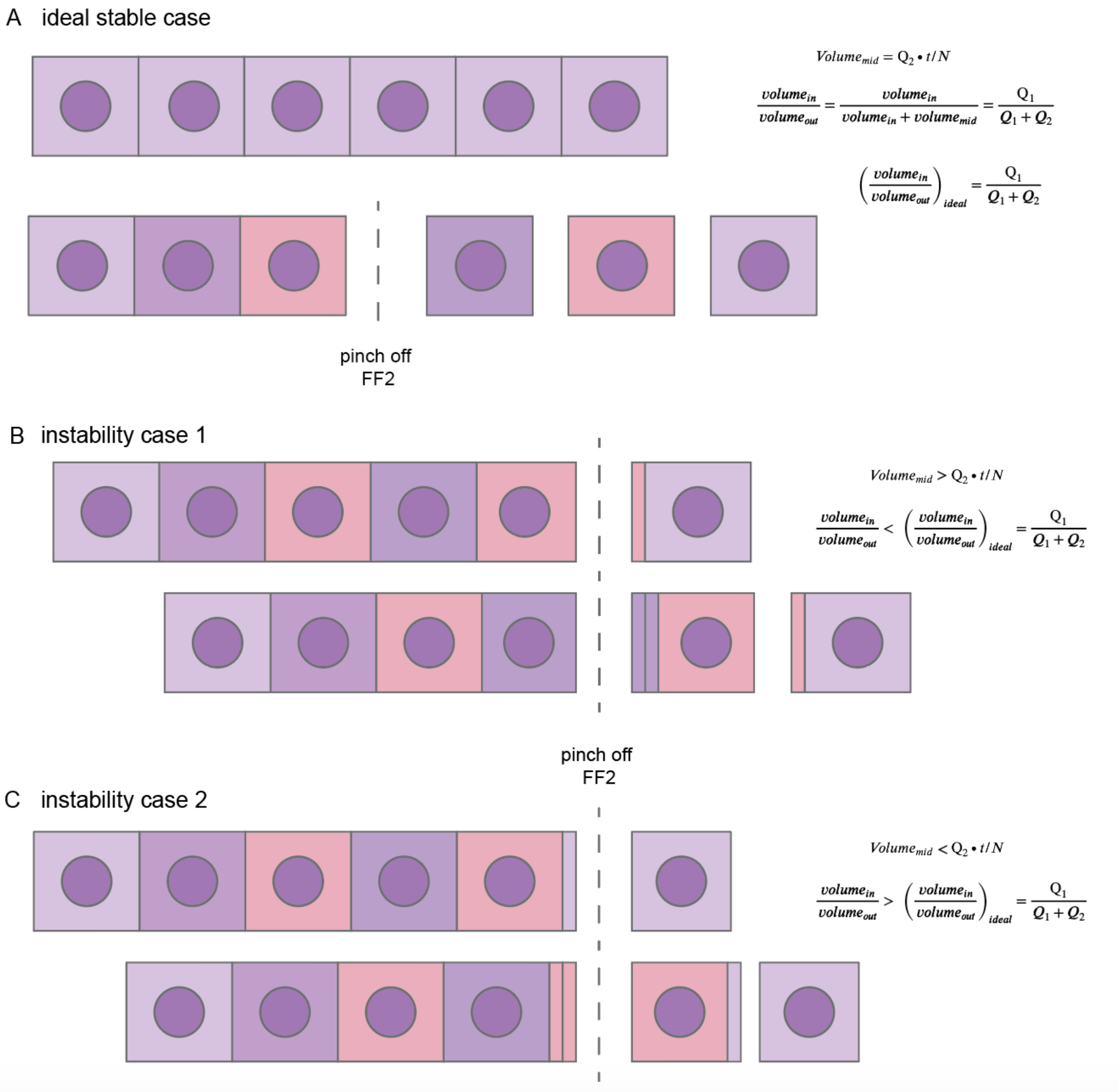

**Figure S11: Stability and instability types in DE droplet formation. (A)** Ideal stable case of droplet formation, where FF2 pinch off occurs with equal spacing so that $volume_{mid}$ equals time / number of droplets. Matching periodicity of oil and inner phases with outer sheath phase regulates FF2 pinch off and generates stable droplets, with inner:total volume ratio equal to Q1/Q1+Q2, as expected by mass conservation**.** **(B**) Case 1 type of droplet formation instability, where FF2 pinch off is later and $volume_{mid}$ is larger than the ideal case, resulting in double core droplets. **(C)** Case 2 type of droplet formation instability, where FF2 pinch off is earlier and $volume_{mid}$ is smaller than the ideal case resulting in a subpopulation of coreless droplets.

#### Figure S12: Representative microscopy images of DE droplets containing fluorescent E. coli.

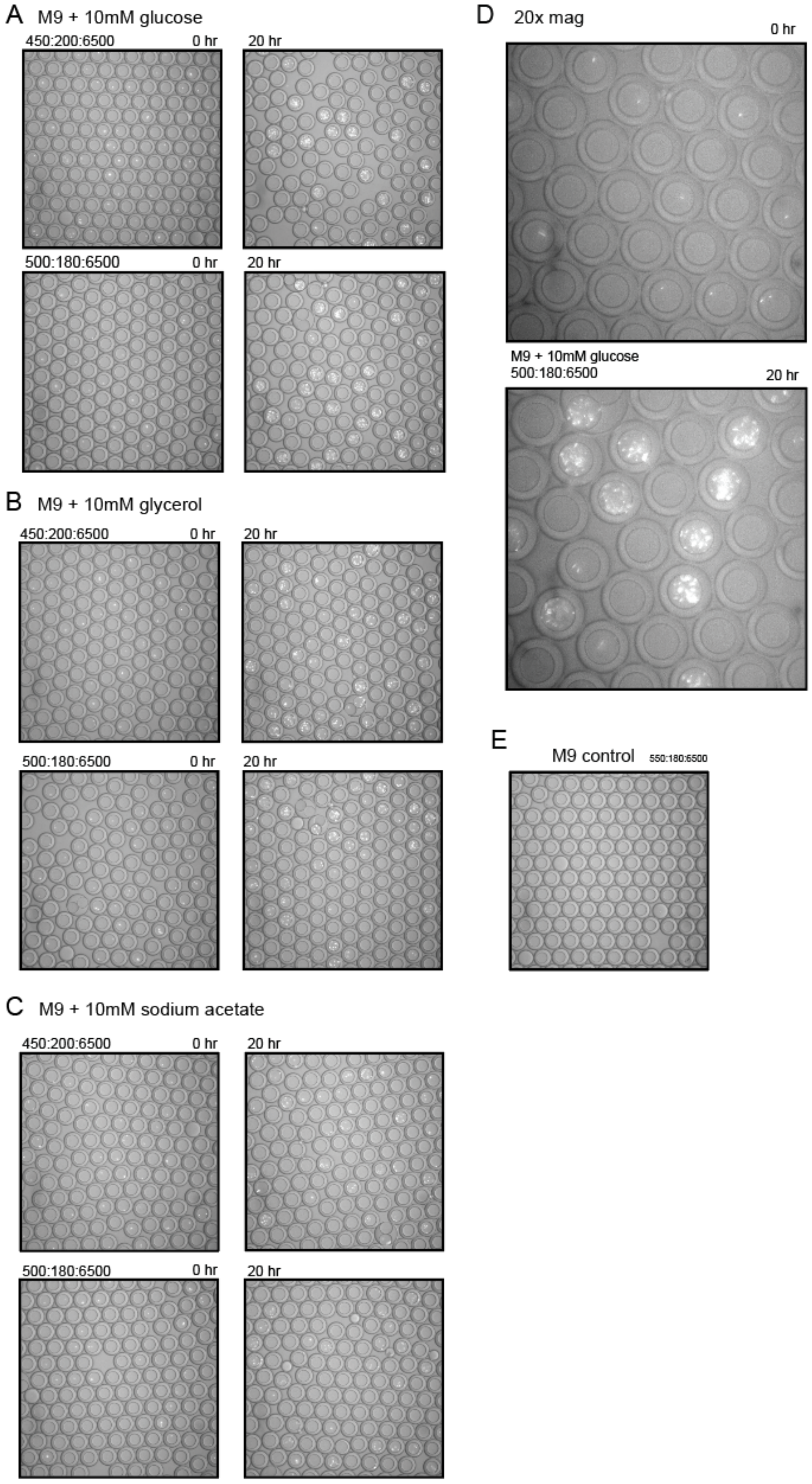

**Figure S12: Representative microscopy images of DE droplets containing fluorescent E. coli. (A-C)** Representative 10x microscopy images of E. coli growth in droplets at 0hr (left) and 20 hr (right) for two flow rate conditions (450:200:6500 top, 500:180:6500 bottom, Q2:Q1:Q3) for each of 3 carbon sources: **(A)** M9 + 10mM glucose, **(B)** M9 + 10mM glycerol, and **(C)** M9 + 10mM sodium acetate. **(D)** Representative microscopy images at 20x of E coli growth at 0hr and 20 hr for one flow rate and carbon source combination. **(E)** Microscopy image of control empty droplets.

**Table S1:** Extended interfacial and bulk fluid parameters

| Surfactant System | Phase | Components | Application | Density (kg/m³ ) | Dynamic viscosity (mPa s) | Interfacial tension with Oil (mN/m) |
|---|---|---|---|---|---|---|
| PBS | Inner | PBS | Control | 1005.58 | 0.931 | 0.543 |
| | Middle | HFE-7500 + 2.2% ionic Krytox | Oxygen permeability | 1619.72 | 1.613 | n/a |
| | Outer | PBS + 1% Tween-20 + 2% Pluronix-F68 | | 1007.97 | 1.303 | 0.318 |
| PBS + 1% Tween-20 | Inner | PBS + 1% Tween-20 | Cell lysis buffer | 1006.48 | 0.988 | 0.319 |
| | Middle | HFE-7500 + 2.2% ionic Krytox | Oxygen permeability | 1619.72 | 1.613 | n/a |
| | Outer | PBS + 1% Tween-20 + 2% Pluronix-F68 | | 1007.97 | 1.303 | 0.318 |
| PBS + 0.9% NP40 | Inner | PBS + 0.9% NP40 inner | Cell lysis buffer | 1006.07 | 1.003 | 1.41 |
| | Middle | HFE-7500 + 2.2% ionic Krytox | Oxygen permeability | 1619.72 | 1.613 | n/a |
| | Outer | PBS + 1% Tween-20 + 2% Pluronix-F68 | | 1007.97 | 1.303 | 0.318 |
| M9 bacterial media | Inner | M9 salts | Bacterial growth media | 1013.0 | 0.861 | 12.84 |
| | Middle | HFE-7500 + 2.2% ionic Krytox | Oxygen permeability | 1619.72 | 1.613 | n/a |
| | Outer | M9 salts + 2% Pluronix-F68 | | 1013.4 | 1.412 | 0.5220 |
| M9 + 25mM glucose | Inner | M9 salts + 25mM glucose | Bacterial growth media with Carbon source | 1017.5 | 0.967 | 11.600 |
| | Middle | HFE-7500 + 2.2% ionic Krytox | Oxygen permeability | 1619.72 | 1.613 | n/a |
| | Outer | M9 salts + 25mM glucose + 2% Pluronix-F68 | | 1017.9 | 1.563 | 0.4580 |
| PBS + 10% PEG 6000 mw | Inner | PBS + 10% PEG 6000 mw | Polymer | 1013.7 | 3.431 | 0.4613 |
| | Middle | HFE-7500 + 2.2% ionic Krytox | Oxygen permeability | 1619.72 | 1.613 | n/a |
| | Outer | PBS + 10% PEG 6000 mw + 2% Pluronix-F68 | | 1014.1 | 6.395 | 0.4550 |
| **Benchmarks:** | | | | | | |
| MilliQ H20 in custom oil | | MilliQ H20 | | | | 17.567 |
| MilliQ H20 in HFE-7500 | | MilliQ H20 | | | | 38.620 |
| M9 Inner in HFE-7500 | | M9 salts | | | | 44.799 |
| PBS in HFE-7500 | | PBS | | | | 40.587 |
| **Salt concentration sweep** | | | | | **Salt concentration mM** | **IFT w/ custom oil** |
| PBS 0.03125x | | PBS, MilliQ H20 | Salt concentration sweep | | 1.18 | 4.832 |
| PBS 0.125x | | PBS, MilliQ H20 | Salt concentration sweep | | 4.74 | 4.199 |
| PBS 0.25x | | PBS, MilliQ H20 | Salt concentration sweep | | 9.48 | 2.576 |
| PBS 0.5x | | PBS, MilliQ H20 | Salt concentration sweep | | 18.95 | 0.907 |
| PBS | | PBS | Salt concentration sweep | | 37.9 | 0.543 |
| PBS 5x | | PBS 10x, MilliQ H20 | Salt concentration sweep | | 189.5 | 0.682 |

| | | | | | | |
|---|---|---|---|---|---|---|
| PBS 10x | | PBS 10x | Salt concentration sweep | | 379 | 11.477 |
| M9 inner | | M9 salts | Salt concentration sweep | | 24.27 | 12.556 |
| **E coli solutions** | | | | | | |
| M9 + 10mM glucose | Inner | M9 salts, glucose | | | | 12.454 |
| M9 + 10mM sodium acetate | Inner | M9 salts, sodium acetate | | | | 13.869 |
| M9 + 10mM glycerol | Inner | M9 salts, glycerol | | | | 10.604 |

#### Table S2: Salt concentration sweep

| Salt concentration sweep | | Components | Application | | Salt concentration mM | IFT w/ custom oil |
|---|---|---|---|---|---|---|
| PBS 0.03125x | | PBS, MilliQ H20 | Salt concentration sweep | | 1.18 | 4.832 |
| PBS 0.125x | | PBS, MilliQ H20 | Salt concentration sweep | | 4.74 | 4.199 |
| PBS 0.25x | | PBS, MilliQ H20 | Salt concentration sweep | | 9.48 | 2.576 |
| PBS 0.5x | | PBS, MilliQ H20 | Salt concentration sweep | | 18.95 | 0.907 |
| PBS | | PBS | Salt concentration sweep | | 37.9 | 0.543 |
| PBS 5x | | PBS 10x, MilliQ H20 | Salt concentration sweep | | 189.5 | 0.682 |
| PBS 10x | | PBS 10x | Salt concentration sweep | | 379 | 11.477 |
| M9 inner | | M9 salts | Salt concentration sweep | | 24.27 | 12.556 |

#### Table S3: E. coli solutions

| E coli solutions | Phase | Components | | | | Interfacial tension with Oil (mN/m) |
|---|---|---|---|---|---|---|
| M9 + 10mM glucose | Inner | M9 salts, glucose | | | | 12.454 |
| M9 + 10mM sodium acetate | Inner | M9 salts, sodium acetate | | | | 13.869 |
| M9 + 10mM glycerol | Inner | M9 salts, glycerol | | | | 10.604 |

#### Table S4: PBS + 1% Tween-20 Original and replicate

| condition_num | trial | rho_in | sigma_in_oil | mu_in | rho_oil | mu_oil | rho_out | sigma_out_oil | mu_out | sweep_condition | Q1 | Q2 | Qt | Q3 | outer_diam_mean | outer_diam_std | inner_diam_mean | inner_diam_std | shell_diam_mean | shell_diam_std | outer_vol_mean | outer_vol_std | inner_vol_mean | inner_vol_std | shell_vol_mean | shell_vol_std | core_total_ratio | shell_total_ratio | core_shell_ratio | n_droplets |
|---|---|---|---|---|---|---|---|---|---|---|---|---|---|---|---|---|---|---|---|---|---|---|---|---|---|---|---|---|---|---|
| **1** | PBS_1per_Tween20 | 1006.478 | 0.000303 | 0.0009875 | 1619.72 | 0.0016135 | 1007.97 | 0.000322 | 0.0013034 | Q1 hold | 230 | 280 | 510 | 6500 | 39.461 | 1.877 | 29.44 | 0.503 | 10.021 | 1.945 | 32.382 | 4.528 | 13.371 | 0.666 | 19.011 | 4.591 | 0.413 | 0.587 | 0.703 | 28 |
| **2** | PBS_1per_Tween20 | 1006.478 | 0.000303 | 0.0009875 | 1619.72 | 0.0016135 | 1007.97 | 0.000322 | 0.0013034 | Q1 hold | 230 | 390 | 620 | 6500 | 39.617 | 1.413 | 27.314 | 0.679 | 12.303 | 1.76 | 32.681 | 3.513 | 10.69 | 0.8 | 21.991 | 3.849 | 0.327 | 0.673 | 0.486 | 92 |
| **3** | PBS_1per_Tween20 | 1006.478 | 0.000303 | 0.0009875 | 1619.72 | 0.0016135 | 1007.97 | 0.000322 | 0.0013034 | Q1 hold | 230 | 395 | 625 | 6500 | 39.639 | 1.814 | 27.42 | 0.632 | 12.219 | 2.085 | 32.814 | 4.438 | 10.812 | 0.755 | 22.002 | 4.701 | 0.329 | 0.671 | 0.491 | 86 |
| **4** | PBS_1per_Tween20 | 1006.478 | 0.000303 | 0.0009875 | 1619.72 | 0.0016135 | 1007.97 | 0.000322 | 0.0013034 | Q2 hold | 110 | 400 | 510 | 6500 | 40.715 | 0.54 | 24.51 | 0 | 16.205 | 0 | 35.357 | 1.4 | 7.71 | 0 | 27.648 | 0 | 0.218 | 0.782 | 0.279 | 89 |
| **5** | PBS_1per_Tween20 | 1006.478 | 0.000303 | 0.0009875 | 1619.72 | 0.0016135 | 1007.97 | 0.000322 | 0.0013034 | Q2 hold | 150 | 400 | 550 | 6500 | 39.614 | 0.622 | 24.376 | 0.398 | 15.238 | 0.736 | 32.573 | 1.581 | 7.589 | 0.356 | 24.983 | 1.618 | 0.233 | 0.767 | 0.304 | 48 |
| **6** | PBS_1per_Tween20 | 1006.478 | 0.000303 | 0.0009875 | 1619.72 | 0.0016135 | 1007.97 | 0.000322 | 0.0013034 | Q2 hold | 190 | 400 | 590 | 6500 | 38.885 | 0.636 | 25.867 | 0.542 | 13.018 | 0.881 | 30.81 | 1.514 | 9.074 | 0.574 | 21.735 | 1.682 | 0.295 | 0.705 | 0.417 | 96 |
| **7** | PBS_1per_Tween20 | 1006.478 | 0.000303 | 0.0009875 | 1619.72 | 0.0016135 | 1007.97 | 0.000322 | 0.0013034 | Q2 hold | 220 | 400 | 620 | 6500 | 38.865 | 0.619 | 26.877 | 0.481 | 11.988 | 0.779 | 30.762 | 1.485 | 10.176 | 0.529 | 20.586 | 1.57 | 0.331 | 0.669 | 0.494 | 97 |
| **8** | PBS_1per_Tween20 | 1006.478 | 0.000303 | 0.0009875 | 1619.72 | 0.0016135 | 1007.97 | 0.000322 | 0.0013034 | Q2 hold | 225 | 400 | 625 | 6500 | 38.725 | 0.606 | 26.923 | 0.436 | 11.802 | 0.665 | 30.429 | 1.372 | 10.226 | 0.479 | 20.203 | 1.356 | 0.336 | 0.664 | 0.506 | 85 |
| **9** | PBS_1per_Tween20 | 1006.478 | 0.000303 | 0.0009875 | 1619.72 | 0.0016135 | 1007.97 | 0.000322 | 0.0013034 | Qt | 230 | 400 | 630 | 6500 | 39.371 | 0.699 | 26.923 | 0.519 | 12.448 | 0.887 | 31.985 | 1.718 | 10.229 | 0.58 | 21.756 | 1.833 | 0.32 | 0.68 | 0.47 | 85 |
| **10** | PBS_1per_Tween20 | 1006.478 | 0.000303 | 0.0009875 | 1619.72 | 0.0016135 | 1007.97 | 0.000322 | 0.0013034 | Qt | 225 | 405 | 630 | 6500 | 39.716 | 1.264 | 26.924 | 0.546 | 12.793 | 1.426 | 32.901 | 3.173 | 10.231 | 0.612 | 22.67 | 3.287 | 0.311 | 0.689 | 0.451 | 93 |

| | | | | | | | | | | | | | | | | | | | | | | | | | | | | | | |
|---|---|---|---|---|---|---|---|---|---|---|---|---|---|---|---|---|---|---|---|---|---|---|---|---|---|---|---|---|---|---|
| **11** | PBS_1per_Tween20 | 1006.478 | 0.000303 | 0.009875 | 1619.72 | 0.016135 | 1007.97 | 0.000322 | 0.013034 | Q1 hold | 230 | 405 | 635 | 6500 | 39.189 | 1.318 | 27.442 | 0.606 | 11.748 | 1.521 | 31.621 | 3.257 | 10.836 | 0.729 | 20.785 | 3.425 | 0.343 | 0.657 | 0.521 | 99 |
| **12** | PBS_1per_Tween20 | 1006.478 | 0.000303 | 0.009875 | 1619.72 | 0.016135 | 1007.97 | 0.000322 | 0.013034 | Qt | 220 | 410 | 630 | 6500 | 39.041 | 0.732 | 27.09 | 0.194 | 11.951 | 0.756 | 31.191 | 1.813 | 10.411 | 0.225 | 20.78 | 1.826 | 0.334 | 0.666 | 0.501 | 89 |
| **13** | PBS_1per_Tween20 | 1006.478 | 0.000303 | 0.009875 | 1619.72 | 0.016135 | 1007.97 | 0.000322 | 0.013034 | Q1 hold | 230 | 410 | 640 | 6500 | 39.157 | 0.839 | 27.155 | 0.5 | 12.002 | 0.925 | 31.48 | 2.059 | 10.495 | 0.586 | 20.984 | 2.06 | 0.333 | 0.667 | 0.5 | 99 |
| **14** | PBS_1per_Tween20 | 1006.478 | 0.000303 | 0.009875 | 1619.72 | 0.016135 | 1007.97 | 0.000322 | 0.013034 | Qt | 190 | 440 | 630 | 6500 | 38.858 | 0.412 | 25.384 | 0.648 | 13.475 | 0.872 | 30.732 | 0.983 | 8.58 | 0.644 | 22.152 | 1.337 | 0.279 | 0.721 | 0.387 | 127 |
| **15** | PBS_1per_Tween20 | 1006.478 | 0.000303 | 0.009875 | 1619.72 | 0.016135 | 1007.97 | 0.000322 | 0.013034 | Qt | 150 | 480 | 630 | 6500 | 39.629 | 0.382 | 24.51 | 0 | 15.119 | 0 | 32.595 | 0.937 | 7.71 | 0 | 24.885 | 0 | 0.237 | 0.763 | 0.31 | 100 |
| **16** | PBS_1per_Tween20 | 1006.478 | 0.000303 | 0.009875 | 1619.72 | 0.016135 | 1007.97 | 0.000322 | 0.013034 | Qt | 110 | 520 | 630 | 6500 | 40.98 | 0.482 | 21.93 | 0 | 19.05 | 0 | 36.049 | 1.258 | 5.522 | 0 | 30.527 | 0 | 0.153 | 0.847 | 0.181 | 86 |
| **17** | PBS_1per_Tween20 | 1006.478 | 0.000303 | 0.009875 | 1619.72 | 0.016135 | 1007.97 | 0.000322 | 0.013034 | Q1 hold | 230 | 520 | 750 | 6500 | 38.628 | 0.879 | 25.785 | 0.37 | 12.843 | 1.008 | 30.227 | 2.226 | 8.982 | 0.385 | 21.245 | 2.313 | 0.297 | 0.703 | 0.423 | 86 |
| **18** | PBS_1per_Tween20 | 1006.478 | 0.000303 | 0.009875 | 1619.72 | 0.016135 | 1007.97 | 0.000322 | 0.013034 | Q1 hold | 230 | 640 | 870 | 6500 | 38.837 | 0.58 | 24.51 | 0 | 14.327 | 0 | 30.691 | 1.382 | 7.71 | 0 | 22.982 | 0 | 0.251 | 0.749 | 0.335 | 90 |
| **19** | PBS_1per_Tween20 | 1006.478 | 0.000303 | 0.009875 | 1619.72 | 0.016135 | 1007.97 | 0.000322 | 0.013034 | Q1 hold | 230 | 760 | 990 | 6500 | 38.891 | 0.455 | 23.555 | 0.568 | 15.336 | 0.749 | 30.813 | 1.087 | 6.855 | 0.509 | 23.958 | 1.227 | 0.222 | 0.778 | 0.286 | 104 |
| **20** | PBS_1per_Tween20_replicate | 1006.478 | 0.000320 | 0.009875 | 1619.72 | 0.016135 | 1007.97 | 0.000328 | 0.013034 | Central condition | 230 | 400 | 630 | 6500 | 40.221 | 0.315 | 28.783 | 0.602 | 11.438 | 0.613 | 34.075 | 0.805 | 12.502 | 0.796 | 21.573 | 0.997 | 0.367 | 0.633 | 0.58 | 80 |
| **21** | PBS_1per_Tween20_replicate | 1006.478 | 0.000320 | 0.009875 | 1619.72 | 0.016135 | 1007.97 | 0.000328 | 0.013034 | Q2 hold | 220 | 400 | 620 | 6500 | 40.231 | 0.332 | 28.326 | 0.26 | 11.905 | 0.389 | 34.102 | 0.845 | 11.903 | 0.314 | 22.199 | 0.855 | 0.349 | 0.651 | 0.536 | 72 |
| **22** | PBS_1per_Tween20_replicate | 1006. | 0.000320 | 0.009875 | 1619 | 0.016135 | 1007 | 0.000328 | 0.013034 | Q2 hold | 225 | 400 | 625 | 65 | 40.162 | 0.361 | 28.372 | 0.34 | 11.79 | 0.471 | 33.928 | 0.922 | 11.963 | 0.43 | 21.964 | 0.979 | 0.353 | 0.647 | 0.545 | 159 |

| | | | | | | | | | | | | | | | | | | | | | | | | | | | | | | |
|---|---|---|---|---|---|---|---|---|---|---|---|---|---|---|---|---|---|---|---|---|---|---|---|---|---|---|---|---|---|---|
| | | 478 | | | .72 | | .97 | | | | | | | 00 | | | | | | | | | | | | | | | | |
| **23** | PBS_1per_Tween20_replicate | 1006.478 | 0.000320 | 0.009875 | 1619.72 | 0.016135 | 1007.97 | 0.000328 | 0.013034 | Q2 hold | 235 | 400 | 635 | 6500 | 40.228 | 0.329 | 28.776 | 0.599 | 11.452 | 0.646 | 34.093 | 0.837 | 12.492 | 0.793 | 21.601 | 1.08 | 0.366 | 0.634 | 0.578 | 75 |
| **24** | PBS_1per_Tween20_replicate | 1006.478 | 0.000320 | 0.009875 | 1619.72 | 0.016135 | 1007.97 | 0.000328 | 0.013034 | Q2 hold | 240 | 400 | 640 | 6500 | 40.184 | 0.675 | 28.688 | 0.552 | 11.496 | 0.786 | 34.004 | 1.734 | 12.377 | 0.731 | 21.627 | 1.747 | 0.364 | 0.636 | 0.572 | 138 |
| **25** | PBS_1per_Tween20_replicate | 1006.478 | 0.000320 | 0.009875 | 1619.72 | 0.016135 | 1007.97 | 0.000328 | 0.013034 | Q1 hold | 230 | 390 | 620 | 6500 | 40.238 | 0.351 | 28.711 | 0.571 | 11.527 | 0.697 | 34.12 | 0.892 | 12.406 | 0.755 | 21.714 | 1.221 | 0.364 | 0.636 | 0.571 | 39 |
| **26** | PBS_1per_Tween20_replicate | 1006.478 | 0.000320 | 0.009875 | 1619.72 | 0.016135 | 1007.97 | 0.000328 | 0.013034 | Q1 hold | 230 | 395 | 625 | 6500 | 40.242 | 0.329 | 28.506 | 0.387 | 11.736 | 0.507 | 34.129 | 0.839 | 12.136 | 0.513 | 21.994 | 0.981 | 0.356 | 0.644 | 0.552 | 51 |
| **27** | PBS_1per_Tween20_replicate | 1006.478 | 0.000320 | 0.009875 | 1619.72 | 0.016135 | 1007.97 | 0.000328 | 0.013034 | Q1 hold | 230 | 405 | 635 | 6500 | 40.47 | 0.295 | 29.326 | 0.573 | 11.144 | 0.525 | 34.71 | 0.752 | 13.22 | 0.759 | 21.49 | 0.818 | 0.381 | 0.619 | 0.615 | 105 |
| **28** | PBS_1per_Tween20_replicate | 1006.478 | 0.000320 | 0.009875 | 1619.72 | 0.016135 | 1007.97 | 0.000328 | 0.013034 | Q1 hold | 230 | 410 | 640 | 6500 | 40.106 | 0.338 | 28.321 | 0.419 | 11.785 | 0.549 | 33.785 | 0.856 | 11.902 | 0.522 | 21.883 | 1.021 | 0.352 | 0.648 | 0.544 | 66 |
| **29** | PBS_1per_Tween20_replicate | 1006.478 | 0.000320 | 0.009875 | 1619.72 | 0.016135 | 1007.97 | 0.000328 | 0.013034 | Q2 hold | 150 | 400 | 550 | 6500 | 40.605 | 0.274 | 25.8 | 0 | 14.805 | 0 | 35.06 | 0.709 | 8.992 | 0 | 26.068 | 0 | 0.256 | 0.744 | 0.345 | 160 |
| **30** | PBS_1per_Tween20_replicate | 1006.478 | 0.000320 | 0.009875 | 1619.72 | 0.016135 | 1007.97 | 0.000328 | 0.013034 | Q2 hold | 190 | 400 | 590 | 6500 | 40.446 | 0.453 | 27.195 | 0.354 | 13.252 | 0.583 | 34.658 | 1.157 | 10.536 | 0.427 | 24.122 | 1.245 | 0.304 | 0.696 | 0.437 | 160 |
| **31** | PBS_1per_Tween20_replicate | 1006.478 | 0.000320 | 0.009875 | 1619.72 | 0.016135 | 1007.97 | 0.000328 | 0.013034 | Q2 hold | 270 | 400 | 670 | 6500 | 40.887 | 1.339 | 30.524 | 0.726 | 10.363 | 1.266 | 35.902 | 3.49 | 14.915 | 1.066 | 20.986 | 3.267 | 0.415 | 0.585 | 0.711 | 68 |
| **32** | PBS_1per_Tween20_replicate | 1006.478 | 0.000320 | 0.009875 | 1619.72 | 0.016135 | 1007.97 | 0.000328 | 0.013034 | Q1 hold | 230 | 280 | 510 | 6500 | 40.265 | 0.948 | 30.751 | 0.961 | 9.514 | 0.583 | 34.237 | 2.42 | 15.27 | 1.441 | 18.967 | 1.529 | 0.446 | 0.554 | 0.805 | 68 |
| **33** | PBS_1per_Tween20_replicate | 1006.478 | 0.000320 | 0.009875 | 1619.72 | 0.016135 | 1007.97 | 0.000328 | 0.013034 | Q1 hold | 230 | 520 | 750 | 6500 | 40 | 0.348 | 26.364 | 0.886 | 13.635 | 1.124 | 33.517 | 0.886 | 9.627 | 0.943 | 23.89 | 1.618 | 0.287 | 0.713 | 0.403 | 64 |

| | | | | | | | | | | | | | | | | | | | | | | | | | | | | | | |
|---|---|---|---|---|---|---|---|---|---|---|---|---|---|---|---|---|---|---|---|---|---|---|---|---|---|---|---|---|---|---|
| **34** | PBS_1per_Tween20_replicate | 1006.478 | 0.000320 | 0.009875 | 1619.72 | 0.016135 | 1007.97 | 0.000328 | 0.013034 | Q1 hold | 230 | 640 | 870 | 6500 | 40.015 | 0.285 | 25.532 | 0.527 | 14.483 | 0.624 | 33.553 | 0.719 | 8.726 | 0.524 | 24.828 | 0.93 | 0.26 | 0.74 | 0.351 | 77 |
| **35** | PBS_1per_Tween20_replicate | 1006.478 | 0.000320 | 0.009875 | 1619.72 | 0.016135 | 1007.97 | 0.000328 | 0.013034 | Q1 hold | 230 | 760 | 990 | 6500 | 39.784 | 0.329 | 24.501 | 0.106 | 15.283 | 0.334 | 32.977 | 0.812 | 7.702 | 0.095 | 25.275 | 0.807 | 0.234 | 0.766 | 0.305 | 147 |
| **36** | PBS_1per_Tween20_replicate | 1006.478 | 0.000320 | 0.009875 | 1619.72 | 0.016135 | 1007.97 | 0.000328 | 0.013034 | Qt | 240 | 390 | 630 | 6500 | 40.482 | 0.649 | 29.167 | 0.632 | 11.315 | 0.885 | 34.764 | 1.632 | 13.01 | 0.836 | 21.753 | 1.796 | 0.374 | 0.626 | 0.598 | 118 |
| **37** | PBS_1per_Tween20_replicate | 1006.478 | 0.000320 | 0.009875 | 1619.72 | 0.016135 | 1007.97 | 0.000328 | 0.013034 | Qt | 235 | 395 | 630 | 6500 | 40.568 | 0.398 | 28.698 | 0.56 | 11.87 | 0.56 | 34.968 | 1.033 | 12.389 | 0.741 | 22.579 | 1.035 | 0.354 | 0.646 | 0.549 | 65 |
| **38** | PBS_1per_Tween20_replicate | 1006.478 | 0.000320 | 0.009875 | 1619.72 | 0.016135 | 1007.97 | 0.000328 | 0.013034 | Qt | 225 | 405 | 630 | 6500 | 40.444 | 0.441 | 28.434 | 0.26 | 12.01 | 0.476 | 34.652 | 1.105 | 12.04 | 0.344 | 22.612 | 1.105 | 0.347 | 0.653 | 0.532 | 119 |
| **39** | PBS_1per_Tween20_replicate | 1006.478 | 0.000320 | 0.009875 | 1619.72 | 0.016135 | 1007.97 | 0.000328 | 0.013034 | Qt | 220 | 410 | 630 | 6500 | 40.501 | 0.315 | 28.366 | 0.4 | 12.135 | 0.489 | 34.792 | 0.808 | 11.958 | 0.505 | 22.834 | 0.916 | 0.344 | 0.656 | 0.524 | 188 |
| **40** | PBS_1per_Tween20_replicate | 1006.478 | 0.000320 | 0.009875 | 1619.72 | 0.016135 | 1007.97 | 0.000328 | 0.013034 | Qt | 150 | 480 | 630 | 6500 | 41.515 | 0.325 | 24.848 | 0.57 | 16.668 | 0.669 | 37.472 | 0.882 | 8.045 | 0.566 | 29.427 | 1.071 | 0.215 | 0.785 | 0.273 | 107 |
| **41** | PBS_1per_Tween20_replicate | 1006.478 | 0.000320 | 0.009875 | 1619.72 | 0.016135 | 1007.97 | 0.000328 | 0.013034 | Qt | 110 | 520 | 630 | 6500 | 42.537 | 0.252 | 23.22 | 0 | 19.317 | 0 | 40.304 | 0.714 | 6.555 | 0 | 33.749 | 0 | 0.163 | 0.837 | 0.194 | 59 |

#### Table S5: All trials no replicates with fluid property parameters

| condition_num | trial | rho_in | sigma_in_oil | mu_in | rho_oil | mu_oil | rho_out | sigma_out_oil | mu_out | sweep_condition | Q1 | Q2 | Qt | Q3 | outer_diam_mean | outer_diam_std | inner_diam_mean | inner_diam_std | shell_diam_mean | shell_diam_std | outer_vol_mean | outer_vol_std | inner_vol_mean | inner_vol_std | shell_vol_mean | shell_vol_std | core_total_ratio | shell_total_ratio | core_shell_ratio | n_droplets |
|---|---|---|---|---|---|---|---|---|---|---|---|---|---|---|---|---|---|---|---|---|---|---|---|---|---|---|---|---|---|---|
| **1** | PBS_1per_Tween20 | 1006.478 | 0.000303 | 0.009875 | 1619.72 | 0.016135 | 1007.97 | 0.000322 | 0.013034 | Q1 hold | 230 | 280 | 510 | 6500 | 39.461 | 1.877 | 29.44 | 0.503 | 10.021 | 1.945 | 32.382 | 4.528 | 13.371 | 0.666 | 19.011 | 4.591 | 0.413 | 0.587 | 0.703 | 28 |
| **2** | PBS_1per_Tween20 | 1006.478 | 0.000303 | 0.009875 | 1619.72 | 0.016135 | 1007.97 | 0.000322 | 0.013034 | Q1 hold | 230 | 390 | 620 | 6500 | 39.617 | 1.413 | 27.314 | 0.679 | 12.303 | 1.76 | 32.681 | 3.513 | 10.69 | 0.8 | 21.991 | 3.849 | 0.327 | 0.673 | 0.486 | 92 |
| **3** | PBS_1per_Tween20 | 1006.478 | 0.000303 | 0.009875 | 1619.72 | 0.016135 | 1007.97 | 0.000322 | 0.013034 | Q1 hold | 230 | 395 | 625 | 6500 | 39.639 | 1.814 | 27.42 | 0.632 | 12.219 | 2.085 | 32.814 | 4.438 | 10.812 | 0.755 | 22.002 | 4.701 | 0.329 | 0.671 | 0.491 | 86 |
| **4** | PBS_1per_Tween20 | 1006.478 | 0.000303 | 0.009875 | 1619.72 | 0.016135 | 1007.97 | 0.000322 | 0.013034 | Q2 hold | 110 | 400 | 510 | 6500 | 40.715 | 0.54 | 24.51 | 0 | 16.205 | 0 | 35.357 | 1.4 | 7.71 | 0 | 27.648 | 0 | 0.218 | 0.782 | 0.279 | 89 |
| **5** | PBS_1per_Tween20 | 1006.478 | 0.000303 | 0.009875 | 1619.72 | 0.016135 | 1007.97 | 0.000322 | 0.013034 | Q2 hold | 150 | 400 | 550 | 6500 | 39.614 | 0.622 | 24.376 | 0.398 | 15.238 | 0.736 | 32.573 | 1.581 | 7.589 | 0.356 | 24.983 | 1.618 | 0.233 | 0.767 | 0.304 | 48 |
| **6** | PBS_1per_Tween20 | 1006.478 | 0.000303 | 0.009875 | 1619.72 | 0.016135 | 1007.97 | 0.000322 | 0.013034 | Q2 hold | 190 | 400 | 590 | 6500 | 38.885 | 0.636 | 25.867 | 0.542 | 13.018 | 0.881 | 30.81 | 1.514 | 9.074 | 0.574 | 21.735 | 1.682 | 0.295 | 0.705 | 0.417 | 96 |
| **7** | PBS_1per_Tween20 | 1006.478 | 0.000303 | 0.009875 | 1619.72 | 0.016135 | 1007.97 | 0.000322 | 0.013034 | Q2 hold | 220 | 400 | 620 | 6500 | 38.865 | 0.619 | 26.877 | 0.481 | 11.988 | 0.779 | 30.762 | 1.485 | 10.176 | 0.529 | 20.586 | 1.57 | 0.331 | 0.669 | 0.494 | 97 |
| **8** | PBS_1per_Tween20 | 1006.478 | 0.000303 | 0.009875 | 1619.72 | 0.016135 | 1007.97 | 0.000322 | 0.013034 | Q2 hold | 225 | 400 | 625 | 6500 | 38.725 | 0.606 | 26.923 | 0.436 | 11.802 | 0.665 | 30.429 | 1.372 | 10.226 | 0.479 | 20.203 | 1.356 | 0.336 | 0.664 | 0.506 | 85 |
| **9** | PBS_1per_Tween20 | 1006.478 | 0.000303 | 0.009875 | 1619.72 | 0.016135 | 1007.97 | 0.000322 | 0.013034 | Qt | 230 | 400 | 630 | 6500 | 39.371 | 0.699 | 26.923 | 0.519 | 12.448 | 0.887 | 31.985 | 1.718 | 10.229 | 0.58 | 21.756 | 1.833 | 0.32 | 0.68 | 0.47 | 85 |

| | | | | | | | | | | | | | | | | | | | | | | | | | | | | | | |
|---|---|---|---|---|---|---|---|---|---|---|---|---|---|---|---|---|---|---|---|---|---|---|---|---|---|---|---|---|---|---|
| **10** | PBS_1per_Tween20 | 1006.478 | 0.000303 | 0.009875 | 1619.72 | 0.016135 | 1007.97 | 0.000322 | 0.013034 | Qt | 225 | 405 | 630 | 6500 | 39.716 | 1.264 | 26.924 | 0.546 | 12.793 | 1.426 | 32.901 | 3.173 | 10.231 | 0.612 | 22.67 | 3.287 | 0.311 | 0.689 | 0.451 | 93 |
| **11** | PBS_1per_Tween20 | 1006.478 | 0.000303 | 0.009875 | 1619.72 | 0.016135 | 1007.97 | 0.000322 | 0.013034 | Q1 hold | 230 | 405 | 635 | 6500 | 39.189 | 1.318 | 27.442 | 0.606 | 11.748 | 1.521 | 31.621 | 3.257 | 10.836 | 0.729 | 20.785 | 3.425 | 0.343 | 0.657 | 0.521 | 99 |
| **12** | PBS_1per_Tween20 | 1006.478 | 0.000303 | 0.009875 | 1619.72 | 0.016135 | 1007.97 | 0.000322 | 0.013034 | Qt | 220 | 410 | 630 | 6500 | 39.041 | 0.732 | 27.09 | 0.194 | 11.951 | 0.756 | 31.191 | 1.813 | 10.411 | 0.225 | 20.78 | 1.826 | 0.334 | 0.666 | 0.501 | 89 |
| **13** | PBS_1per_Tween20 | 1006.478 | 0.000303 | 0.009875 | 1619.72 | 0.016135 | 1007.97 | 0.000322 | 0.013034 | Q1 hold | 230 | 410 | 640 | 6500 | 39.157 | 0.839 | 27.155 | 0.5 | 12.002 | 0.925 | 31.48 | 2.059 | 10.495 | 0.586 | 20.984 | 2.06 | 0.333 | 0.667 | 0.5 | 99 |
| **14** | PBS_1per_Tween20 | 1006.478 | 0.000303 | 0.009875 | 1619.72 | 0.016135 | 1007.97 | 0.000322 | 0.013034 | Qt | 190 | 440 | 630 | 6500 | 38.858 | 0.412 | 25.384 | 0.648 | 13.475 | 0.872 | 30.732 | 0.983 | 8.58 | 0.644 | 22.152 | 1.337 | 0.279 | 0.721 | 0.387 | 127 |
| **15** | PBS_1per_Tween20 | 1006.478 | 0.000303 | 0.009875 | 1619.72 | 0.016135 | 1007.97 | 0.000322 | 0.013034 | Qt | 150 | 480 | 630 | 6500 | 39.629 | 0.382 | 24.51 | 0 | 15.119 | 0 | 32.595 | 0.937 | 7.71 | 0 | 24.885 | 0 | 0.237 | 0.763 | 0.31 | 100 |
| **16** | PBS_1per_Tween20 | 1006.478 | 0.000303 | 0.009875 | 1619.72 | 0.016135 | 1007.97 | 0.000322 | 0.013034 | Qt | 110 | 520 | 630 | 6500 | 40.98 | 0.482 | 21.93 | 0 | 19.05 | 0 | 36.049 | 1.258 | 5.522 | 0 | 30.527 | 0 | 0.153 | 0.847 | 0.181 | 86 |
| **17** | PBS_1per_Tween20 | 1006.478 | 0.000303 | 0.009875 | 1619.72 | 0.016135 | 1007.97 | 0.000322 | 0.013034 | Q1 hold | 230 | 520 | 750 | 6500 | 38.628 | 0.879 | 25.785 | 0.37 | 12.843 | 1.008 | 30.227 | 2.226 | 8.982 | 0.385 | 21.245 | 2.313 | 0.297 | 0.703 | 0.423 | 86 |
| **18** | PBS_1per_Tween20 | 1006.478 | 0.000303 | 0.009875 | 1619.72 | 0.016135 | 1007.97 | 0.000322 | 0.013034 | Q1 hold | 230 | 640 | 870 | 6500 | 38.837 | 0.58 | 24.51 | 0 | 14.327 | 0 | 30.691 | 1.382 | 7.71 | 0 | 22.982 | 0 | 0.251 | 0.749 | 0.335 | 90 |
| **19** | PBS_1per_Tween20 | 1006.478 | 0.000303 | 0.009875 | 1619.72 | 0.016135 | 1007.97 | 0.000322 | 0.013034 | Q1 hold | 230 | 760 | 990 | 6500 | 38.891 | 0.455 | 23.555 | 0.568 | 15.336 | 0.749 | 30.813 | 1.087 | 6.855 | 0.509 | 23.958 | 1.227 | 0.222 | 0.778 | 0.286 | 104 |
| **20** | PBS | 1005.58 | 0.000543 | 0.009311 | 1619.72 | 0.016135 | 1007.97 | 0.000318 | 0.013034 | Q1 hold | 230 | 280 | 510 | 6500 | 45.136 | 0.628 | 34.9 | 0.623 | 10.236 | 0.765 | 48.173 | 2.024 | 22.278 | 1.197 | 25.895 | 2.075 | 0.462 | 0.538 | 0.86 | 111 |
| **21** | PBS | 1005.58 | 0.000543 | 0.009311 | 1619 | 0.016135 | 1007 | 0.000318 | 0.013034 | Qt | 270 | 360 | 630 | 65 | 43.252 | 0.607 | 33.448 | 0.334 | 9.804 | 0.673 | 42.39 | 1.785 | 19.599 | 0.568 | 22.791 | 1.839 | 0.462 | 0.538 | 0.86 | 84 |

| | | | | | | | | | | | | | | | | | | | | | | | | | | | | | | |
|---|---|---|---|---|---|---|---|---|---|---|---|---|---|---|---|---|---|---|---|---|---|---|---|---|---|---|---|---|---|---|
| | | | | | .72 | | .97 | | | | | | | 00 | | | | | | | | | | | | | | | | |
| **22** | PBS | 1005.58 | 0.000543 | 0.009311 | 1619.72 | 0.016135 | 1007.97 | 0.000318 | 0.013034 | Qt | 240 | 390 | 630 | 6500 | 43.359 | 0.739 | 31.002 | 0.23 | 12.357 | 0.8 | 42.717 | 2.174 | 15.604 | 0.362 | 27.113 | 2.244 | 0.365 | 0.635 | 0.576 | 92 |
| **23** | PBS | 1005.58 | 0.000543 | 0.009311 | 1619.72 | 0.016135 | 1007.97 | 0.000318 | 0.013034 | Q2 hold | 110 | 400 | 510 | 6500 | 47.996 | 0.751 | 26.854 | 0.587 | 21.142 | 1.043 | 57.934 | 2.632 | 10.154 | 0.656 | 47.78 | 2.844 | 0.175 | 0.825 | 0.213 | 142 |
| **24** | PBS | 1005.58 | 0.000543 | 0.009311 | 1619.72 | 0.016135 | 1007.97 | 0.000318 | 0.013034 | Q2 hold | 150 | 400 | 550 | 6500 | 45.754 | 0.462 | 28.435 | 0.735 | 17.319 | 0.946 | 50.168 | 1.521 | 12.063 | 0.934 | 38.105 | 1.944 | 0.24 | 0.76 | 0.317 | 93 |
| **25** | PBS | 1005.58 | 0.000543 | 0.009311 | 1619.72 | 0.016135 | 1007.97 | 0.000318 | 0.013034 | Q2 hold | 190 | 400 | 590 | 6500 | 44.22 | 0.765 | 30.451 | 0.635 | 13.77 | 1.06 | 45.316 | 2.382 | 14.803 | 0.916 | 30.513 | 2.665 | 0.327 | 0.673 | 0.485 | 76 |
| **26** | PBS | 1005.58 | 0.000543 | 0.009311 | 1619.72 | 0.016135 | 1007.97 | 0.000318 | 0.013034 | Qt | 230 | 400 | 630 | 6500 | 43.813 | 0.471 | 31.587 | 0.649 | 12.227 | 0.923 | 44.052 | 1.42 | 16.521 | 1.019 | 27.531 | 2.011 | 0.375 | 0.625 | 0.6 | 70 |
| **27** | PBS | 1005.58 | 0.000543 | 0.009311 | 1619.72 | 0.016135 | 1007.97 | 0.000318 | 0.013034 | Q2 hold | 270 | 400 | 670 | 6500 | 46.246 | 1.024 | 35.337 | 0.839 | 10.908 | 1.229 | 51.861 | 3.434 | 23.143 | 1.654 | 28.717 | 3.591 | 0.446 | 0.554 | 0.806 | 89 |
| **28** | PBS | 1005.58 | 0.000543 | 0.009311 | 1619.72 | 0.016135 | 1007.97 | 0.000318 | 0.013034 | Qt | 220 | 410 | 630 | 6500 | 43.846 | 0.528 | 30.525 | 0.613 | 13.321 | 0.858 | 44.155 | 1.605 | 14.911 | 0.885 | 29.244 | 1.929 | 0.338 | 0.662 | 0.51 | 95 |
| **29** | PBS | 1005.58 | 0.000543 | 0.009311 | 1619.72 | 0.016135 | 1007.97 | 0.000318 | 0.013034 | Qt | 190 | 440 | 630 | 6500 | 45.071 | 0.509 | 29.643 | 0.326 | 15.429 | 0.6 | 47.958 | 1.613 | 13.643 | 0.446 | 34.315 | 1.672 | 0.284 | 0.716 | 0.398 | 94 |
| **30** | PBS | 1005.58 | 0.000543 | 0.009311 | 1619.72 | 0.016135 | 1007.97 | 0.000318 | 0.013034 | Qt | 150 | 480 | 630 | 6500 | 46.699 | 0.729 | 27.992 | 0.594 | 18.707 | 0.982 | 53.362 | 2.48 | 11.499 | 0.718 | 41.863 | 2.64 | 0.215 | 0.785 | 0.275 | 113 |
| **31** | PBS | 1005.58 | 0.000543 | 0.009311 | 1619.72 | 0.016135 | 1007.97 | 0.000318 | 0.013034 | Q1 hold | 230 | 520 | 750 | 6500 | 44.248 | 0.42 | 29.554 | 0.371 | 14.694 | 0.625 | 45.372 | 1.286 | 13.522 | 0.491 | 31.85 | 1.486 | 0.298 | 0.702 | 0.425 | 100 |
| **32** | PBS | 1005.58 | 0.000543 | 0.009311 | 1619.72 | 0.016135 | 1007.97 | 0.000318 | 0.013034 | Q1 hold | 230 | 640 | 870 | 6500 | 44.577 | 0.509 | 28.206 | 0.557 | 16.371 | 0.856 | 46.399 | 1.587 | 11.763 | 0.686 | 34.636 | 1.906 | 0.254 | 0.746 | 0.34 | 89 |

| | | | | | | | | | | | | | | | | | | | | | | | | | | | | | | |
|---|---|---|---|---|---|---|---|---|---|---|---|---|---|---|---|---|---|---|---|---|---|---|---|---|---|---|---|---|---|---|
| **33** | PBS | 1005.58 | 0.000543 | 0.0009311 | 1619.72 | 0.0016135 | 1007.97 | 0.000318 | 0.0013034 | Q1 hold | 230 | 760 | 990 | 6500 | 45.795 | 0.935 | 27.69 | 0.647 | 18.105 | 0.953 | 50.349 | 3.031 | 11.135 | 0.782 | 39.214 | 2.88 | 0.221 | 0.779 | 0.284 | 101 |
| **34** | NP40_inner | 1006.07 | 0.00141 | 0.0010028 | 1619.72 | 0.0016135 | 1007.97 | 0.000318 | 0.0013034 | Q1 hold | 240 | 400 | 640 | 6500 | 38.29 | 0.678 | 26.497 | 0.646 | 11.792 | 0.878 | 29.421 | 1.573 | 9.758 | 0.71 | 19.663 | 1.644 | 0.332 | 0.668 | 0.496 | 111 |
| **35** | NP40_inner | 1006.07 | 0.00141 | 0.0010028 | 1619.72 | 0.0016135 | 1007.97 | 0.000318 | 0.0013034 | Q1 hold | 240 | 450 | 690 | 6500 | 38.366 | 0.547 | 25.627 | 0.497 | 12.739 | 0.696 | 29.587 | 1.29 | 8.822 | 0.501 | 20.765 | 1.327 | 0.298 | 0.702 | 0.425 | 127 |
| **36** | NP40_inner | 1006.07 | 0.00141 | 0.0010028 | 1619.72 | 0.0016135 | 1007.97 | 0.000318 | 0.0013034 | Qt | 280 | 460 | 740 | 6500 | 37.776 | 0.519 | 27.09 | 0 | 10.686 | 0 | 28.242 | 1.216 | 10.409 | 0 | 17.833 | 0 | 0.369 | 0.631 | 0.584 | 106 |
| **37** | NP40_inner | 1006.07 | 0.00141 | 0.0010028 | 1619.72 | 0.0016135 | 1007.97 | 0.000318 | 0.0013034 | Qt | 250 | 490 | 740 | 6500 | 38.939 | 0.464 | 25.959 | 0.426 | 12.98 | 0.66 | 30.927 | 1.098 | 9.167 | 0.468 | 21.76 | 1.235 | 0.296 | 0.704 | 0.421 | 178 |
| **38** | NP40_inner | 1006.07 | 0.00141 | 0.0010028 | 1619.72 | 0.0016135 | 1007.97 | 0.000318 | 0.0013034 | Q2 hold | 140 | 500 | 640 | 6500 | 40.587 | 0.6 | 23.22 | 0 | 17.367 | 0 | 35.03 | 1.491 | 6.555 | 0 | 28.474 | 0 | 0.187 | 0.813 | 0.23 | 134 |
| **39** | NP40_inner | 1006.07 | 0.00141 | 0.0010028 | 1619.72 | 0.0016135 | 1007.97 | 0.000318 | 0.0013034 | Q2 hold | 190 | 500 | 690 | 6500 | 39.207 | 0.537 | 24.235 | 0.553 | 14.972 | 0.793 | 31.574 | 1.298 | 7.464 | 0.498 | 24.109 | 1.416 | 0.236 | 0.764 | 0.31 | 136 |
| **40** | NP40_inner | 1006.07 | 0.00141 | 0.0010028 | 1619.72 | 0.0016135 | 1007.97 | 0.000318 | 0.0013034 | Q2 hold | 245 | 500 | 745 | 6500 | 38.823 | 0.741 | 25.832 | 0.201 | 12.991 | 0.773 | 30.672 | 1.766 | 9.027 | 0.22 | 21.645 | 1.786 | 0.294 | 0.706 | 0.417 | 122 |
| **41** | NP40_inner | 1006.07 | 0.00141 | 0.0010028 | 1619.72 | 0.0016135 | 1007.97 | 0.000318 | 0.0013034 | Q2 hold | 270 | 500 | 770 | 6500 | 39.012 | 0.365 | 27.09 | 0 | 11.922 | 0 | 31.095 | 0.869 | 10.409 | 0 | 20.686 | 0 | 0.335 | 0.665 | 0.503 | 89 |
| **42** | NP40_inner | 1006.07 | 0.00141 | 0.0010028 | 1619.72 | 0.0016135 | 1007.97 | 0.000318 | 0.0013034 | Q2 hold | 300 | 500 | 800 | 6500 | 39.584 | 1.017 | 27.146 | 0.269 | 12.438 | 0.864 | 32.539 | 2.603 | 10.477 | 0.325 | 22.062 | 2.395 | 0.322 | 0.678 | 0.475 | 23 |
| **43** | NP40_inner | 1006.07 | 0.00141 | 0.0010028 | 1619.72 | 0.0016135 | 1007.97 | 0.000318 | 0.0013034 | Q2 hold | 330 | 500 | 830 | 6500 | 39.152 | 0.432 | 28.076 | 0.578 | 11.076 | 0.729 | 31.436 | 1.05 | 11.602 | 0.703 | 19.834 | 1.278 | 0.369 | 0.631 | 0.585 | 106 |
| **44** | NP40_inner | 1006.07 | 0.00141 | 0.0010028 | 1619 | 0.0016135 | 1007 | 0.000318 | 0.0013034 | Q2 hold | 360 | 500 | 860 | 65 | 39.255 | 0.528 | 24.6 | 0.33 | 14.655 | 0.638 | 31.689 | 1.28 | 7.799 | 0.328 | 23.89 | 1.338 | 0.246 | 0.754 | 0.326 | 143 |

| | | | | | | | | | | | | | | | | | | | | | | | | | | | | | | |
|---|---|---|---|---|---|---|---|---|---|---|---|---|---|---|---|---|---|---|---|---|---|---|---|---|---|---|---|---|---|---|
| | | | | | .72 | | .97 | | | | | | | 00 | | | | | | | | | | | | | | | | |
| **45** | NP40_inner | 1006.07 | 0.00141 | 0.010028 | 1619.72 | 0.016135 | 1007.97 | 0.000318 | 0.013034 | Qt | 235 | 505 | 740 | 6500 | 38.007 | 0.808 | 24.552 | 0.229 | 13.455 | 0.827 | 28.784 | 1.695 | 7.751 | 0.228 | 21.033 | 1.696 | 0.269 | 0.731 | 0.369 | 154 |
| **46** | NP40_inner | 1006.07 | 0.00141 | 0.010028 | 1619.72 | 0.016135 | 1007.97 | 0.000318 | 0.013034 | Qt | 230 | 510 | 740 | 6500 | 38.032 | 0.522 | 24.519 | 0.108 | 13.513 | 0.564 | 28.82 | 1.18 | 7.718 | 0.107 | 21.101 | 1.215 | 0.268 | 0.732 | 0.366 | 144 |
| **47** | NP40_inner | 1006.07 | 0.00141 | 0.010028 | 1619.72 | 0.016135 | 1007.97 | 0.000318 | 0.013034 | Qt | 190 | 550 | 740 | 6500 | 39.388 | 0.463 | 24.51 | 0 | 14.878 | 0 | 32.008 | 1.125 | 7.71 | 0 | 24.299 | 0 | 0.241 | 0.759 | 0.317 | 132 |
| **48** | NP40_inner | 1006.07 | 0.00141 | 0.010028 | 1619.72 | 0.016135 | 1007.97 | 0.000318 | 0.013034 | Q1 hold | 240 | 550 | 790 | 6500 | 38.454 | 0.441 | 24.559 | 0.248 | 13.895 | 0.563 | 29.785 | 1.036 | 7.758 | 0.247 | 22.027 | 1.129 | 0.26 | 0.74 | 0.352 | 79 |
| **49** | NP40_inner | 1006.07 | 0.00141 | 0.010028 | 1619.72 | 0.016135 | 1007.97 | 0.000318 | 0.013034 | Qt | 140 | 600 | 740 | 6500 | 41.145 | 1.164 | 22.124 | 0.463 | 19.021 | 1.272 | 36.551 | 2.58 | 5.677 | 0.371 | 30.874 | 2.629 | 0.155 | 0.845 | 0.184 | 100 |
| **50** | NP40_inner | 1006.07 | 0.00141 | 0.010028 | 1619.72 | 0.016135 | 1007.97 | 0.000318 | 0.013034 | Q1 hold | 240 | 600 | 840 | 6500 | 39.065 | 0.805 | 24.356 | 0.455 | 14.709 | 0.995 | 31.254 | 1.969 | 7.573 | 0.411 | 23.681 | 2.085 | 0.242 | 0.758 | 0.32 | 109 |
| **51** | NP40_inner | 1006.07 | 0.00141 | 0.010028 | 1619.72 | 0.016135 | 1007.97 | 0.000318 | 0.013034 | Q1 hold | 240 | 650 | 890 | 6500 | 38.835 | 0.651 | 23.821 | 0.646 | 15.014 | 1.011 | 30.692 | 1.55 | 7.093 | 0.578 | 23.599 | 1.765 | 0.231 | 0.769 | 0.301 | 118 |
| **52** | NP40_inner | 1006.07 | 0.00141 | 0.010028 | 1619.72 | 0.016135 | 1007.97 | 0.000318 | 0.013034 | Q1 hold | 240 | 700 | 940 | 6500 | 38.413 | 0.414 | 23.22 | 0 | 15.193 | 0 | 29.689 | 0.956 | 6.555 | 0 | 23.134 | 0 | 0.221 | 0.779 | 0.283 | 99 |
| **53** | NP40_inner | 1006.07 | 0.00141 | 0.010028 | 1619.72 | 0.016135 | 1007.97 | 0.000318 | 0.013034 | Q1 hold | 240 | 750 | 990 | 6500 | 38.64 | 0.379 | 23.22 | 0 | 15.42 | 0 | 30.215 | 0.89 | 6.555 | 0 | 23.659 | 0 | 0.217 | 0.783 | 0.277 | 32 |
| **54** | NP40_inner | 1006.07 | 0.00141 | 0.010028 | 1619.72 | 0.016135 | 1007.97 | 0.000318 | 0.013034 | Q1 hold | 240 | 800 | 1040 | 6500 | 38.493 | 0.447 | 23.22 | 0 | 15.273 | 0 | 29.876 | 1.037 | 6.555 | 0 | 23.32 | 0 | 0.219 | 0.781 | 0.281 | 162 |
| **55** | NP40_inner | 1006.07 | 0.00141 | 0.010028 | 1619.72 | 0.016135 | 1007.97 | 0.000318 | 0.013034 | Q1 hold | 240 | 850 | 1090 | 6500 | 38.751 | 0.549 | 21.93 | 0 | 16.821 | 0 | 30.486 | 1.287 | 5.522 | 0 | 24.964 | 0 | 0.181 | 0.819 | 0.221 | 38 |

| | | | | | | | | | | | | | | | | | | | | | | | | | | | | | | |
|---|---|---|---|---|---|---|---|---|---|---|---|---|---|---|---|---|---|---|---|---|---|---|---|---|---|---|---|---|---|---|
| **56** | NP40_inner | 1006.07 | 0.00141 | 0.0010028 | 1619.72 | 0.0016135 | 1007.97 | 0.000318 | 0.0013034 | Q1 hold | 240 | 495 | 735 | 6500 | 39.907 | 0.396 | 26.197 | 0.603 | 13.71 | 0.793 | 33.288 | 0.993 | 9.428 | 0.663 | 23.859 | 1.311 | 0.283 | 0.717 | 0.395 | 39 |
| **57** | NP40_inner | 1006.07 | 0.00141 | 0.0010028 | 1619.72 | 0.0016135 | 1007.97 | 0.000318 | 0.0013034 | Qt | 245 | 495 | 740 | 6500 | 39.189 | 0.623 | 25.855 | 0.262 | 13.334 | 0.702 | 31.537 | 1.543 | 9.052 | 0.288 | 22.485 | 1.6 | 0.287 | 0.713 | 0.403 | 94 |
| **58** | NP40_inner | 1006.07 | 0.00141 | 0.0010028 | 1619.72 | 0.0016135 | 1007.97 | 0.000318 | 0.0013034 | Q2 hold | 230 | 500 | 730 | 6500 | 38.874 | 0.498 | 25.8 | 0 | 13.074 | 0 | 30.775 | 1.192 | 8.992 | 0 | 21.783 | 0 | 0.292 | 0.708 | 0.413 | 78 |
| **59** | NP40_inner | 1006.07 | 0.00141 | 0.0010028 | 1619.72 | 0.0016135 | 1007.97 | 0.000318 | 0.0013034 | Q2 hold | 235 | 500 | 735 | 6500 | 38.928 | 0.745 | 25.8 | 0 | 13.128 | 0 | 30.92 | 1.784 | 8.992 | 0 | 21.928 | 0 | 0.291 | 0.709 | 0.41 | 105 |
| **60** | NP40_inner | 1006.07 | 0.00141 | 0.0010028 | 1619.72 | 0.0016135 | 1007.97 | 0.000318 | 0.0013034 | Q2 hold | 240 | 500 | 740 | 6500 | 38.877 | 0.585 | 25.077 | 0.665 | 13.8 | 0.994 | 30.786 | 1.397 | 8.275 | 0.665 | 22.512 | 1.694 | 0.269 | 0.731 | 0.368 | 116 |
| **61** | NP40_inner | 1006.07 | 0.00141 | 0.0010028 | 1619.72 | 0.0016135 | 1007.97 | 0.000318 | 0.0013034 | Q2 hold | 250 | 500 | 750 | 6500 | 38.519 | 0.491 | 25.37 | 0.752 | 13.149 | 1.027 | 29.938 | 1.15 | 8.572 | 0.765 | 21.366 | 1.575 | 0.286 | 0.714 | 0.401 | 54 |
| **62** | NP40_inner | 1006.07 | 0.00141 | 0.0010028 | 1619.72 | 0.0016135 | 1007.97 | 0.000318 | 0.0013034 | Q1 hold | 240 | 505 | 745 | 6500 | 38.778 | 0.439 | 25.304 | 0.66 | 13.474 | 0.919 | 30.544 | 1.045 | 8.5 | 0.66 | 22.043 | 1.428 | 0.278 | 0.722 | 0.386 | 91 |
| **63** | NP40_inner | 1006.07 | 0.00141 | 0.0010028 | 1619.72 | 0.0016135 | 1007.97 | 0.000318 | 0.0013034 | Q1 hold | 240 | 510 | 750 | 6500 | 38.879 | 0.598 | 25.395 | 0.747 | 13.484 | 1.13 | 30.793 | 1.428 | 8.597 | 0.761 | 22.195 | 1.866 | 0.279 | 0.721 | 0.387 | 86 |
| **64** | NP40_inner | 1006.07 | 0.00141 | 0.0010028 | 1619.72 | 0.0016135 | 1007.97 | 0.000318 | 0.0013034 | Q1 hold | 90 | 650 | 740 | 6500 | 44.592 | 0.639 | 20.913 | 0.532 | 23.679 | 0.821 | 46.454 | 1.969 | 4.798 | 0.379 | 41.656 | 1.996 | 0.103 | 0.897 | 0.115 | 52 |
| **65** | M9 | 1013 | 0.01284 | 0.0008615 | 1619.72 | 0.0016135 | 1013.4 | 0.000522 | 0.0014117 | Q1 hold | 250 | 350 | 600 | 6500 | 41.897 | 1.831 | 30.796 | 0.431 | 11.101 | 1.949 | 38.725 | 4.921 | 15.301 | 0.623 | 23.424 | 5.06 | 0.395 | 0.605 | 0.653 | 157 |
| **66** | M9 | 1013 | 0.01284 | 0.0008615 | 1619.72 | 0.0016135 | 1013.4 | 0.000522 | 0.0014117 | Q1 hold | 250 | 400 | 650 | 6500 | 41.919 | 0.893 | 29.659 | 0.118 | 12.26 | 0.901 | 38.621 | 2.476 | 13.661 | 0.157 | 24.96 | 2.481 | 0.354 | 0.646 | 0.547 | 119 |
| **67** | M9 | 1013 | 0.01284 | 0.0008615 | 1619 | 0.0016135 | 1013.4 | 0.000522 | 0.0014117 | Q1 hold | 250 | 440 | 690 | 65 | 41.334 | 0.571 | 28.38 | 0 | 12.954 | 0 | 36.997 | 1.611 | 11.968 | 0 | 25.029 | 0 | 0.323 | 0.677 | 0.478 | 103 |

| | | | | | | | | | | | | | | | | | | | | | | | | | | | | | | |
|---|---|---|---|---|---|---|---|---|---|---|---|---|---|---|---|---|---|---|---|---|---|---|---|---|---|---|---|---|---|---|
| | | | | | .72 | | | | | | | | | 00 | | | | | | | | | | | | | | | | |
| **68** | M9 | 1013 | 0.01284 | 0.008615 | 1619.72 | 0.016135 | 1013.4 | 0.000522 | 0.014117 | Qt | 290 | 410 | 700 | 6500 | 41.929 | 0.428 | 30.96 | 0 | 10.969 | 0 | 38.609 | 1.181 | 15.538 | 0 | 23.07 | 0 | 0.402 | 0.598 | 0.674 | 105 |
| **69** | M9 | 1013 | 0.01284 | 0.008615 | 1619.72 | 0.016135 | 1013.4 | 0.000522 | 0.014117 | Qt | 270 | 430 | 700 | 6500 | 42.33 | 0.86 | 30.28 | 0.646 | 12.05 | 1.025 | 39.763 | 2.45 | 14.557 | 0.933 | 25.207 | 2.536 | 0.366 | 0.634 | 0.577 | 148 |
| **70** | M9 | 1013 | 0.01284 | 0.008615 | 1619.72 | 0.016135 | 1013.4 | 0.000522 | 0.014117 | Qt | 260 | 440 | 700 | 6500 | 41.838 | 0.657 | 29.356 | 0.556 | 12.482 | 0.893 | 38.374 | 1.821 | 13.26 | 0.736 | 25.114 | 2.021 | 0.346 | 0.654 | 0.528 | 115 |
| **71** | M9 | 1013 | 0.01284 | 0.008615 | 1619.72 | 0.016135 | 1013.4 | 0.000522 | 0.014117 | Q1 hold | 250 | 445 | 695 | 6500 | 41.025 | 0.491 | 28.38 | 0 | 12.645 | 0 | 36.168 | 1.296 | 11.968 | 0 | 24.2 | 0 | 0.331 | 0.669 | 0.495 | 140 |
| **72** | M9 | 1013 | 0.01284 | 0.008615 | 1619.72 | 0.016135 | 1013.4 | 0.000522 | 0.014117 | Qt | 255 | 445 | 700 | 6500 | 42.066 | 0.854 | 29.386 | 0.537 | 12.68 | 1.095 | 39.023 | 2.34 | 13.3 | 0.711 | 25.723 | 2.579 | 0.341 | 0.659 | 0.517 | 109 |
| **73** | M9 | 1013 | 0.01284 | 0.008615 | 1619.72 | 0.016135 | 1013.4 | 0.000522 | 0.014117 | Q2 hold | 210 | 450 | 660 | 6500 | 42.196 | 0.631 | 27.919 | 0.621 | 14.277 | 0.919 | 39.363 | 1.807 | 11.411 | 0.75 | 27.953 | 2.009 | 0.29 | 0.71 | 0.408 | 123 |
| **74** | M9 | 1013 | 0.01284 | 0.008615 | 1619.72 | 0.016135 | 1013.4 | 0.000522 | 0.014117 | Q2 hold | 240 | 450 | 690 | 6500 | 40.979 | 0.945 | 28.38 | 0 | 12.599 | 0 | 36.088 | 2.25 | 11.968 | 0 | 24.119 | 0 | 0.332 | 0.668 | 0.496 | 118 |
| **75** | M9 | 1013 | 0.01284 | 0.008615 | 1619.72 | 0.016135 | 1013.4 | 0.000522 | 0.014117 | Q2 hold | 245 | 450 | 695 | 6500 | 41.547 | 0.68 | 28.448 | 0.393 | 13.099 | 0.827 | 37.58 | 1.874 | 12.061 | 0.51 | 25.518 | 2.001 | 0.321 | 0.679 | 0.473 | 95 |
| **76** | M9 | 1013 | 0.01284 | 0.008615 | 1619.72 | 0.016135 | 1013.4 | 0.000522 | 0.014117 | Q2 hold | 250 | 450 | 700 | 6500 | 41.972 | 0.714 | 29.185 | 0.628 | 12.787 | 0.865 | 38.748 | 1.971 | 13.034 | 0.831 | 25.714 | 2.003 | 0.336 | 0.664 | 0.507 | 117 |
| **77** | M9 | 1013 | 0.01284 | 0.008615 | 1619.72 | 0.016135 | 1013.4 | 0.000522 | 0.014117 | Q2 hold | 255 | 450 | 705 | 6500 | 41.357 | 1.274 | 28.393 | 0.846 | 12.964 | 1.63 | 37.133 | 2.708 | 12.016 | 1.072 | 25.117 | 3.094 | 0.324 | 0.676 | 0.478 | 101 |
| **78** | M9 | 1013 | 0.01284 | 0.008615 | 1619.72 | 0.016135 | 1013.4 | 0.000522 | 0.014117 | Q2 hold | 260 | 450 | 710 | 6500 | 41.127 | 0.536 | 28.368 | 0.126 | 12.76 | 0.569 | 36.442 | 1.425 | 11.953 | 0.153 | 24.489 | 1.456 | 0.328 | 0.672 | 0.488 | 104 |

| | | | | | | | | | | | | | | | | | | | | | | | | | | | | | | |
|---|---|---|---|---|---|---|---|---|---|---|---|---|---|---|---|---|---|---|---|---|---|---|---|---|---|---|---|---|---|---|
| **79** | M9 | 1013 | 0.01284 | 0.0008615 | 1619.72 | 0.0016135 | 1013.4 | 0.000522 | 0.0014117 | Q2 hold | 350 | 450 | 800 | 6500 | 41.276 | 0.611 | 30.96 | 0 | 10.316 | 0 | 36.846 | 1.644 | 15.538 | 0 | 21.307 | 0 | 0.422 | 0.578 | 0.729 | 134 |
| **80** | M9 | 1013 | 0.01284 | 0.0008615 | 1619.72 | 0.0016135 | 1013.4 | 0.000522 | 0.0014117 | Q2 hold | 400 | 450 | 850 | 6500 | 42.227 | 0.799 | 32.298 | 0.246 | 9.929 | 0.77 | 39.467 | 2.308 | 17.645 | 0.418 | 21.822 | 2.226 | 0.447 | 0.553 | 0.809 | 107 |
| **81** | M9 | 1013 | 0.01284 | 0.0008615 | 1619.72 | 0.0016135 | 1013.4 | 0.000522 | 0.0014117 | Q2 hold | 450 | 450 | 900 | 6500 | 42.353 | 0.795 | 33.125 | 0.605 | 9.228 | 0.903 | 39.821 | 2.3 | 19.051 | 1.029 | 20.771 | 2.332 | 0.478 | 0.522 | 0.917 | 112 |
| **82** | M9 | 1013 | 0.01284 | 0.0008615 | 1619.72 | 0.0016135 | 1013.4 | 0.000522 | 0.0014117 | Q2 hold | 550 | 450 | 1000 | 6500 | 42.532 | 0.857 | 33.54 | 0 | 8.992 | 0 | 40.335 | 2.472 | 19.755 | 0 | 20.579 | 0 | 0.49 | 0.51 | 0.96 | 108 |
| **83** | M9 | 1013 | 0.01284 | 0.0008615 | 1619.72 | 0.0016135 | 1013.4 | 0.000522 | 0.0014117 | Q2 hold | 650 | 450 | 1100 | 6500 | 43.553 | 1.002 | 36.046 | 0.301 | 7.507 | 1.013 | 43.324 | 2.993 | 24.528 | 0.595 | 18.796 | 2.992 | 0.566 | 0.434 | 1.305 | 105 |
| **84** | M9 | 1013 | 0.01284 | 0.0008615 | 1619.72 | 0.0016135 | 1013.4 | 0.000522 | 0.0014117 | Qt | 245 | 455 | 700 | 6500 | 42.036 | 0.821 | 29.102 | 0.686 | 12.935 | 1.11 | 38.937 | 2.337 | 12.926 | 0.908 | 26.012 | 2.571 | 0.332 | 0.668 | 0.497 | 118 |
| **85** | M9 | 1013 | 0.01284 | 0.0008615 | 1619.72 | 0.0016135 | 1013.4 | 0.000522 | 0.0014117 | Q1 hold | 250 | 455 | 705 | 6500 | 41.309 | 0.64 | 28.38 | 0.501 | 12.929 | 0.76 | 36.935 | 1.722 | 11.979 | 0.635 | 24.956 | 1.747 | 0.324 | 0.676 | 0.48 | 107 |
| **86** | M9 | 1013 | 0.01284 | 0.0008615 | 1619.72 | 0.0016135 | 1013.4 | 0.000522 | 0.0014117 | Qt | 240 | 460 | 700 | 6500 | 42.048 | 0.62 | 28.589 | 0.718 | 13.459 | 0.984 | 38.95 | 1.725 | 12.258 | 0.924 | 26.693 | 2.01 | 0.315 | 0.685 | 0.459 | 105 |
| **87** | M9 | 1013 | 0.01284 | 0.0008615 | 1619.72 | 0.0016135 | 1013.4 | 0.000522 | 0.0014117 | Q1 hold | 250 | 460 | 710 | 6500 | 41.473 | 0.534 | 28.351 | 0.192 | 13.122 | 0.582 | 37.368 | 1.454 | 11.933 | 0.232 | 25.435 | 1.49 | 0.319 | 0.681 | 0.469 | 133 |
| **88** | M9 | 1013 | 0.01284 | 0.0008615 | 1619.72 | 0.0016135 | 1013.4 | 0.000522 | 0.0014117 | Qt | 220 | 480 | 700 | 6500 | 42.458 | 0.675 | 28.268 | 0.364 | 14.19 | 0.719 | 40.106 | 1.889 | 11.833 | 0.44 | 28.273 | 1.875 | 0.295 | 0.705 | 0.419 | 104 |
| **89** | M9 | 1013 | 0.01284 | 0.0008615 | 1619.72 | 0.0016135 | 1013.4 | 0.000522 | 0.0014117 | Qt | 200 | 500 | 700 | 6500 | 43.175 | 1.254 | 27.469 | 0.59 | 15.707 | 1.338 | 42.239 | 2.986 | 10.867 | 0.713 | 31.372 | 3.005 | 0.257 | 0.743 | 0.346 | 109 |
| **90** | M9 | 1013 | 0.01284 | 0.0008615 | 1619 | 0.0016135 | 1013.4 | 0.000522 | 0.0014117 | Q1 hold | 250 | 530 | 780 | 65 | 41.932 | 0.787 | 28.105 | 0.531 | 13.826 | 0.988 | 38.643 | 2.095 | 11.636 | 0.641 | 27.007 | 2.257 | 0.301 | 0.699 | 0.431 | 108 |

| | | | | | | | | | | | | | | | | | | | | | | | | | | | | | | |
|---|---|---|---|---|---|---|---|---|---|---|---|---|---|---|---|---|---|---|---|---|---|---|---|---|---|---|---|---|---|---|
| | | | | | .72 | | | | | | | | | 00 | | | | | | | | | | | | | | | | |
| **91** | M9 | 1013 | 0.01284 | 0.008615 | 1619.72 | 0.016135 | 1013.4 | 0.000522 | 0.014117 | Q1 hold | 250 | 650 | 900 | 6500 | 42.589 | 0.905 | 27.059 | 0.257 | 15.53 | 0.874 | 40.5 | 2.343 | 10.377 | 0.289 | 30.123 | 2.297 | 0.256 | 0.744 | 0.344 | 125 |
| **92** | M9 | 1013 | 0.01284 | 0.008615 | 1619.72 | 0.016135 | 1013.4 | 0.000522 | 0.014117 | Q1 hold | 250 | 750 | 1000 | 6500 | 42.991 | 1.352 | 26.481 | 0.647 | 16.51 | 1.575 | 41.715 | 3.099 | 9.74 | 0.711 | 31.975 | 3.279 | 0.233 | 0.767 | 0.305 | 108 |
| **93** | M9 | 1013 | 0.01284 | 0.008615 | 1619.72 | 0.016135 | 1013.4 | 0.000522 | 0.014117 | Q1 hold | 250 | 850 | 1100 | 6500 | 43.739 | 0.501 | 25.662 | 0.437 | 18.077 | 0.772 | 43.831 | 1.505 | 8.856 | 0.438 | 34.975 | 1.709 | 0.202 | 0.798 | 0.253 | 112 |
| **94** | M9 | 1013 | 0.01284 | 0.008615 | 1619.72 | 0.016135 | 1013.4 | 0.000522 | 0.014117 | Qt | 290 | 410 | 700 | 6500 | 41.929 | 0.428 | 30.96 | 0 | 10.969 | 0 | 38.609 | 1.181 | 15.538 | 0 | 23.07 | 0 | 0.402 | 0.598 | 0.674 | 105 |
| **95** | M9 | 1013 | 0.01284 | 0.008615 | 1619.72 | 0.016135 | 1013.4 | 0.000522 | 0.014117 | Qt | 270 | 430 | 700 | 6500 | 42.33 | 0.86 | 30.28 | 0.646 | 12.05 | 1.025 | 39.763 | 2.45 | 14.557 | 0.933 | 25.207 | 2.536 | 0.366 | 0.634 | 0.577 | 148 |
| **96** | M9 | 1013 | 0.01284 | 0.008615 | 1619.72 | 0.016135 | 1013.4 | 0.000522 | 0.014117 | Qt | 200 | 500 | 700 | 6500 | 43.175 | 1.254 | 27.469 | 0.59 | 15.707 | 1.338 | 42.239 | 2.986 | 10.867 | 0.713 | 31.372 | 3.005 | 0.257 | 0.743 | 0.346 | 109 |
| **97** | M9_glucose | 1017.5 | 0.0116 | 0.009668 | 1619.72 | 0.016135 | 1017.9 | 0.000458 | 0.015630 | Q1 hold | 200 | 250 | 450 | 6500 | 46.456 | 0.883 | 35.55 | 0.644 | 10.907 | 1 | 52.553 | 3.024 | 23.547 | 1.273 | 29.006 | 3.074 | 0.448 | 0.552 | 0.812 | 95 |
| **98** | M9_glucose | 1017.5 | 0.0116 | 0.009668 | 1619.72 | 0.016135 | 1017.9 | 0.000458 | 0.015630 | Q1 hold | 200 | 300 | 500 | 6500 | 45.053 | 0.842 | 32.798 | 0.641 | 12.255 | 1.103 | 47.93 | 2.699 | 18.494 | 1.089 | 29.437 | 3.005 | 0.386 | 0.614 | 0.628 | 106 |
| **99** | M9_glucose | 1017.5 | 0.0116 | 0.009668 | 1619.72 | 0.016135 | 1017.9 | 0.000458 | 0.015630 | Qt | 350 | 300 | 650 | 6500 | 44.091 | 1.349 | 35.636 | 0.655 | 8.455 | 1.243 | 45.004 | 4.103 | 23.72 | 1.289 | 21.284 | 3.789 | 0.527 | 0.473 | 1.114 | 96 |
| **100** | M9_glucose | 1017.5 | 0.0116 | 0.009668 | 1619.72 | 0.016135 | 1017.9 | 0.000458 | 0.015630 | Q1 hold | 200 | 350 | 550 | 6500 | 44.914 | 0.998 | 32.124 | 0.387 | 12.79 | 1.098 | 47.507 | 3 | 17.364 | 0.608 | 30.143 | 3.108 | 0.366 | 0.634 | 0.576 | 51 |
| **101** | M9_glucose | 1017.5 | 0.0116 | 0.009668 | 1619.72 | 0.016135 | 1017.9 | 0.000458 | 0.015630 | Qt | 300 | 350 | 650 | 6500 | 42.573 | 0.536 | 33.192 | 0.576 | 9.382 | 0.588 | 40.422 | 1.533 | 19.163 | 0.978 | 21.259 | 1.41 | 0.474 | 0.526 | 0.901 | 100 |

| | | | | | | | | | | | | | | | | | | | | | | | | | | | | | | |
|---|---|---|---|---|---|---|---|---|---|---|---|---|---|---|---|---|---|---|---|---|---|---|---|---|---|---|---|---|---|---|
| **102** | M9_glucose | 1017.5 | 0.0116 | 0.0009668 | 1619.72 | 0.0016135 | 1017.9 | 0.000458 | 0.0015630 | Q1 hold | 200 | 400 | 600 | 6500 | 45.504 | 0.812 | 30.935 | 0.179 | 14.569 | 0.884 | 49.381 | 2.672 | 15.502 | 0.258 | 33.879 | 2.763 | 0.314 | 0.686 | 0.458 | 103 |
| **103** | M9_glucose | 1017.5 | 0.0116 | 0.0009668 | 1619.72 | 0.0016135 | 1017.9 | 0.000458 | 0.0015630 | Qt | 250 | 400 | 650 | 6500 | 42.69 | 1.016 | 30.866 | 0.337 | 11.824 | 1.095 | 40.807 | 3.009 | 15.403 | 0.486 | 25.404 | 3.082 | 0.377 | 0.623 | 0.606 | 110 |
| **104** | M9_glucose | 1017.5 | 0.0116 | 0.0009668 | 1619.72 | 0.0016135 | 1017.9 | 0.000458 | 0.0015630 | Q1 hold | 200 | 440 | 640 | 6500 | 43.543 | 0.817 | 29.528 | 0.442 | 14.015 | 0.865 | 43.272 | 2.478 | 13.489 | 0.59 | 29.783 | 2.452 | 0.312 | 0.688 | 0.453 | 109 |
| **105** | M9_glucose | 1017.5 | 0.0116 | 0.0009668 | 1619.72 | 0.0016135 | 1017.9 | 0.000458 | 0.0015630 | Qt | 210 | 440 | 650 | 6500 | 42.781 | 0.713 | 29.146 | 0.687 | 13.634 | 0.917 | 41.03 | 2.064 | 12.986 | 0.9 | 28.044 | 2.125 | 0.316 | 0.684 | 0.463 | 101 |
| **106** | M9_glucose | 1017.5 | 0.0116 | 0.0009668 | 1619.72 | 0.0016135 | 1017.9 | 0.000458 | 0.0015630 | Q1 hold | 200 | 445 | 645 | 6500 | 43.284 | 0.546 | 29.186 | 0.653 | 14.098 | 0.895 | 42.481 | 1.648 | 13.037 | 0.859 | 29.444 | 1.933 | 0.307 | 0.693 | 0.443 | 104 |
| **107** | M9_glucose | 1017.5 | 0.0116 | 0.0009668 | 1619.72 | 0.0016135 | 1017.9 | 0.000458 | 0.0015630 | Qt | 205 | 445 | 650 | 6500 | 43.171 | 0.566 | 29.774 | 0.353 | 13.397 | 0.617 | 42.149 | 1.612 | 13.826 | 0.509 | 28.323 | 1.609 | 0.328 | 0.672 | 0.488 | 124 |
| **108** | M9_glucose | 1017.5 | 0.0116 | 0.0009668 | 1619.72 | 0.0016135 | 1017.9 | 0.000458 | 0.0015630 | Q2 hold | 150 | 450 | 600 | 6500 | 44.747 | 0.543 | 28.051 | 0.566 | 16.697 | 0.853 | 46.934 | 1.709 | 11.57 | 0.683 | 35.364 | 1.952 | 0.247 | 0.753 | 0.327 | 94 |
| **109** | M9_glucose | 1017.5 | 0.0116 | 0.0009668 | 1619.72 | 0.0016135 | 1017.9 | 0.000458 | 0.0015630 | Q2 hold | 190 | 450 | 640 | 6500 | 43.356 | 0.757 | 28.989 | 0.667 | 14.367 | 0.994 | 42.711 | 2.313 | 12.776 | 0.879 | 29.935 | 2.445 | 0.299 | 0.701 | 0.427 | 127 |
| **110** | M9_glucose | 1017.5 | 0.0116 | 0.0009668 | 1619.72 | 0.0016135 | 1017.9 | 0.000458 | 0.0015630 | Q2 hold | 195 | 450 | 645 | 6500 | 43.213 | 0.595 | 29.427 | 0.507 | 13.786 | 0.807 | 42.275 | 1.773 | 13.354 | 0.671 | 28.921 | 1.937 | 0.316 | 0.684 | 0.462 | 122 |
| **111** | M9_glucose | 1017.5 | 0.0116 | 0.0009668 | 1619.72 | 0.0016135 | 1017.9 | 0.000458 | 0.0015630 | Qt | 200 | 450 | 650 | 6500 | 43.211 | 0.977 | 29.464 | 0.475 | 13.747 | 1.119 | 42.312 | 2.922 | 13.403 | 0.628 | 28.908 | 3.033 | 0.317 | 0.683 | 0.464 | 119 |
| **112** | M9_glucose | 1017.5 | 0.0116 | 0.0009668 | 1619.72 | 0.0016135 | 1017.9 | 0.000458 | 0.0015630 | Q2 hold | 205 | 450 | 655 | 6500 | 42.996 | 0.637 | 29.419 | 0.513 | 13.577 | 0.874 | 41.645 | 1.874 | 13.343 | 0.679 | 28.302 | 2.087 | 0.32 | 0.68 | 0.471 | 113 |
| **113** | M9_glucose | 1017.5 | 0.0116 | 0.0009668 | 1619 | 0.0016135 | 1017.9 | 0.000458 | 0.0015630 | Q2 hold | 210 | 450 | 660 | 65 | 43.134 | 0.63 | 29.658 | 0.122 | 13.475 | 0.643 | 42.046 | 1.873 | 13.661 | 0.161 | 28.386 | 1.881 | 0.325 | 0.675 | 0.481 | 112 |

| | | | | | | | | | | | | | | | | | | | | | | | | | | | | | | |
|---|---|---|---|---|---|---|---|---|---|---|---|---|---|---|---|---|---|---|---|---|---|---|---|---|---|---|---|---|---|---|
| | | | | | .72 | | | | | | | | | 00 | | | | | | | | | | | | | | | | |
| **114** | M9_glucose | 1017.5 | 0.0116 | 0.0009668 | 1619.72 | 0.0016135 | 1017.9 | 0.000458 | 0.0015630 | Q2 hold | 250 | 450 | 700 | 6500 | 42.413 | 0.867 | 30.54 | 0.945 | 11.873 | 1.168 | 39.996 | 2.387 | 14.957 | 1.407 | 25.039 | 2.528 | 0.374 | 0.626 | 0.597 | 172 |
| **115** | M9_glucose | 1017.5 | 0.0116 | 0.0009668 | 1619.72 | 0.0016135 | 1017.9 | 0.000458 | 0.0015630 | Q2 hold | 300 | 450 | 750 | 6500 | 42.165 | 0.741 | 31.149 | 0.554 | 11.016 | 0.828 | 39.287 | 2.099 | 15.839 | 0.863 | 23.448 | 2.101 | 0.403 | 0.597 | 0.676 | 205 |
| **116** | M9_glucose | 1017.5 | 0.0116 | 0.0009668 | 1619.72 | 0.0016135 | 1017.9 | 0.000458 | 0.0015630 | Q2 hold | 350 | 450 | 800 | 6500 | 42.445 | 1.237 | 32.598 | 0.664 | 9.847 | 1.269 | 40.138 | 3.432 | 18.16 | 1.115 | 21.979 | 3.397 | 0.452 | 0.548 | 0.826 | 63 |
| **117** | M9_glucose | 1017.5 | 0.0116 | 0.0009668 | 1619.72 | 0.0016135 | 1017.9 | 0.000458 | 0.0015630 | Qt | 195 | 455 | 650 | 6500 | 43.56 | 0.547 | 29.61 | 0.273 | 13.95 | 0.609 | 43.298 | 1.616 | 13.596 | 0.361 | 29.701 | 1.652 | 0.314 | 0.686 | 0.458 | 129 |
| **118** | M9_glucose | 1017.5 | 0.0116 | 0.0009668 | 1619.72 | 0.0016135 | 1017.9 | 0.000458 | 0.0015630 | Q1 hold | 200 | 455 | 655 | 6500 | 43.031 | 1.107 | 29.406 | 0.522 | 13.625 | 1.132 | 41.798 | 2.639 | 13.327 | 0.691 | 28.471 | 2.603 | 0.319 | 0.681 | 0.468 | 132 |
| **119** | M9_glucose | 1017.5 | 0.0116 | 0.0009668 | 1619.72 | 0.0016135 | 1017.9 | 0.000458 | 0.0015630 | Qt | 190 | 460 | 650 | 6500 | 43.62 | 0.546 | 27.935 | 0.64 | 15.685 | 0.937 | 43.478 | 1.633 | 11.432 | 0.777 | 32.046 | 1.971 | 0.263 | 0.737 | 0.357 | 113 |
| **120** | M9_glucose | 1017.5 | 0.0116 | 0.0009668 | 1619.72 | 0.0016135 | 1017.9 | 0.000458 | 0.0015630 | Q1 hold | 200 | 460 | 660 | 6500 | 43.562 | 0.713 | 29.035 | 0.774 | 14.527 | 1.04 | 43.319 | 2.17 | 12.843 | 1.005 | 30.476 | 2.373 | 0.296 | 0.704 | 0.421 | 130 |
| **121** | M9_glucose | 1017.5 | 0.0116 | 0.0009668 | 1619.72 | 0.0016135 | 1017.9 | 0.000458 | 0.0015630 | Q1 hold | 200 | 600 | 800 | 6500 | 45.305 | 0.722 | 28.534 | 0.681 | 16.77 | 0.965 | 48.726 | 2.35 | 12.185 | 0.875 | 36.54 | 2.461 | 0.25 | 0.75 | 0.333 | 117 |
| **122** | M9_glucose | 1017.5 | 0.0116 | 0.0009668 | 1619.72 | 0.0016135 | 1017.9 | 0.000458 | 0.0015630 | Q1 hold | 200 | 700 | 900 | 6500 | 46.05 | 0.86 | 27.849 | 0.638 | 18.201 | 1.106 | 51.184 | 2.87 | 11.326 | 0.771 | 39.857 | 3.022 | 0.221 | 0.779 | 0.284 | 119 |
| **123** | M9_glucose | 1017.5 | 0.0116 | 0.0009668 | 1619.72 | 0.0016135 | 1017.9 | 0.000458 | 0.0015630 | Q1 hold | 200 | 800 | 1000 | 6500 | 45.843 | 0.715 | 26.542 | 0.64 | 19.302 | 0.969 | 50.483 | 2.357 | 9.807 | 0.704 | 40.676 | 2.469 | 0.194 | 0.806 | 0.241 | 120 |
| **124** | M9_glucose | 1017.5 | 0.0116 | 0.0009668 | 1619.72 | 0.0016135 | 1017.9 | 0.000458 | 0.0015630 | Q1 hold | 200 | 900 | 1100 | 6500 | 46.585 | 0.666 | 25.405 | 0.597 | 21.18 | 0.947 | 52.967 | 2.268 | 8.599 | 0.594 | 44.368 | 2.414 | 0.162 | 0.838 | 0.194 | 111 |

| | | | | | | | | | | | | | | | | | | | | | | | | | | | | | | |
|---|---|---|---|---|---|---|---|---|---|---|---|---|---|---|---|---|---|---|---|---|---|---|---|---|---|---|---|---|---|---|
| **125** | M9_glucose | 1017.5 | 0.0116 | 0.009668 | 1619.72 | 0.016135 | 1017.9 | 0.000458 | 0.015630 | Q1 hold | 200 | 100 | 120 | 650 | 47.149 | 0.797 | 24.944 | 0.612 | 22.205 | 1.155 | 54.926 | 2.775 | 8.141 | 0.609 | 46.785 | 3.029 | 0.148 | 0.852 | 0.174 | 113 |
| **126** | M9_glucose | 1017.5 | 0.0116 | 0.009668 | 1619.72 | 0.016135 | 1017.9 | 0.000458 | 0.015630 | Q1 hold | 200 | 120 | 140 | 650 | 47.868 | 0.839 | 24.177 | 0.568 | 23.691 | 1.145 | 57.481 | 3.062 | 7.411 | 0.508 | 50.07 | 3.25 | 0.129 | 0.871 | 0.148 | 89 |
| **127** | PEG_10per | 1013.7 | 0.000461 | 0.003431 | 1619.72 | 0.016135 | 1014.1 | 0.000455 | 0.006395 | Q1 hold | 200 | 350 | 550 | 650 | 38.849 | 0.926 | 27.208 | 0.373 | 11.641 | 0.95 | 30.753 | 2.191 | 10.552 | 0.451 | 20.2 | 2.17 | 0.343 | 0.657 | 0.522 | 142 |
| **128** | PEG_10per | 1013.7 | 0.000461 | 0.003431 | 1619.72 | 0.016135 | 1014.1 | 0.000455 | 0.006395 | Q1 hold | 200 | 370 | 570 | 650 | 39.155 | 0.616 | 27.039 | 0.253 | 12.116 | 0.69 | 31.455 | 1.494 | 10.353 | 0.278 | 21.102 | 1.548 | 0.329 | 0.671 | 0.491 | 126 |
| **129** | PEG_10per | 1013.7 | 0.000461 | 0.003431 | 1619.72 | 0.016135 | 1014.1 | 0.000455 | 0.006395 | Q1 hold | 200 | 390 | 590 | 650 | 39.229 | 0.772 | 25.8 | 0 | 13.429 | 0 | 31.647 | 1.915 | 8.992 | 0 | 22.655 | 0 | 0.284 | 0.716 | 0.397 | 118 |
| **130** | PEG_10per | 1013.7 | 0.000461 | 0.003431 | 1619.72 | 0.016135 | 1014.1 | 0.000455 | 0.006395 | Qt | 210 | 390 | 600 | 650 | 39.253 | 0.757 | 26.984 | 0.355 | 12.269 | 0.92 | 31.702 | 1.863 | 10.293 | 0.39 | 21.409 | 2.004 | 0.325 | 0.675 | 0.481 | 122 |
| **131** | PEG_10per | 1013.7 | 0.000461 | 0.003431 | 1619.72 | 0.016135 | 1014.1 | 0.000455 | 0.006395 | Qt | 205 | 395 | 600 | 650 | 38.618 | 0.578 | 27.052 | 0.22 | 11.566 | 0.662 | 30.176 | 1.344 | 10.367 | 0.242 | 19.809 | 1.418 | 0.344 | 0.656 | 0.523 | 101 |
| **132** | PEG_10per | 1013.7 | 0.000461 | 0.003431 | 1619.72 | 0.016135 | 1014.1 | 0.000455 | 0.006395 | Q2 hold | 150 | 400 | 550 | 650 | 40.273 | 0.851 | 25.626 | 0.442 | 14.647 | 1.07 | 34.247 | 2.171 | 8.819 | 0.44 | 25.428 | 2.341 | 0.258 | 0.742 | 0.347 | 126 |
| **133** | PEG_10per | 1013.7 | 0.000461 | 0.003431 | 1619.72 | 0.016135 | 1014.1 | 0.000455 | 0.006395 | Q2 hold | 160 | 400 | 560 | 650 | 39.52 | 0.604 | 24.65 | 0.402 | 14.87 | 0.759 | 32.341 | 1.482 | 7.849 | 0.4 | 24.492 | 1.574 | 0.243 | 0.757 | 0.32 | 166 |
| **134** | PEG_10per | 1013.7 | 0.000461 | 0.003431 | 1619.72 | 0.016135 | 1014.1 | 0.000455 | 0.006395 | Q2 hold | 180 | 400 | 580 | 650 | 39.696 | 0.812 | 25.792 | 0.101 | 13.904 | 0.828 | 32.792 | 2.011 | 8.984 | 0.101 | 23.808 | 2.023 | 0.274 | 0.726 | 0.377 | 162 |
| **135** | PEG_10per | 1013.7 | 0.000461 | 0.003431 | 1619.72 | 0.016135 | 1014.1 | 0.000455 | 0.006395 | Q2 hold | 190 | 400 | 590 | 650 | 38.973 | 0.614 | 25.8 | 0 | 13.173 | 0 | 31.019 | 1.467 | 8.992 | 0 | 22.027 | 0 | 0.29 | 0.71 | 0.408 | 92 |
| **136** | PEG_10per | 1013.7 | 0.000461 | 0.003431 | 1619 | 0.016135 | 1014.1 | 0.000455 | 0.006395 | Qt | 200 | 400 | 600 | 65 | 39.086 | 0.75 | 27.019 | 0.296 | 12.067 | 0.829 | 31.3 | 1.848 | 10.331 | 0.325 | 20.969 | 1.902 | 0.33 | 0.67 | 0.493 | 127 |

| | | | | | | | | | | | | | | | | | | | | | | | | | | | | | | |
|---|---|---|---|---|---|---|---|---|---|---|---|---|---|---|---|---|---|---|---|---|---|---|---|---|---|---|---|---|---|---|
| | | | | | .72 | | | | | | | | | 00 | | | | | | | | | | | | | | | | |
| **137** | PEG_10per | 1013.7 | 0.000461 | 0.003431 | 1619.72 | 0.016135 | 1014.1 | 0.000455 | 0.006395 | Q2 hold | 220 | 400 | 620 | 6500 | 38.909 | 0.544 | 27.071 | 0.154 | 11.838 | 0.59 | 30.861 | 1.295 | 10.389 | 0.169 | 20.472 | 1.334 | 0.337 | 0.663 | 0.507 | 139 |
| **138** | PEG_10per | 1013.7 | 0.000461 | 0.003431 | 1619.72 | 0.016135 | 1014.1 | 0.000455 | 0.006395 | Qt | 195 | 405 | 600 | 6500 | 39.154 | 0.623 | 25.809 | 0.108 | 13.345 | 0.63 | 31.453 | 1.494 | 9.002 | 0.119 | 22.451 | 1.496 | 0.286 | 0.714 | 0.401 | 142 |
| **139** | PEG_10per | 1013.7 | 0.000461 | 0.003431 | 1619.72 | 0.016135 | 1014.1 | 0.000455 | 0.006395 | Q1 hold | 200 | 405 | 605 | 6500 | 39.223 | 0.949 | 26.937 | 0.419 | 12.286 | 1.069 | 31.651 | 2.371 | 10.241 | 0.46 | 21.41 | 2.449 | 0.324 | 0.676 | 0.478 | 118 |
| **140** | PEG_10per | 1013.7 | 0.000461 | 0.003431 | 1619.72 | 0.016135 | 1014.1 | 0.000455 | 0.006395 | Qt | 190 | 410 | 600 | 6500 | 39.253 | 0.742 | 25.846 | 0.285 | 13.407 | 0.735 | 31.701 | 1.821 | 9.043 | 0.309 | 22.658 | 1.779 | 0.285 | 0.715 | 0.399 | 141 |
| **141** | PEG_10per | 1013.7 | 0.000461 | 0.003431 | 1619.72 | 0.016135 | 1014.1 | 0.000455 | 0.006395 | Q1 hold | 200 | 410 | 610 | 6500 | 38.859 | 0.705 | 26.564 | 0.637 | 12.295 | 1.061 | 30.755 | 1.673 | 9.831 | 0.7 | 20.923 | 1.968 | 0.32 | 0.68 | 0.47 | 103 |
| **142** | PEG_10per | 1013.7 | 0.000461 | 0.003431 | 1619.72 | 0.016135 | 1014.1 | 0.000455 | 0.006395 | Qt | 170 | 430 | 600 | 6500 | 39.727 | 0.731 | 25.694 | 0.355 | 14.033 | 0.89 | 32.861 | 1.848 | 8.887 | 0.353 | 23.975 | 1.968 | 0.27 | 0.73 | 0.371 | 134 |
| **143** | PEG_10per | 1013.7 | 0.000461 | 0.003431 | 1619.72 | 0.016135 | 1014.1 | 0.000455 | 0.006395 | Qt | 150 | 450 | 600 | 6500 | 40.495 | 0.952 | 24.702 | 0.461 | 15.792 | 1.037 | 34.825 | 2.36 | 7.901 | 0.458 | 26.924 | 2.384 | 0.227 | 0.773 | 0.293 | 141 |
| **144** | PEG_10per | 1013.7 | 0.000461 | 0.003431 | 1619.72 | 0.016135 | 1014.1 | 0.000455 | 0.006395 | Q1 hold | 200 | 500 | 700 | 6500 | 39.026 | 0.93 | 25.8 | 0 | 13.226 | 0 | 31.174 | 2.079 | 8.992 | 0 | 22.182 | 0 | 0.288 | 0.712 | 0.405 | 162 |
| **145** | PEG_10per | 1013.7 | 0.000461 | 0.003431 | 1619.72 | 0.016135 | 1014.1 | 0.000455 | 0.006395 | Q1 hold | 200 | 550 | 750 | 6500 | 39.756 | 0.735 | 24.51 | 0 | 15.246 | 0 | 32.935 | 1.848 | 7.71 | 0 | 25.226 | 0 | 0.234 | 0.766 | 0.306 | 120 |
| **146** | PEG_10per | 1013.7 | 0.000461 | 0.003431 | 1619.72 | 0.016135 | 1014.1 | 0.000455 | 0.006395 | Q1 hold | 200 | 600 | 800 | 6500 | 39.866 | 0.725 | 24.499 | 0.119 | 15.367 | 0.736 | 33.207 | 1.832 | 7.7 | 0.107 | 25.507 | 1.836 | 0.232 | 0.768 | 0.302 | 117 |
| **147** | PEG_10per | 1013.7 | 0.000461 | 0.003431 | 1619.72 | 0.016135 | 1014.1 | 0.000455 | 0.006395 | Q1 hold | 200 | 750 | 950 | 6500 | 40.335 | 0.542 | 23.611 | 0.645 | 16.725 | 0.959 | 34.379 | 1.384 | 6.907 | 0.572 | 27.472 | 1.652 | 0.201 | 0.799 | 0.251 | 109 |

| | | | | | | | | | | | | | | | | | | | | | | | | | | | | | | |
|---|---|---|---|---|---|---|---|---|---|---|---|---|---|---|---|---|---|---|---|---|---|---|---|---|---|---|---|---|---|---|
| **148** | PEG_10per | 1013.7 | 0.000461 | 0.003431 | 1619.72 | 0.0016135 | 1014.1 | 0.000455 | 0.006395 | Q1 hold | 200 | 850 | 1050 | 6500 | 40.22 | 0.658 | 23.091 | 0.389 | 17.129 | 0.84 | 34.094 | 1.684 | 6.452 | 0.311 | 27.642 | 1.784 | 0.189 | 0.811 | 0.233 | 120 |
| **149** | PEG_10per | 1013.7 | 0.000461 | 0.003431 | 1619.72 | 0.0016135 | 1014.1 | 0.000455 | 0.006395 | Q1 hold | 200 | 950 | 1150 | 6500 | 40.889 | 0.872 | 21.875 | 0.529 | 19.013 | 1.14 | 35.843 | 2.321 | 5.491 | 0.396 | 30.353 | 2.457 | 0.153 | 0.847 | 0.181 | 142 |
| **150** | PEG_10per | 1013.7 | 0.000461 | 0.003431 | 1619.72 | 0.0016135 | 1014.1 | 0.000455 | 0.006395 | Q1 hold | 200 | 1050 | 1250 | 6500 | 41.069 | 0.726 | 21.363 | 0.643 | 19.706 | 1.107 | 36.302 | 1.946 | 5.118 | 0.458 | 31.184 | 2.133 | 0.141 | 0.859 | 0.164 | 116 |
| **151** | PEG_10per | 1013.7 | 0.000461 | 0.003431 | 1619.72 | 0.0016135 | 1014.1 | 0.000455 | 0.006395 | Q1 hold | 200 | 1200 | 1400 | 6500 | 41.477 | 0.829 | 20.807 | 0.532 | 20.67 | 1.099 | 37.406 | 2.256 | 4.726 | 0.371 | 32.68 | 2.382 | 0.126 | 0.874 | 0.145 | 108 |
| **152** | PEG_10per | 1013.7 | 0.000461 | 0.003431 | 1619.72 | 0.0016135 | 1014.1 | 0.000455 | 0.006395 | Q1 hold | 200 | 395 | 595 | 6500 | 38.984 | 0.756 | 25.968 | 0.469 | 13.016 | 0.773 | 31.056 | 1.832 | 9.178 | 0.511 | 21.878 | 1.757 | 0.296 | 0.704 | 0.42 | 115 |

##### Table S6: Replicate sweeps with fluid property parameters

| condition_num | trial | rho_in | sigma_in_oil | mu_in | rho_oil | mu_oil | rho_out | sigma_out_oil | mu_out | sweep_condition | Q1 | Q2 | Qt | Q3 | outer_diam_mean | outer_diam_std | inner_diam_mean | inner_diam_std | shell_diam_mean | shell_diam_std | outer_vol_mean | outer_vol_std | inner_vol_mean | inner_vol_std | shell_vol_mean | shell_vol_std | core_total_ratio | shell_total_ratio | core_shell_ratio | n_droplets |
|---|---|---|---|---|---|---|---|---|---|---|---|---|---|---|---|---|---|---|---|---|---|---|---|---|---|---|---|---|---|---|
| **1** | PBS_1per_Tween20_replicate | 1006.478 | 0.000320 | 0.009875 | 1619.72 | 0.016135 | 1007.97 | 0.000328 | 0.013034 | Central condition | 230 | 400 | 630 | 6500 | 40.221 | 0.315 | 28.783 | 0.602 | 11.438 | 0.613 | 34.075 | 0.805 | 12.502 | 0.796 | 21.573 | 0.997 | 0.367 | 0.633 | 0.58 | 80 |
| **2** | PBS_1per_Tween20_replicate | 1006.478 | 0.000320 | 0.009875 | 1619.72 | 0.016135 | 1007.97 | 0.000328 | 0.013034 | Q2 hold | 220 | 400 | 620 | 6500 | 40.231 | 0.332 | 28.326 | 0.26 | 11.905 | 0.389 | 34.102 | 0.845 | 11.903 | 0.314 | 22.199 | 0.855 | 0.349 | 0.651 | 0.536 | 72 |
| **3** | PBS_1per_Tween20_replicate | 1006.478 | 0.000320 | 0.009875 | 1619.72 | 0.016135 | 1007.97 | 0.000328 | 0.013034 | Q2 hold | 225 | 400 | 625 | 6500 | 40.162 | 0.361 | 28.372 | 0.34 | 11.79 | 0.471 | 33.928 | 0.922 | 11.963 | 0.43 | 21.964 | 0.979 | 0.353 | 0.647 | 0.545 | 159 |
| **4** | PBS_1per_Tween20_replicate | 1006.478 | 0.000320 | 0.009875 | 1619.72 | 0.016135 | 1007.97 | 0.000328 | 0.013034 | Q2 hold | 235 | 400 | 635 | 6500 | 40.228 | 0.329 | 28.776 | 0.599 | 11.452 | 0.646 | 34.093 | 0.837 | 12.492 | 0.793 | 21.601 | 1.08 | 0.366 | 0.634 | 0.578 | 75 |
| **5** | PBS_1per_Tween20_replicate | 1006.478 | 0.000320 | 0.009875 | 1619.72 | 0.016135 | 1007.97 | 0.000328 | 0.013034 | Q2 hold | 240 | 400 | 640 | 6500 | 40.184 | 0.675 | 28.688 | 0.552 | 11.496 | 0.786 | 34.004 | 1.734 | 12.377 | 0.731 | 21.627 | 1.747 | 0.364 | 0.636 | 0.572 | 138 |
| **6** | PBS_1per_Tween20_replicate | 1006.478 | 0.000320 | 0.009875 | 1619.72 | 0.016135 | 1007.97 | 0.000328 | 0.013034 | Q1 hold | 230 | 390 | 620 | 6500 | 40.238 | 0.351 | 28.711 | 0.571 | 11.527 | 0.697 | 34.12 | 0.892 | 12.406 | 0.755 | 21.714 | 1.221 | 0.364 | 0.636 | 0.571 | 39 |
| **7** | PBS_1per_Tween20_replicate | 1006.478 | 0.000320 | 0.009875 | 1619.72 | 0.016135 | 1007.97 | 0.000328 | 0.013034 | Q1 hold | 230 | 395 | 625 | 6500 | 40.242 | 0.329 | 28.506 | 0.387 | 11.736 | 0.507 | 34.129 | 0.839 | 12.136 | 0.513 | 21.994 | 0.981 | 0.356 | 0.644 | 0.552 | 51 |
| **8** | PBS_1per_Tween20_replicate | 1006.478 | 0.000320 | 0.009875 | 1619.72 | 0.016135 | 1007.97 | 0.000328 | 0.013034 | Q1 hold | 230 | 405 | 635 | 6500 | 40.47 | 0.295 | 29.326 | 0.573 | 11.144 | 0.525 | 34.71 | 0.752 | 13.22 | 0.759 | 21.49 | 0.818 | 0.381 | 0.619 | 0.615 | 105 |
| **9** | PBS_1per_Tween20_replicate | 1006.478 | 0.000320 | 0.009875 | 1619.72 | 0.016135 | 1007.97 | 0.000328 | 0.013034 | Q1 hold | 230 | 410 | 640 | 6500 | 40.106 | 0.338 | 28.321 | 0.419 | 11.785 | 0.549 | 33.785 | 0.856 | 11.902 | 0.522 | 21.883 | 1.021 | 0.352 | 0.648 | 0.544 | 66 |
| **10** | PBS_1per_Tween20_replicate | 1006.478 | 0.000320 | 0.009875 | 1619.72 | 0.016135 | 1007.97 | 0.000328 | 0.013034 | Q2 hold | 150 | 400 | 550 | 6500 | 40.605 | 0.274 | 25.8 | 0 | 14.805 | 0 | 35.06 | 0.709 | 8.992 | 0 | 26.068 | 0 | 0.256 | 0.744 | 0.345 | 160 |

| | | | | | | | | | | | | | | | | | | | | | | | | | | | | | |
|---|---|---|---|---|---|---|---|---|---|---|---|---|---|---|---|---|---|---|---|---|---|---|---|---|---|---|---|---|---|
| **11** | PBS_1per_Tween20_replicate | 1006.478 | 0.000320 | 0.009875 | 1619.72 | 0.016135 | 1007.97 | 0.000328 | 0.013034 | Q2 hold | 190 | 400 | 590 | 6500 | 40.446 | 0.453 | 27.195 | 0.354 | 13.252 | 0.583 | 34.658 | 1.157 | 10.536 | 0.427 | 24.122 | 1.245 | 0.304 | 0.696 | 0.437 | 160 |
| **12** | PBS_1per_Tween20_replicate | 1006.478 | 0.000320 | 0.009875 | 1619.72 | 0.016135 | 1007.97 | 0.000328 | 0.013034 | Q2 hold | 270 | 400 | 670 | 6500 | 40.887 | 1.339 | 30.524 | 0.726 | 10.363 | 1.266 | 35.902 | 3.49 | 14.915 | 1.066 | 20.986 | 3.267 | 0.415 | 0.585 | 0.711 | 68 |
| **13** | PBS_1per_Tween20_replicate | 1006.478 | 0.000320 | 0.009875 | 1619.72 | 0.016135 | 1007.97 | 0.000328 | 0.013034 | Q1 hold | 230 | 280 | 510 | 6500 | 40.265 | 0.948 | 30.751 | 0.961 | 9.514 | 0.583 | 34.237 | 2.42 | 15.27 | 1.441 | 18.967 | 1.529 | 0.446 | 0.554 | 0.805 | 68 |
| **14** | PBS_1per_Tween20_replicate | 1006.478 | 0.000320 | 0.009875 | 1619.72 | 0.016135 | 1007.97 | 0.000328 | 0.013034 | Q1 hold | 230 | 520 | 750 | 6500 | 40 | 0.348 | 26.364 | 0.886 | 13.635 | 1.124 | 33.517 | 0.886 | 9.627 | 0.943 | 23.89 | 1.618 | 0.287 | 0.713 | 0.403 | 64 |
| **15** | PBS_1per_Tween20_replicate | 1006.478 | 0.000320 | 0.009875 | 1619.72 | 0.016135 | 1007.97 | 0.000328 | 0.013034 | Q1 hold | 230 | 640 | 870 | 6500 | 40.015 | 0.285 | 25.532 | 0.527 | 14.483 | 0.624 | 33.553 | 0.719 | 8.726 | 0.524 | 24.828 | 0.93 | 0.26 | 0.74 | 0.351 | 77 |
| **16** | PBS_1per_Tween20_replicate | 1006.478 | 0.000320 | 0.009875 | 1619.72 | 0.016135 | 1007.97 | 0.000328 | 0.013034 | Q1 hold | 230 | 760 | 990 | 6500 | 39.784 | 0.329 | 24.501 | 0.106 | 15.283 | 0.334 | 32.977 | 0.812 | 7.702 | 0.095 | 25.275 | 0.807 | 0.234 | 0.766 | 0.305 | 147 |
| **17** | PBS_1per_Tween20_replicate | 1006.478 | 0.000320 | 0.009875 | 1619.72 | 0.016135 | 1007.97 | 0.000328 | 0.013034 | Qt | 240 | 390 | 630 | 6500 | 40.482 | 0.649 | 29.167 | 0.632 | 11.315 | 0.885 | 34.764 | 1.632 | 13.01 | 0.836 | 21.753 | 1.796 | 0.374 | 0.626 | 0.598 | 118 |
| **18** | PBS_1per_Tween20_replicate | 1006.478 | 0.000320 | 0.009875 | 1619.72 | 0.016135 | 1007.97 | 0.000328 | 0.013034 | Qt | 235 | 395 | 630 | 6500 | 40.568 | 0.398 | 28.698 | 0.56 | 11.87 | 0.56 | 34.968 | 1.033 | 12.389 | 0.741 | 22.579 | 1.035 | 0.354 | 0.646 | 0.549 | 65 |
| **19** | PBS_1per_Tween20_replicate | 1006.478 | 0.000320 | 0.009875 | 1619.72 | 0.016135 | 1007.97 | 0.000328 | 0.013034 | Qt | 225 | 405 | 630 | 6500 | 40.444 | 0.441 | 28.434 | 0.26 | 12.01 | 0.476 | 34.652 | 1.105 | 12.04 | 0.344 | 22.612 | 1.105 | 0.347 | 0.653 | 0.532 | 119 |
| **20** | PBS_1per_Tween20_replicate | 1006.478 | 0.000320 | 0.009875 | 1619.72 | 0.016135 | 1007.97 | 0.000328 | 0.013034 | Qt | 220 | 410 | 630 | 6500 | 40.501 | 0.315 | 28.366 | 0.4 | 12.135 | 0.489 | 34.792 | 0.808 | 11.958 | 0.505 | 22.834 | 0.916 | 0.344 | 0.656 | 0.524 | 188 |
| **21** | PBS_1per_Tween20_replicate | 1006.478 | 0.000320 | 0.009875 | 1619.72 | 0.016135 | 1007.97 | 0.000328 | 0.013034 | Qt | 150 | 480 | 630 | 6500 | 41.515 | 0.325 | 24.848 | 0.57 | 16.668 | 0.669 | 37.472 | 0.882 | 8.045 | 0.566 | 29.427 | 1.071 | 0.215 | 0.785 | 0.273 | 107 |
| **22** | PBS_1per_Tween20_replicate | 1006. | 0.000320 | 0.009875 | 1619 | 0.016135 | 1007 | 0.000328 | 0.013034 | Qt | 110 | 520 | 630 | 65 | 42.537 | 0.252 | 23.22 | 0 | 19.317 | 0 | 40.304 | 0.714 | 6.555 | 0 | 33.749 | 0 | 0.163 | 0.837 | 0.194 | 59 |

| | | | | | | | | | | | | | | | | | | | | | | | | | | | | | | |
|---|---|---|---|---|---|---|---|---|---|---|---|---|---|---|---|---|---|---|---|---|---|---|---|---|---|---|---|---|---|---|
| | | 478 | | | .72 | | .97 | | | | | | | 00 | | | | | | | | | | | | | | | | |
| **23** | PBS_1per_Tween20_replicate | 1006.478 | 0.000320 | 0.009875 | 1619.72 | 0.016135 | 1007.97 | 0.000328 | 0.013034 | Qt | 190 | 440 | 630 | 6500 | 40.732 | 0.294 | 27.09 | 0 | 13.642 | 0 | 35.39 | 0.769 | 10.409 | 0 | 24.981 | 0 | 0.294 | 0.706 | 0.417 | 53 |
| **24** | NP40_inner_replicate | 1006.07 | 0.00141 | 0.010028 | 1619.72 | 0.016135 | 1007.97 | 0.000318 | 0.013034 | Q1 hold | 230 | 390 | 620 | 6500 | 39.947 | 1.05 | 26.796 | 0.544 | 13.151 | 1.307 | 33.444 | 2.603 | 10.086 | 0.598 | 23.359 | 2.822 | 0.302 | 0.698 | 0.432 | 92 |
| **25** | NP40_inner_replicate | 1006.07 | 0.00141 | 0.010028 | 1619.72 | 0.016135 | 1007.97 | 0.000318 | 0.013034 | Qt | 240 | 390 | 630 | 6500 | 38.399 | 0.679 | 26.906 | 0.454 | 11.493 | 0.897 | 29.673 | 1.585 | 10.207 | 0.499 | 19.466 | 1.763 | 0.344 | 0.656 | 0.524 | 98 |
| **26** | NP40_inner_replicate | 1006.07 | 0.00141 | 0.010028 | 1619.72 | 0.016135 | 1007.97 | 0.000318 | 0.013034 | Q1 hold | 230 | 395 | 625 | 6500 | 39.43 | 1.228 | 27.039 | 0.253 | 12.391 | 1.296 | 32.19 | 2.998 | 10.353 | 0.278 | 21.837 | 3.059 | 0.322 | 0.678 | 0.474 | 101 |
| **27** | NP40_inner_replicate | 1006.07 | 0.00141 | 0.010028 | 1619.72 | 0.016135 | 1007.97 | 0.000318 | 0.013034 | Qt | 235 | 395 | 630 | 6500 | 38.502 | 0.543 | 26.189 | 0.596 | 12.313 | 0.726 | 29.903 | 1.285 | 9.419 | 0.655 | 20.484 | 1.322 | 0.315 | 0.685 | 0.46 | 73 |
| **28** | NP40_inner_replicate | 1006.07 | 0.00141 | 0.010028 | 1619.72 | 0.016135 | 1007.97 | 0.000318 | 0.013034 | Q2 hold | 110 | 400 | 510 | 6500 | 41.252 | 0.457 | 23.22 | 0 | 18.032 | 0 | 36.769 | 1.217 | 6.555 | 0 | 30.214 | 0 | 0.178 | 0.822 | 0.217 | 114 |
| **29** | NP40_inner_replicate | 1006.07 | 0.00141 | 0.010028 | 1619.72 | 0.016135 | 1007.97 | 0.000318 | 0.013034 | Q2 hold | 150 | 400 | 550 | 6500 | 39.793 | 0.417 | 24.51 | 0 | 15.283 | 0 | 33.004 | 1.037 | 7.71 | 0 | 25.295 | 0 | 0.234 | 0.766 | 0.305 | 118 |
| **30** | NP40_inner_replicate | 1006.07 | 0.00141 | 0.010028 | 1619.72 | 0.016135 | 1007.97 | 0.000318 | 0.013034 | Q2 hold | 220 | 400 | 620 | 6500 | 39.628 | 1.122 | 26.885 | 0.474 | 12.743 | 1.338 | 32.661 | 2.749 | 10.184 | 0.521 | 22.477 | 2.942 | 0.312 | 0.688 | 0.453 | 107 |
| **31** | NP40_inner_replicate | 1006.07 | 0.00141 | 0.010028 | 1619.72 | 0.016135 | 1007.97 | 0.000318 | 0.013034 | Q2 hold | 225 | 400 | 625 | 6500 | 39.453 | 1.439 | 27.013 | 0.401 | 12.44 | 1.575 | 32.282 | 3.55 | 10.328 | 0.45 | 21.954 | 3.672 | 0.32 | 0.68 | 0.47 | 101 |
| **32** | NP40_inner_replicate | 1006.07 | 0.00141 | 0.010028 | 1619.72 | 0.016135 | 1007.97 | 0.000318 | 0.013034 | Q2 hold | 230 | 400 | 630 | 6500 | 39.866 | 1.236 | 27.09 | 0.548 | 12.776 | 1.458 | 33.269 | 3.057 | 10.422 | 0.633 | 22.847 | 3.256 | 0.313 | 0.687 | 0.456 | 123 |
| **33** | NP40_inner_replicate | 1006.07 | 0.00141 | 0.010028 | 1619.72 | 0.016135 | 1007.97 | 0.000318 | 0.013034 | Q2 hold | 235 | 400 | 635 | 6500 | 39.187 | 1.392 | 27.172 | 0.415 | 12.015 | 1.566 | 31.627 | 3.396 | 10.512 | 0.492 | 21.115 | 3.573 | 0.332 | 0.668 | 0.498 | 94 |

| | | | | | | | | | | | | | | | | | | | | | | | | | | | | | | |
|---|---|---|---|---|---|---|---|---|---|---|---|---|---|---|---|---|---|---|---|---|---|---|---|---|---|---|---|---|---|---|
| **34** | NP40_inner_replicate | 1006.07 | 0.00141 | 0.0010028 | 1619.72 | 0.0016135 | 1007.97 | 0.000318 | 0.0013034 | Q2 hold | 240 | 400 | 640 | 6500 | 39.691 | 1.761 | 27.423 | 0.568 | 12.268 | 1.938 | 32.93 | 4.351 | 10.812 | 0.686 | 22.119 | 4.51 | 0.328 | 0.672 | 0.489 | 93 |
| **35** | NP40_inner_replicate | 1006.07 | 0.00141 | 0.0010028 | 1619.72 | 0.0016135 | 1007.97 | 0.000318 | 0.0013034 | Qt | 225 | 405 | 630 | 6500 | 38.194 | 0.353 | 25.837 | 0.28 | 12.357 | 0.479 | 29.18 | 0.81 | 9.034 | 0.302 | 20.147 | 0.901 | 0.31 | 0.69 | 0.448 | 105 |
| **36** | NP40_inner_replicate | 1006.07 | 0.00141 | 0.0010028 | 1619.72 | 0.0016135 | 1007.97 | 0.000318 | 0.0013034 | Q1 hold | 230 | 405 | 635 | 6500 | 39.358 | 1.172 | 27.045 | 0.237 | 12.312 | 1.232 | 32.006 | 2.849 | 10.36 | 0.261 | 21.646 | 2.9 | 0.324 | 0.676 | 0.479 | 115 |
| **37** | NP40_inner_replicate | 1006.07 | 0.00141 | 0.0010028 | 1619.72 | 0.0016135 | 1007.97 | 0.000318 | 0.0013034 | Qt | 220 | 410 | 630 | 6500 | 38.257 | 0.41 | 25.618 | 0.452 | 12.639 | 0.627 | 29.328 | 0.943 | 8.811 | 0.449 | 20.517 | 1.068 | 0.3 | 0.7 | 0.429 | 92 |
| **38** | NP40_inner_replicate | 1006.07 | 0.00141 | 0.0010028 | 1619.72 | 0.0016135 | 1007.97 | 0.000318 | 0.0013034 | Q1 hold | 230 | 410 | 640 | 6500 | 39.597 | 1.016 | 26.721 | 0.586 | 12.876 | 1.213 | 32.571 | 2.493 | 10.004 | 0.644 | 22.567 | 2.62 | 0.307 | 0.693 | 0.443 | 91 |
| **39** | NP40_inner_replicate | 1006.07 | 0.00141 | 0.0010028 | 1619.72 | 0.0016135 | 1007.97 | 0.000318 | 0.0013034 | Qt | 190 | 440 | 630 | 6500 | 38.098 | 0.372 | 24.496 | 0.136 | 13.602 | 0.417 | 28.962 | 0.848 | 7.697 | 0.122 | 21.265 | 0.878 | 0.266 | 0.734 | 0.362 | 90 |
| **40** | NP40_inner_replicate | 1006.07 | 0.00141 | 0.0010028 | 1619.72 | 0.0016135 | 1007.97 | 0.000318 | 0.0013034 | Qt | 150 | 480 | 630 | 6500 | 39.133 | 0.395 | 22.565 | 0.65 | 16.568 | 0.842 | 31.388 | 0.954 | 6.031 | 0.52 | 25.357 | 1.197 | 0.192 | 0.808 | 0.238 | 67 |
| **41** | NP40_inner_replicate | 1006.07 | 0.00141 | 0.0010028 | 1619.72 | 0.0016135 | 1007.97 | 0.000318 | 0.0013034 | Qt | 110 | 520 | 630 | 6500 | 42.833 | 0.532 | 21.059 | 0.607 | 21.774 | 0.762 | 41.167 | 1.531 | 4.902 | 0.432 | 36.264 | 1.544 | 0.119 | 0.881 | 0.135 | 120 |
| **42** | NP40_inner_replicate | 1006.07 | 0.00141 | 0.0010028 | 1619.72 | 0.0016135 | 1007.97 | 0.000318 | 0.0013034 | Q1 hold | 230 | 520 | 750 | 6500 | 39.649 | 0.594 | 26.154 | 0.578 | 13.496 | 0.818 | 32.659 | 1.459 | 9.381 | 0.635 | 23.278 | 1.576 | 0.287 | 0.713 | 0.403 | 124 |
| **43** | NP40_inner_replicate | 1006.07 | 0.00141 | 0.0010028 | 1619.72 | 0.0016135 | 1007.97 | 0.000318 | 0.0013034 | Q1 hold | 230 | 640 | 870 | 6500 | 39.878 | 0.445 | 24.51 | 0 | 15.368 | 0 | 33.216 | 1.114 | 7.71 | 0 | 25.506 | 0 | 0.232 | 0.768 | 0.302 | 86 |
| **44** | NP40_inner_replicate | 1006.07 | 0.00141 | 0.0010028 | 1619.72 | 0.0016135 | 1007.97 | 0.000318 | 0.0013034 | Q1 hold | 230 | 760 | 990 | 6500 | 40.515 | 0.71 | 23.265 | 0.31 | 17.25 | 0.578 | 34.854 | 1.947 | 6.597 | 0.289 | 28.257 | 1.773 | 0.189 | 0.811 | 0.233 | 86 |

#### Table S7: Instability conditions

| condition_num | trial | sweep_condition | Q1 | Q2 | Qt | Q3 | flags | outer_diam_mean | outer_diam_std | inner_diam_mean | inner_diam_std | shell_diam_mean | shell_diam_std | outer_vol_mean | outer_vol_std | inner_vol_mean | inner_vol_std | shell_vol_mean | shell_vol_std | core_total_ratio | shell_total_ratio | core_shell_ratio | n_droplets |
|---|---|---|---|---|---|---|---|---|---|---|---|---|---|---|---|---|---|---|---|---|---|---|---|
| 1 | PBS_1per_Tween20 | Qt | 240 | 390 | 630 | 6500 | instability case 1 | 39.47 | 2.095 | 27.765 | 0.648 | 11.705 | 2.416 | 32.463 | 5.047 | 11.225 | 0.783 | 21.238 | 5.387 | 0.346 | 0.654 | 0.529 | 86 |
| 2 | PBS_1per_Tween20 | Qt | 235 | 395 | 630 | 6500 | instability case 1 | 40.173 | 1.65 | 27.66 | 0.674 | 12.513 | 1.987 | 34.114 | 4.08 | 11.1 | 0.81 | 23.015 | 4.425 | 0.325 | 0.675 | 0.482 | 86 |
| 3 | PBS_1per_Tween20 | Q2 hold | 235 | 400 | 635 | 6500 | instability case 1 | 39.262 | 1.802 | 27.524 | 0.729 | 11.738 | 2.171 | 31.886 | 4.331 | 10.941 | 0.866 | 20.945 | 4.705 | 0.343 | 0.657 | 0.522 | 107 |
| 4 | PBS_1per_Tween20 | Q2 hold | 240 | 400 | 640 | 6500 | instability case 1 | 39.474 | 1.936 | 27.728 | 0.724 | 11.746 | 2.273 | 32.436 | 4.732 | 11.185 | 0.88 | 21.25 | 5.073 | 0.345 | 0.655 | 0.526 | 97 |
| 5 | PBS_1per_Tween20_replicate | Q2 hold | 110 | 400 | 510 | 6500 | pull case 2 | 39.641 | 2.199 | 24.412 | 0.344 | 15.229 | 2.247 | 32.908 | 5.177 | 7.622 | 0.308 | 25.286 | 5.206 | 0.232 | 0.768 | 0.301 | 66 |
| 7 | NP40_inner | Q1 hold | 240 | 220 | 460 | 6500 | pull case 1 | 38.979 | 1.03 | 30.686 | 0.862 | 8.293 | 1.637 | 31.074 | 2.478 | 15.165 | 1.243 | 15.91 | 3.287 | 0.488 | 0.512 | 0.953 | 66 |
| 8 | NP40_inner | Q1 hold | 240 | 320 | 560 | 6500 | instability case 1 | 40.101 | 1.423 | 29.007 | 0.684 | 11.094 | 1.475 | 33.889 | 3.534 | 12.8 | 0.9 | 21.089 | 3.501 | 0.378 | 0.622 | 0.607 | 72 |
| 9 | NP40_inner | Q1 hold | 240 | 350 | 590 | 6500 | pull case 1 | 38.807 | 1.721 | 27.041 | 0.561 | 11.766 | 1.949 | 30.778 | 3.997 | 10.366 | 0.643 | 20.411 | 4.215 | 0.337 | 0.663 | 0.508 | 132 |
| 10 | NP40_inner | Qt | 300 | 440 | 740 | 6500 | instability case 1 | 38.624 | 1.503 | 27.232 | 0.497 | 11.391 | 1.66 | 30.305 | 3.518 | 10.585 | 0.59 | 19.72 | 3.664 | 0.349 | 0.651 | 0.537 | 163 |

| | | | | | | | | | | | | | | | | | | | | | |
|---|---|---|---|---|---|---|---|---|---|---|---|---|---|---|---|---|---|---|---|---|---|
| **11** | M9 | Qt | 310 | 390 | 700 | 6500 | instability case 1 | 41.933 | 1.438 | 30.96 | 0 | 10.973 | 0 | 38.742 | 3.981 | 15.538 | 0 | 23.204 | 0 | 0.401 | 0.599 | 0.67 | 124 |
| **12** | M9 | Q2 hold | 170 | 450 | 620 | 6500 | pull case 2 | 39.853 | 1.379 | 28.103 | 0.532 | 11.75 | 1.431 | 33.253 | 2.925 | 11.633 | 0.643 | 21.62 | 2.916 | 0.35 | 0.65 | 0.538 | 121 |
| **13** | M9 | Qt | 170 | 530 | 700 | 6500 | pull case 2 | 40.87 | 0.676 | 26.328 | 0.762 | 14.542 | 1.103 | 35.772 | 1.775 | 9.578 | 0.822 | 26.194 | 2.081 | 0.268 | 0.732 | 0.366 | 22 |
| **14** | PEG_10per | Q2 hold | 100 | 400 | 500 | 6500 | pull case 2 | 39.311 | 0.926 | 23.2 | 0.16 | 16.111 | 1.02 | 31.861 | 2.333 | 6.539 | 0.128 | 25.322 | 2.407 | 0.205 | 0.795 | 0.258 | 65 |
| **15** | PEG_10per | Q2 hold | 230 | 400 | 630 | 6500 | pull case 2 | 37.553 | 1.497 | 28.144 | 0.501 | 9.409 | 1.746 | 27.862 | 3.48 | 11.684 | 0.605 | 16.178 | 3.759 | 0.419 | 0.581 | 0.722 | 104 |
| **16** | NP40_inner_replicate | Q1 hold | 230 | 280 | 510 | 6500 | instability case 2 | 40.275 | 1.411 | 29.553 | 0.373 | 10.722 | 1.473 | 34.331 | 3.617 | 13.521 | 0.493 | 20.81 | 3.666 | 0.394 | 0.606 | 0.65 | 99 |
| **17** | NP40_inner_replicate | Qt | 310 | 320 | 630 | 6500 | pull case 1 | 39.548 | 2.035 | 29.822 | 0.528 | 9.726 | 2.183 | 32.635 | 4.974 | 13.899 | 0.75 | 18.736 | 5.145 | 0.426 | 0.574 | 0.742 | 34 |
| **18** | NP40_inner_replicate | Q2 hold | 70 | 400 | 470 | 6500 | pull case 2 | 39.068 | 0.44 | 21.93 | 0 | 17.138 | 0 | 31.233 | 1.047 | 5.522 | 0 | 25.711 | 0 | 0.177 | 0.823 | 0.215 | 74 |
| **19** | NP40_inner_replicate | Qt | 270 | 360 | 630 | 6500 | instability case 1 | 39.722 | 1.817 | 28.341 | 0.624 | 11.382 | 2.137 | 33.02 | 4.448 | 11.936 | 0.787 | 21.084 | 4.81 | 0.361 | 0.639 | 0.566 | 99 |
